# Enzyme-Constrained Metabolic Modeling Reveals CO_2_-Dependent Growth Control in *Picochlorum celeri*

**DOI:** 10.64898/2026.10.30.763171

**Authors:** Niaz Bahar Chowdhury, Yasin Torres-Tiji, Angela Cintolesi, Matthew C Posewitz

## Abstract

*Picochlorum celeri* grows 11.6 times faster at elevated CO_2_ than in air, yet the abundance of its metabolic enzymes is only 17% higher. To explain this, we reconstructed iPiccel954, a genome-scale metabolic model with 3525 reactions, 2973 metabolites and 954 genes, and eciPiccel954, its enzyme-constrained version limited by the measured enzyme abundance in air and at elevated CO_2_. Calibrating the enzyme-constrained models against the measured proteome showed that a similar protein budget supports very different growth depending on how it is allocated. With urea as the nitrogen source, urea carboxylase was the only step whose relaxation alone raised growth, by 627% in air and 78% at elevated CO_2_. Bottleneck analysis showed that growth is limited by many enzymes spread across the network in both conditions, that the limiting enzymes differ between the two, and that control over starch metabolism seen in air disappears at elevated CO_2_. Measured proteomes agreed with these predictions, since the predicted bottleneck enzymes were more abundant than other enzymes. Rapid growth in *P. celeri* therefore reflects how the protein budget is allocated more than how large it is. Therefore, eciPiccel954 provides a quantitative basis for engineering this alga for biotechnological application.

## INTRODUCTION

Microalgae convert light and CO_2_ into biomass and high-value molecules, which makes them attractive for biotechnology (Dupont-Thibert et al., 2026), and species that grow rapidly are especially valuable for this purpose (Torres-Tiji et al., 2020). The marine chlorophyte *Picochlorum celeri* is one such species. Its small unicellular cells carry a compact genome of 13.7 Mbp per haploid copy, about ten times smaller than that of *Chlamydomonas reinhardtii* (Becker et al., 2020; Merchant et al., 2007), and comparisons across the *Picochlorum* lineage point to streamlining through lost pathways and fewer gene copies (Bec et al., 2025). With supplemental CO_2_, cultures of *P. celeri* double every 2-3 hours and have averaged more than 30 gm^−2^day^−1^ during summer outdoor cultivation (Krishnan et al., 2021; Weissman et al., 2018), but without it productivity falls by more than tenfold (Torres-Tiji et al., 2026).

The reason for this dependence from systems level is still unclear. RuBisCO binds CO_2_ weakly and also reacts with oxygen, so photosynthesis suffers when CO_2_ is scarce (Flamholz et al., 2019), and the carbon-concentrating mechanism of *P. celeri* appears too rudimentary to compensate (Ekness et al., 2026). Therefore, how carbon limitation spreads to the rest of metabolism are not known. A recent multiomics study compared *P. celeri* grown in ambient and elevated CO_2_ and found that cells in air invested less in cytosolic translation and in external nitrogen acquisition, while accumulating more carbon fixation and photorespiratory enzymes (Torres-Tiji et al., 2026). Such data reveal which proteins change but measuring a protein does not tell us about the flux through its reaction (Canelas et al., 2010), and abundance alone cannot show whether the capacity of individual enzymes limits growth. It also cannot show whether that limit is shared by many enzymes or concentrated in a few, which holds a significant importance in the biotechnological application. A limit shared by many enzymes cannot be relieved by modifying a single gene, whereas a limit concentrated in one enzyme identifies a clear engineering target.

Genome-scale metabolic models (GEMs) can address this concern as they link genes to reactions and predict growth and fluxes by assuming that cells operate at steady state and optimize an objective such as growth (Nikoloski et al., 2015). Such models now exist for several green microalgae, including *Chlamydomonas reinhardtii*, *Chlorella vulgaris*, *Chlorella ohadii*, *Scenedesmus obliquus* and *Microchloropsis gaditana*, and they have mostly been used to compare cultivation conditions and to identify targets for strain improvement (Dupont-Thibert et al., 2026; Ray et al., 2023; Soleymani et al., 2025). Why growth itself differs between conditions of the same organism, and whether enzyme capacity limits it, has been addressed less often. Predictions improve when reaction fluxes are limited by enzyme turnover numbers (k_cat_) and by the total protein content of the cell, which gives better estimates of growth, fluxes and enzyme allocation (Arend et al., 2023; Ferreira et al., 2024). The enzyme-constrained genome-scale metabolic models (ecGEMs), which tie each flux to the abundance and k_cat_ of its enzyme, is widely used for this purpose (Chowdhury et al., 2026; Sanchez et al., 2017; Shahreen et al., 2024; Shahreen et al., 2025). It was recently applied to the very fast-growing alga *Chlorella ohadii*, which doubles in about 1.4 hours to compare species and to identify targets for improving growth (Soleymani et al., 2025). ecGEMs have thus been used to predict growth and to compare species, but they have rarely been used to explain how a large change in growth within one organism arises from a nearly constant protein budget. *P. celeri* offers a case to test this, because its growth changes more than ten-fold with CO_2_ supply while the amount of enzyme measured in the model changes little.

This is the central puzzle of this study. Growth of *P. celeri* increases 11.6-fold from air to elevated CO_2_, yet the summed measured abundance of the enzymes represented in the model rises by only 17%. If growth were set by the size of the enzyme pool, the two would scale together. We therefore asked how the similar protein budget can support such different growth, which enzymes limit growth in each condition, and how nitrogen assimilation from urea, the sole added nitrogen source in the cultivation medium (Torres-Tiji et al., 2026), is coupled to carbon metabolism.

To answer these questions, we reconstructed iPiccel954, a GEM of *P. celeri* with 3525 reactions, 2973 metabolites and 954 genes, and reconstructed its ecGEM, eciPiccel954, using measured enzyme abundances in air and at elevated CO_2_. We calibrated the enzyme k_cat_ with an iterative procedure that first relaxes the bounds of reactions whose flux exceeds their k_cat_-derived limit and then raises the k_cat_ of the reaction consuming the largest share of the protein pool until the protein demand fits the measured pool. We then quantified the control that each enzyme exerts on growth with k_cat_ control coefficients and used a nitrogen-carbon coupling analysis to test which steps of urea assimilation limit growth, before comparing the predicted limiting enzymes with the measured proteomes. Our results show that growth is capped by many enzymes spread across the network in both conditions, that the limiting enzymes change with CO_2_ supply, and that rapid growth reflects how the protein budget is allocated more than how large it is. These results point to where engineering effort on *P. celeri* is most likely to pay off.

## RESULTS

### Reconstruction and curation of the iPiccel954

We reconstructed the *P. celeri* GEM in two phases, an assembly phase and a curation phase (Fig. 1A). The assembly phase combined a draft built with Rapid Annotation of Photosynthetic Systems (RAPS) (Metcalf et al., 2020) from iCre1355 of *Chlamydomonas reinhardtii* (Imam et al., 2015) and a homology comparison with iPicre1230 (Tec-Campos et al., 2026), a GEM for *Picochlorum renovo*, and it produced an uncurated model, iPiccel1084, with 4344 reactions, 3235 metabolites and 1084 genes in 9 compartments. We then curated this draft GEM in stages that addressed biomass, mass balance, annotation and thermodynamically infeasible cycles (TICs).

**Fig. 1.**
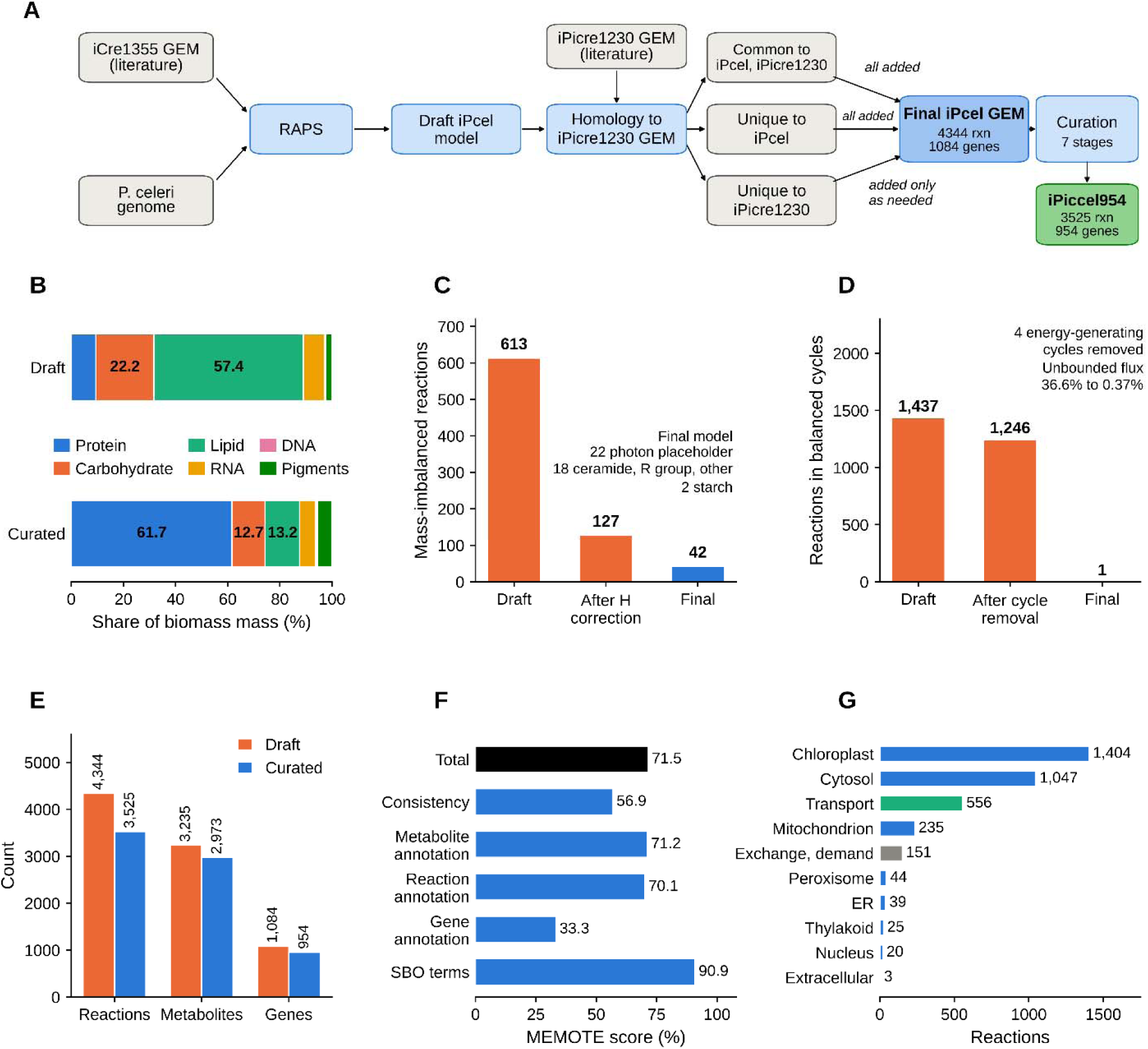
Reconstruction and curation of the iPiccel954. (A) Reconstruction pipeline from the draft to the curated model. (B) Biomass composition before and after curation. (C) Mass-imbalanced reactions at each stage. (D) Reactions in stoichiometrically balanced cycles at each stage. (E) Model size before and after curation. (F) MEMOTE scores of the final model. (G) Reactions by compartment and type.

The draft biomass reaction summed to 1643.8 gmmol^−1^, and its composition did not resemble the measured composition of *P. celeri*, with protein at only 9.5% and lipid at 57.4% of the non-maintenance mass. We rebuilt it from published measurements (LaPanse et al., 2024), which brought protein to the dominant fraction and the total to 1000 gmol^−1^ (Fig. 1B, Supplementary Fig. S1A). We left the growth-associated maintenance requirement of 39.24 mmol ATPgDW^−1^ unchanged from iCre1355. The arginine coefficient in the biomass protein pool is lower than the protein composition would suggest, and we treat it as a limitation of the current biomass definition.

Of the 4344 reactions, 613 were elementally imbalanced, and 486 of these differed only in hydrogen (Fig. 1C). We corrected 485 of the hydrogen imbalances and 36 metabolite formulas. The remaining hydrogen-only reaction, NDP7er, supplies a proton that the network depends on, and we kept it unchanged. 42 imbalances remain. 22 involve the photon placeholder element, 12 involve ceramide and fatty acid species, 3 involve generic R groups, 2 involve starch degradation, 2 involve the hydrogen content of electron carriers and 1 is NDP7er (Supplementary Fig. S1B). The photon, R-group and starch species cannot be given a fixed formula. We also transferred cross-database annotations to 52.6% of metabolites and 46.7% of reactions and assigned Systems Biology Ontology (SBO) terms to every reaction, metabolite and gene.

The draft also contained four energy-generating cycles that produced NADPH, NADH, FADH2 or the glutamate and 2-oxoglutarate couple without any input (Fritzemeier et al., 2017). We removed them without altering any gene-reaction rule, and we restricted the remaining directions that allowed unbounded flux. The share of reactions with unbounded flux fell from 36.6% to 0.37%, and the number of balanced cycles fell from 1437 to 1246 after mass balancing and to 1 in the final model (Fig. 1D).

The final model, iPiccel954, contains 3525 reactions, 2973 metabolites and 954 genes (Fig. 1E). Its MEMOTE total score is 71.5% (Lieven et al., 2020). The consistency score of 56.9% reflects the photon placeholder, which prevents the stoichiometric consistency test from passing (Fig. 1F). Of the 3525 reactions, 1238 have a gene association, 556 are transport reactions and 640 allow negative flux, and they are distributed across nine compartments (Fig. 1G, Supplementary Fig. S1C). Fatty acid reactions account for 1700 of the total, and only 6% of them have a gene association, which lowers the model-wide share to 35% (Supplementary Fig. S1D). The model therefore has 3.7 reactions per gene, or 2.0 once the fatty acid reactions are excluded (Supplementary Fig. S1E). The most connected metabolites are shown in Supplementary Fig. S1F.

### Structural comparison with other algal models and validation with experimental data

We compared iPiccel954 with four published green algal GEMs, namely iCre1355, iCO1515 of *Chlorella ohadii* (Soleymani et al., 2025), iCZ843 of *Chlorella vulgaris* (Zuniga et al., 2016) and iAJ526 of *Chlorella variabilis* (Juneja et al., 2016). With 954 genes, iPiccel954 is smaller than iCO1515 and iCre1355, which have 1515 and 1460 genes, and larger than iCZ843 and iAJ526, which have 843 and 531 genes (Fig. 2A). Its 3525 reactions are more than in any of the other models, which range from 1455 to 2516 (Fig. 2B). Its 2973 metabolites are second only to the 3517 of iCO1515 (Fig. 2C), and its nine compartments fall between the eight of iCO1515 and the eleven of iCre1355 (Fig. 2D). The network is therefore large in reactions relative to its gene content.

**Fig. 2.**
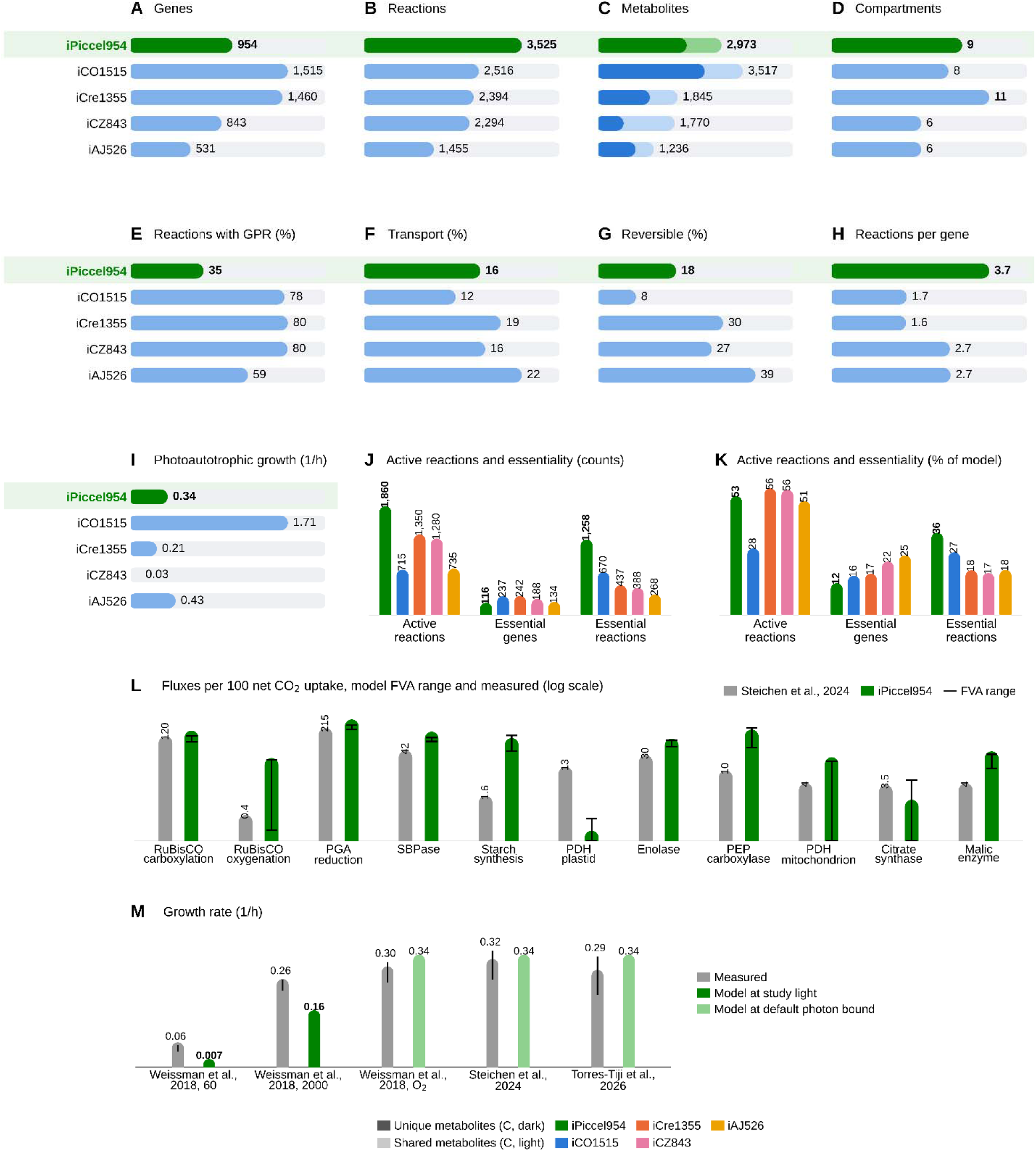
Structural comparison and experimental validation of iPiccel954. (A) Number of genes. (B) Number of reactions. (C) Number of metabolites, with distinct chemical species in the dark segment and compartment copies in the light segment. (D) Number of compartments. (E) Reactions with a gene association. (F) Transport reactions. (G) Reactions that allow negative flux. (H) Reactions per gene. (I) Photoautotrophic growth rate. (J) Active reactions, essential genes and essential reactions. (K) Active reactions, essential genes and essential reactions as a percentage of the model. (L) Model flux range from flux variability analysis and measured fluxes from Steichen et al., 2024, per 100 net CO_2_ uptake. (M) Measured growth rates from literature and model growth rates at the matching light input where it is known.

This difference is most visible in gene association. Only 35% of the reactions in iPiccel954 have a gene association, against 59% to 80% in the published models (Fig. 2E). The gap comes from the fatty acid reactions, which account for 1700 of the 3,525 reactions, and only 6% of them have a gene association (Supplementary Fig. S2A). These reactions describe individual acyl-chain species and cannot be tied to single genes. The other pathways are much better supported, for example with 96% of glycerolipid reactions and 77% of cofactor biosynthesis reactions associated with a gene (Supplementary Fig. S2A). Transport reactions make up 16% of iPiccel954, within the range of 12% to 22% of the published models (Fig. 2F), and 18% of its reactions allow negative flux, against 8% to 39% in the others (Fig. 2G). The model has 3.7 reactions per gene, more than double the 1.6 to 2.7 of the published models (Fig. 2H). When the fatty acid reactions are excluded, the share of reactions with a gene association rises to 62%, transport reactions make up 21%, 24% of reactions allow negative flux and the number of reactions per gene falls to 2.0, all within or close to the range of the published models (Supplementary Fig. S2B to E).

The functional properties follow the same pattern. Under photoautotrophic conditions iPiccel954 predicts a growth rate of 0.34 h^−1^, which lies within the range of 0.03 to 1.71 h^−1^ reported for the published models (Fig. 2I). In total, 1860 reactions are active, which is 53% of the model and similar to the 51% to 56% of iCre1355, iCZ843 and iAJ526 (Fig. 2J and K). The model has 116 essential genes and 1258 essential reactions, which are 12% of its genes and 36% of its reactions (Fig. 2J and K). Essential genes are therefore a smaller share of the model than in the published models, where they range from 16% to 25%, and essential reactions are a larger share than the 17% to 27% elsewhere (Fig. 2K). The fatty acid reactions explain most of this difference. They account for 1329 of the active reactions and 984 of the essential reactions, because the biomass reaction requires each of the many lipid species it contains. Without them, 531 reactions are active, which is 29% of the remaining 1825, and 274 reactions are essential, which is 15%, while the 116 essential genes make up 12% of the remaining 931 genes.

We tested the model against measurements from four studies of *P. celeri*. At the measured growth rate of 0.323 h^−1^, fluxes per 100 units of net CO_2_ uptake agree with the model in the Calvin-Benson cycle and central metabolism (Steichen et al., 2024) (Fig. 2L). Three of the 11 measured fluxes fall inside the flux ranges allowed by the model, namely RuBisCO oxygenation, mitochondrial pyruvate dehydrogenase and citrate synthase. RuBisCO carboxylation and PGA reduction are within 1.5-fold of the model ranges, with measured values of 120 and 215 against model minimums of 132 and 322. The model also allows flux through SBPase, enolase, PEP carboxylase, malic enzyme and starch synthesis at levels above those measured, which marks the reactions where further constraints from flux data would be most useful. At its default photon bound the model grows at 0.34 h^−1^, which is within about 30% of the rates of 0.26 to 0.32 h^−1^ reported in three studies (Steichen et al., 2024; Torres-Tiji et al., 2026; Weissman et al., 2018) (Fig. 2M). With the light input set to the 60 and 2000 µmolm^−2^s^−1^ used in one of these studies (Weissman et al., 2018), the model grew at 0.007 and 0.16 h^−1^ against the measured 0.06 and 0.26 h^−1^, so growth at high light is within 1.6-fold of the measurement and rises with light as observed. Removing lutein from the biomass also left growth unchanged, in agreement with the lutein-deficient strain (Cano et al., 2023).

### Reconstruction and calibration of the eciPiccel954

We reconstructed the ecGEM from iPiccel954 by splitting reversible reactions into irreversible pairs and assigning a k_cat_ to each of 1282 reactions with TurNuP, which predicts k_cat_ from amino acid sequence and substrates (Kroll et al., 2023) (Fig. 3A). Each k_cat_ gave a protein cost per unit flux, defined as the molecular weight (MW) divided by k_cat_, and an upper bound on the reaction flux. Protein abundances came from the global proteome of wild-type *P. celeri* grown at 2.25% CO_2_ and in air (Torres-Tiji et al., 2026). The proteome was quantified with TMTpro isobaric labeling and expressed as ggDW^−1^ using the measured protein and biomass per cell. We set the protein pool to the summed abundance of the enzymes present in the model, which is 0.0778 ggDW^−1^ in WT-Air and 0.0913 ggDW^−1^ in WT-CO_2_. The pool is 17% higher at elevated CO_2_, whereas growth is 11.6-fold higher.

**Fig. 3.**
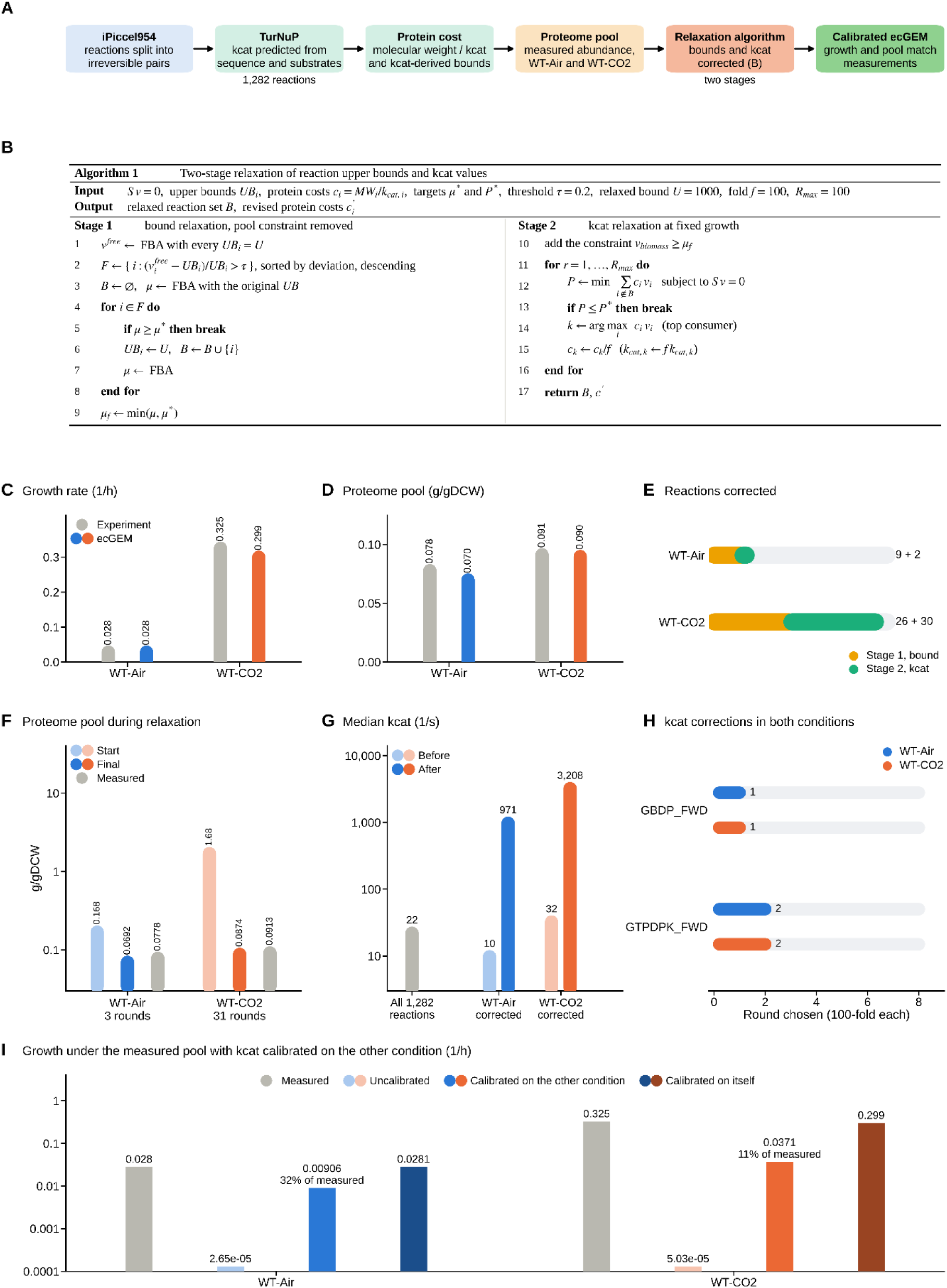
ecGEM reconstruction and relaxation algorithm and leave-out test. (A) ecGEM reconstruction pipeline. (B) Two-stage relaxation algorithm. (C, D) Growth rate and proteome pool, measured and ecGEM. (E) Reactions corrected in each stage. (F) Proteome pool at the start and end of relaxation and measured. (G) Median k_cat_ before and after correction. (H) Reactions corrected in both conditions, by round. (I) Growth under the measured pool with k_cat_ calibrated on the other condition.

The uncalibrated ecGEM could not reproduce the measured growth in either condition. With the k_cat_-derived bounds and the measured pool, growth was near zero or infeasible, and removing the pool constraint alone did not reach the measured growth in either condition. We therefore corrected the reaction bounds and k_cat_ values with a two-stage relaxation algorithm (Fig. 3B). In the first stage, we ranked reactions by how far their flux without the pool constraint exceeded their k_cat_-derived bound, using a threshold of 20%, and raised their bounds to 1000 mmolgDW^−1^h^−1^ until the measured growth was reached or all flagged reactions had been relaxed. Reactions relaxed in this stage were excluded from the pool accounting, because their k_cat_ values were considered unreliable. In the second stage, we fixed growth at the rate reached in the first stage and minimized the protein demand of the remaining reactions. We then raised the k_cat_ of the largest consumer 100-fold and repeated this until the demand fell to the measured pool.

After calibration, the model reproduces the measured growth rate of 0.028 h^−1^ in WT-Air and reaches 0.299 h^−1^ in WT-CO_2_, which is 92% of the measured 0.325 h^−1^, and its pools of 0.0692 and 0.0874 g/gDCW are close to the measured 0.0778 and 0.0913 g/gDCW (Fig. 3C and D). Because the calibration targets these values, the agreement shows that the algorithm converged and is not an independent validation of the model. WT-Air needed 9 bound relaxations and 2 k_cat_ corrections over 3 rounds (Fig. 3E, Supplementary Fig. S3A). WT-CO_2_ needed 26 bound relaxations and 30 k_cat_ corrections over 31 rounds (Fig. 3E, Supplementary Fig. S3B). The first stage could not reach the measured growth in WT-CO_2_ even after all 26 flagged bounds were relaxed, which is why growth stays below the target. The WT-CO_2_ correction is therefore much larger than the WT-Air correction. Before the k_cat_ corrections, the minimized protein demand at the growth reached in the first stage was 0.168 ggDW^−1^ in WT-Air and 1.68 ggDW^−1^ in WT-CO_2_, which is 2.2 and 18 times the measured pools. After correction it fell to 0.0692 and 0.0874 ggDW^−1^ against measured pools of 0.0778 and 0.0913 ggDW^−1^ (Fig. 3F).

The corrected reactions differ from the rest of the network in k_cat_. The median predicted k_cat_ of all 1282 reactions is 22 s^−1^. The median of the corrected reactions was 10 s^−1^ in WT-Air and 32 s^−1^ in WT-CO_2_ before correction, and it rose to 971 and 3208 s^−1^ afterwards (Fig. 3G). Both reactions corrected in WT-Air were also corrected in WT-CO_2_ (Fig. 3H). These reactions, GBDP_FWD and GTPDPK_FWD, were the largest consumers in the first two rounds and each needed one 100-fold correction in both conditions (Fig. 3H). Both are ppGpp reactions that depend on the same gene, and the MEMOTE report flags them as able to carry unbounded flux, so their large protein cost may reflect a network artifact and not a kinetic error. A change of this size can also point to a mismatched k_cat_ source or a missing isozyme or alternative route in the network. The reactions with the largest protein cost before boosting in WT-CO_2_ include glycerate kinase, the two ppGpp reactions, argininosuccinate synthase, phosphoglycolate phosphatase and triosephosphate isomerase (Supplementary Fig. S3C). Most predicted k_cat_ values lie between 1 and 100 s^−1^, so the corrected values are far above the typical range (Supplementary Fig. S3D). In the calibrated models, the ten largest consumers of the pool used 2.7% to 15.6% of the pool each in WT-Air and 0.0% to 1.3% in WT-CO_2_ (Supplementary Fig. S3E). Cost and bound were assigned to 1273 reactions in WT-Air and 1256 in WT-CO_2_, and 253 and 233 of them carry flux (Supplementary Fig. S3F). The difference of 17 reactions corresponds to the difference in bound relaxations between the conditions.

To test whether the corrections transfer between conditions, we applied the corrections found for one condition to the other and maximized growth under the measured pool of the target condition (Fig. 3I). The corrections from WT-CO_2_ gave 0.0091 h^−1^ in WT-Air, which is 32% of the measured 0.028 h^−1^. The corrections from WT-Air gave 0.037 h^−1^ in WT-CO_2_, which is 11% of the measured 0.325 h^−1^. Without corrections, the models reached 2.7 × 10 and 5.0 × 10 h^−1^, and the corrections from the condition itself gave 0.028 and 0.299 h^−1^. The corrections therefore transfer only partly in both directions, and the calibration is a partial test and not a validation (Supplementary Table S3D).

### Nitrogen and carbon are coupled through three nodes

Next, to test whether the measured proteome can account for the faster growth at elevated CO_2_, we used the calibrated ecGEMs of WT-Air and WT-CO_2_ and made urea their only nitrogen source. We blocked two gene-less urea shortcuts, so that urea nitrogen has to pass through urea carboxylase and then allophanate hydrolase, and we assigned glutamate synthase to NADH on the basis of its cofactor-binding pocket. We did not apply the protein pool constraint in this analysis. We instead set the upper bounds of the two urea enzymes directly from their measured abundance, as the product of k_cat_ and abundance divided by MW. Nitrogen and carbon metabolism meet at three nodes in these models. The first is the bicarbonate pool used by urea carboxylase, the second is the recapture of CO_2_ released by allophanate hydrolase, and the third is the 2-oxoglutarate that glutamate synthase consumes.

The capacity of urea carboxylase rises from 0.026 mmolgDW^−1^h^−1^ in WT-Air to 0.277 mmolgDW^−1^h^−1^ in WT-CO_2_, while that of allophanate hydrolase rises from 0.187 to 0.494 mmolgDW^−1^h^−1^ (Fig. 4A). This difference follows from enzyme abundance. Urea carboxylase increases from 0.05 to 0.57 mggDW^−1^, whereas allophanate hydrolase increases from 0.06 to 0.15 mggDW^−1^, and the other enzymes at the three nodes change by 1.2 to 1.5-fold (Fig. 4B). The capacity of urea carboxylase therefore rises 10.8-fold, which matches the 10.8-fold increase in the growth of the urea-fed ecGEMs and approaches the 11.6-fold increase seen in the experiment (Fig. 4C).

**Fig. 4.**
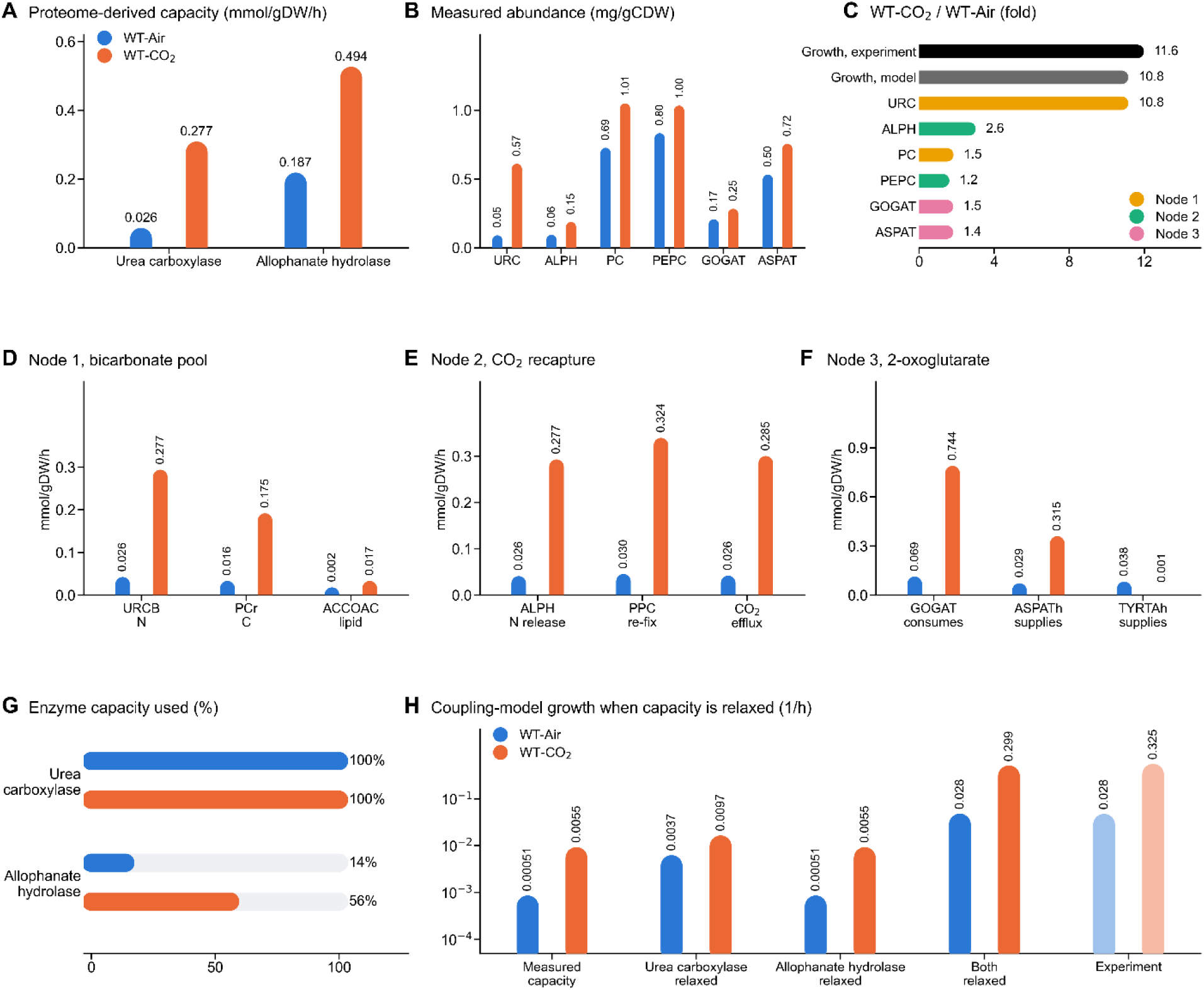
Nitrogen and carbon coupling with urea as the nitrogen source. (A) Capacity of urea carboxylase and allophanate hydrolase from measured abundance. (B) Measured abundance of the coupling enzymes. (C) Fold change of growth and enzyme abundance between WT-CO_2_ and WT-Air. (D) Node 1 fluxes in the bicarbonate pool. (E) Node 2 fluxes in CO_2_ recapture. (F) Node 3 fluxes around 2-oxoglutarate. (G) Share of enzyme capacity used. (H) Growth of the coupling model when enzyme capacity is relaxed.

Flux through the three nodes follows the same pattern. At the bicarbonate pool, urea carboxylase carries 0.026 mmolgDW^−1^h^−1^ in WT-Air and 0.277 in WT-CO_2_, and it competes with pyruvate carboxylase, which carries 0.016 and 0.175 mmolgDW^−1^h^−1^, while acetyl-CoA carboxylase for lipid synthesis carries 0.002 and 0.016 mmolgDW^−1^h^−1^ (Fig. 4D). At the second node, allophanate hydrolase releases CO_2_ at the same rate at which urea carboxylase fixes it, and PEP carboxylase re-fixes 0.030 and 0.324 mmolgDW^−1^h^−1^, which is close to the CO_2_ efflux of 0.026 and 0.285 (Fig. 4E). At the third node, glutamate synthase consumes 0.069 and 0.744 mmolgDW^−1^h^−1^ of 2-oxoglutarate, which aspartate and tyrosine transaminases supply at 0.029 and 0.315 mmolgDW^−1^h^−1^ and at 0.038 and 0.412 mmolgDW^−1^h^−1^ (Fig. 4F). Fluxes through urea carboxylase, PEP carboxylase and glutamate synthase rise about 11-fold, in step with growth.

Urea carboxylase is the limiting step. It runs at 100% of its capacity in both conditions, whereas allophanate hydrolase uses 14% in WT-Air and 56% in WT-CO_2_ (Fig. 4G). At the measured capacity, the urea-fed ecGEMs grow at 0.00051 h^−1^ in WT-Air and 0.0055 h^−1^ in WT-CO_2_, which is about 2% of the measured rates (Fig. 4H). Relaxing urea carboxylase alone increases growth by 627% in WT-Air and by 78% in WT-CO_2_, and relaxing allophanate hydrolase alone has no effect (Fig. 4H, Supplementary Fig. S4A and B). Once urea carboxylase is relaxed, allophanate hydrolase becomes the limit, and relaxing both enzymes raises growth to 0.028 and 0.299 h^−1^, which matches the experiment in WT-Air and reaches 92% of it in WT-CO_2_ (Fig. 4H, Supplementary Fig. S4C). This match is expected, because the capacities no longer limit growth and the models were calibrated to these rates, so it is not an independent test. The independent result is the 10.8-fold scaling of growth with the measured capacity of urea carboxylase, whereas the absolute rates at measured capacity are far below the experiment.

The result is robust to the model assumptions. Using an NADPH-dependent glutamate synthase does not change growth, whereas the original bound file instead of the calibrated one lowers growth to 3 × 10 h^−1^ in WT-Air and 5 × 10 h^−1^ in WT-CO_2_ (Supplementary Fig. S4D). The growth fold of 10.8 holds when both capacities are raised 10-fold and stays at 10.6 when they are free, but it falls to 2.6 when only urea carboxylase is raised 10-fold (Supplementary Fig. S4E). When we relaxed each bound 10-fold in turn, only urea carboxylase changed growth, and the other 216 active bounded reactions changed it by less than 1% (Supplementary Fig. S4F).

### Metabolic bottleneck analysis pinpoints growth limiting enzymes

To find the enzymes that limit growth, we maximized growth under the calibrated protein pool without a growth target, measured how much growth changes when the k_cat_ of each reaction is raised by 10%, and defined the control coefficient as the relative change in growth divided by the relative change in k_cat_ (Fig. 5A). We called a reaction a bottleneck when its coefficient was at least 0.01. We then summed the coefficients of the reactions in each pathway to obtain a pathway control, repeated the analysis after random 2-fold errors in all k_cat_ values, and compared the predictions with the measured proteome. The number of bottlenecks depends on the threshold. In WT-Air it falls from 60 at 0.001 to 8 at 0.03 and to one at 0.1, and in WT-CO_2_ it falls from 90 to 9 and to none (Supplementary Fig. S5A).

**Fig. 5.**
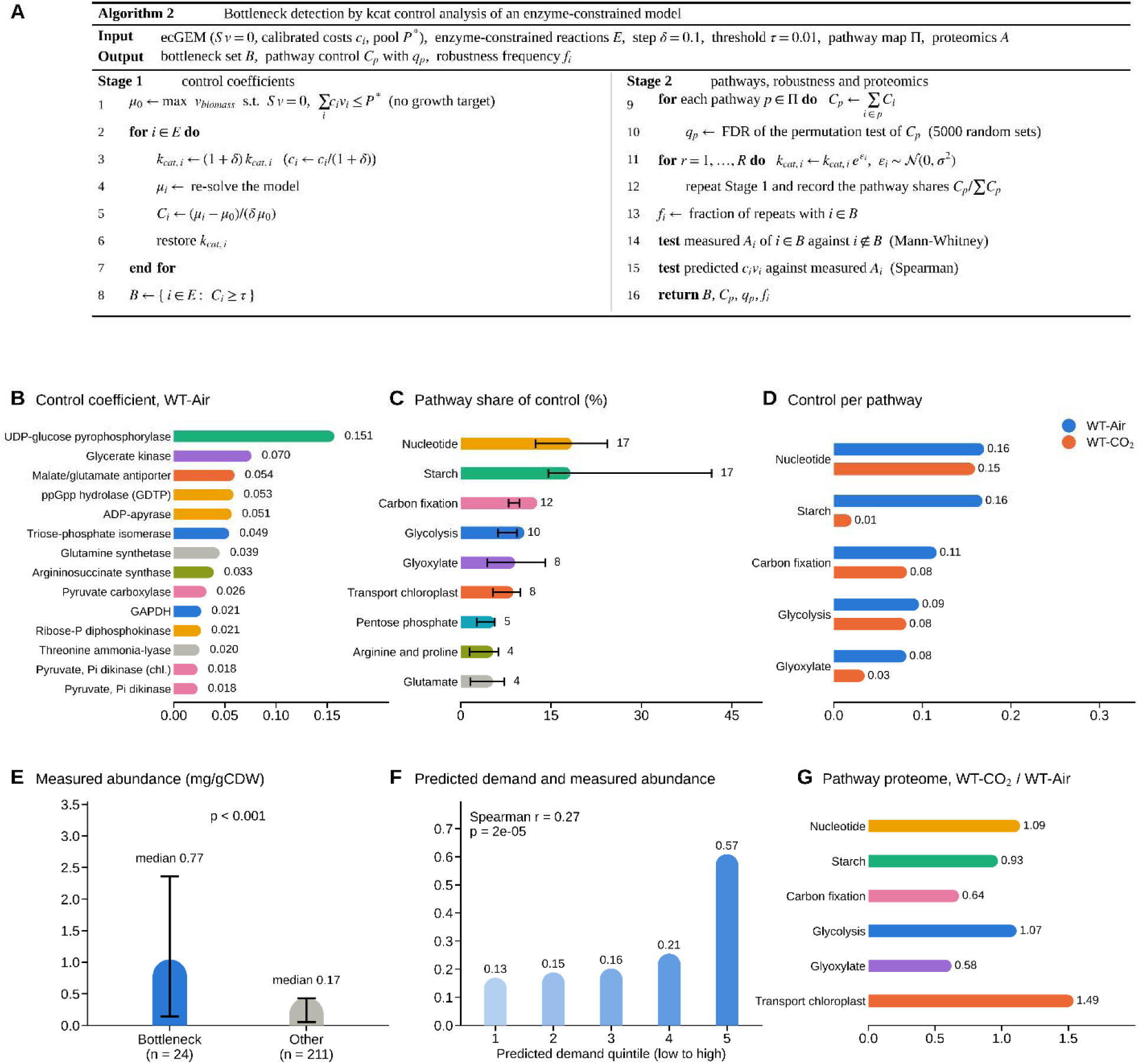
Bottleneck analysis of eciPiccel954. (A) Algorithm for detecting bottlenecks by k_cat_ control analysis. (B) Reactions with the highest control coefficients in WT-Air, colored by pathway. (C) Pathwa share of total control. (D) Control per pathway in WT-Air and WT-CO_2_. (E) Measured abundance of bottleneck enzymes compared with all other enzymes. (F) Measured abundance across quintiles of predicted enzyme demand. (G) Pathway proteome shares in WT-CO_2_ relative to WT-Air.

Both calibrated models have 25 bottleneck reactions, and the control is spread across several pathways. The coefficients sum to 0.93 in WT-Air and 0.92 in WT-CO_2_, which is close to one and shows that the protein pool is the main constraint on growth in both conditions. In WT-Air the largest coefficients belong to UDP-glucose pyrophosphorylase, glycerate kinase, the malate/glutamate antiporter, ppGpp hydrolase and ADP-apyrase, with values of 0.151, 0.070, 0.054, 0.053 and 0.051 (Fig. 5B). Nucleotide metabolism and starch metabolism each hold 17% of the total control, followed by carbon fixation with 12%, glycolysis with 10%, and glyoxylate metabolism and chloroplast transport with 8% each (Fig. 5C). No pathway holds more control than expected by chance after correction for multiple testing, since the lowest q value is 0.17, so the bottlenecks are better described as distributed across the network than as enriched in particular pathways. This is what one expects when many enzymes compete for a limited protein pool.

Control at elevated CO_2_ is also spread over 25 reactions, but they are largely different ones. Only UDP-glucose pyrophosphorylase is a bottleneck in both conditions, and its coefficient falls from 0.151 in WT-Air to 0.012 in WT-CO_2_. The largest coefficients in WT-CO_2_ belong to the two ppGpp reactions, with 0.053 each, followed by sulfate transport, argininosuccinate lyase and the cytochrome b6/f complex, with 0.050, 0.046 and 0.044, and photosynthesis holds 13% of the control against 2% in WT-Air. Starch metabolism loses almost all its control, which falls from 0.16 in WT-Air to 0.01 in WT-CO_2_, and glyoxylate metabolism falls from 0.08 to 0.03, whereas nucleotide metabolism keeps 0.15 and carbon fixation and glycolysis keep 0.08 each (Fig. 5D).

The protein pool is the constraint that creates these bottlenecks. Lowering the protein cost of a single reaction 10-fold, 100-fold or to zero gives at most 17%, 19% and 20% more growth in WT-Air, and only one reaction gives 10% or more in each case (Supplementary Fig. S5B). Growth also scales with the pool in WT-Air, at 25%, 50% and 75% of the calibrated growth when the pool is reduced to 0.25, 0.5 and 0.75 times the calibrated value, and at 1.5, 2 and 4 times the calibrated growth when the pool is enlarged by the same factors (Supplementary Fig. S5C). In WT-CO_2_ growth also falls in proportion when the pool is reduced, but it rises by only 6% when the pool is enlarged, because the model then reaches its maximal growth of 0.337 h^−1^ (Supplementary Fig. S5C). The size of the k_cat_ step has no effect on the result, since steps of 5%, 10% and 20% each give 25 bottlenecks, all of them shared, and the coefficients at 5% and 10% have a Spearman correlation of 0.98 (Supplementary Fig. S5D).

The measured proteome supports the predictions. The 24 bottleneck enzymes of WT-Air have a median abundance of 0.77 mggDW^−1^, against 0.17 mggDW^−1^ for the 211 other enzymes (Fig. 5E, p < 0.001). The demand predicted by the model, which is the protein cost multiplied by the flux, correlates weakly with the measured abundance (Spearman r = 0.27, p = 2 × 10), and the enzymes in the highest quintile of predicted demand have a median abundance of 0.57 mggDW^−1^, against 0.13 to 0.21 mggDW^−1^ in the lower quintiles (Fig. 5F). When we compared the two conditions, the abundance of carbon fixation enzymes was 36% lower at elevated CO_2_ and that of glyoxylate enzymes was 42% lower, which is consistent with the lower control of both pathways (Fig. 5G). The pattern is weaker elsewhere, since starch metabolism is only 7% less abundant while its control falls almost to zero, and nucleotide metabolism is 9% more abundant and keeps its control (Fig. 5G).

The identity of individual bottlenecks is sensitive to k_cat_ error, but the set of limiting reactions is stable. In 6 draws with 2-fold random errors, each of the twelve highest-ranked reactions in WT-Air was called a bottleneck in 67% to 100% of the draws (Supplementary Fig. S5E). The ranking of the control coefficients agreed well with the calibrated ranking in all draws, with Spearman correlations of 0.84 to 0.97 (Supplementary Fig. S5F). The number of bottlenecks in WT-Air ranged from 12 to 23, and in WT-CO_2_ it ranged from 20 to 23 in five draws and was zero in one draw in which growth reached its maximum and the pool no longer limited it (Supplementary Fig. S5G). The top reaction of the calibrated WT-CO_2_ model, a ppGpp reaction, was the top reaction in none of the draws, although each of the twelve most frequent reactions was a bottleneck in 5 of the 6 draws (Supplementary Fig. S5H). The spread of control over many reactions is therefore robust, whereas the enzyme that holds the largest share is not. We therefore regard individual reactions as candidates and place more weight on the role of the protein pool.

## DISCUSSION

The iPiccel954 matches the experimental outcomes in the growth rate at high light, in the fluxes through RuBisCO carboxylation and PGA reduction, and in the unchanged growth of a lutein-deficient strain (Cano et al., 2023; Steichen et al., 2024; Torres-Tiji et al., 2026; Weissman et al., 2018), which supports its use as the basis of the ecGEM. The central result is that *P. celeri* does not grow 11.6-fold faster at elevated CO_2_ by building 11.6 times more enzymes. The measured protein pool is only 17% larger, so growth per unit of protein rises about 10-fold. The uncorrected models show the same tension. Their minimal protein demand scales with growth, while the measured pools barely move. Closing this gap required 9 bound relaxations and 2 k_cat_ corrections in air and 26 bound relaxations and 30 k_cat_ corrections at elevated CO_2_. The two k_cat_ corrections needed for the slow state are therefore both among those needed for the fast state. Applying the corrections of one condition to the other recovered only 32% of the measured growth in air and 11% at elevated CO_2_ (Fig. 3I), so the corrections are specific to each condition. The same enzymes may therefore run at a much higher fraction of their capacity when CO_2_ is plentiful, which would be a protein efficiency effect (Chen and Nielsen, 2019). The TurNuP k_cat_ values (Kroll et al., 2023) were low to fit the measured pool in both conditions, and the shortfall was larger at elevated CO_2_, where 30 reactions needed their k_cat_ raised against 2 in air. Machine-learning k_cat_ predictions are known to improve proteome-limited models but remain uncertain (Heckmann et al., 2018), and k_cat_ curation is a standard step in building ecGEMs (Domenzain et al., 2022).

The protein pool is the real limit in both conditions, and what changes with CO_2_ is where the control sits. In air, 25 reactions share control, spread over nucleotide metabolism, starch metabolism, carbon fixation, glycolysis and organellar transport. Lowering the protein cost of any single reaction raises growth by at most 20%, whereas enlarging the pool raises growth in proportion, so capacity and not any single kinetic step caps growth. Diffuse control is what a proteome allocated close to optimally looks like, because control is shared among enzymes and cannot be concentrated in one (Kacser et al., 1995). Elevated CO_2_ moves this pattern and does not break it. Control stays spread over 25 reactions, almost none of which limit growth in air, starch metabolism loses the 17% of the control it holds in air, and photosynthesis gains it. Carbon fixation keeps part of its control while its enzymes become 36% less abundant. The cell appears to release proteome from carbon fixation once CO_2_ is no longer scarce, an allocation shift that resembles the shifts between proteome sectors seen when substrates change (Basan et al., 2015; Chowdhury et al., 2025; Jahn et al., 2018). Part of the remaining control sits in amino acid synthesis, for example at argininosuccinate lyase, where enzymes are consistently highly saturated across organisms and conditions (Domenzain et al., 2022). We interpret these patterns cautiously. No pathway holds more control than expected by chance after correction for multiple testing, three of the highest ranked reactions are ppGpp reactions that the MEMOTE report flags as able to carry unbounded flux, and the top reaction of WT-CO_2_ was not the top reaction in any draw with two-fold k_cat_ errors. The number of bottlenecks and the spread of control are therefore robust features of the model, whereas the named enzymes depend on the k_cat_ values.

The nitrogen coupling shows which part of the response is built by enzyme amount and which is not. Urea carboxylase abundance rises 10.8-fold, in step with growth, and the enzyme runs at full capacity in both conditions. Allophanate hydrolase rises only 2.6-fold and uses 14% of its capacity in air and 56% at elevated CO_2_. Pyruvate carboxylase and PEP carboxylase, which supply carbon skeletons for nitrogen assimilation, change by 1.5-fold or less. The pattern suggests that the nitrogen entry step is scaled to demand, the next step keeps spare capacity, and the carbon supply to the nitrogen route is met mainly by greater substrate availability and not by more enzymes. This matches the long-standing view that carbon and nitrogen assimilation are tied through carbon skeletons and anaplerotic carbon fixation (Huppe and Turpin, 1994). The 10.8-fold scaling of the model matches the 11.6-fold observed growth, which is the one result in this work that was not fitted. It must still be read with care, because at measured capacity the model reaches only about 2% of the observed rate.

Two testable predictions follow. First, in either condition, no single k_cat_ improvement should raise growth by more than about 20%, and only a larger protein pool should raise it in proportion. Second, the limiting enzymes should shift with CO_2_ supply, with UDP-glucose pyrophosphorylase as the first candidate in air, so its overexpression may increase growth in air but not at elevated CO_2_, where its control falls from 0.151 to 0.012. Table 1 lists these predictions and further candidate experiments ranked by the strength of the model evidence. The proteome data support the direction of these predictions, since bottleneck enzymes are more abundant than the rest (0.77 against 0.17 mggDW^−1^), although predicted demand and abundance correlate only weakly overall (r = 0.27). These conclusions also depend on calibration to a single growth rate per condition, on corrections that transfer only partly between conditions, on a WT-CO_2_ model that reaches 92% of the measured growth, and on bottleneck identities that shift under two-fold k_cat_ error, so we regard the ranked bottlenecks as potential candidates. The base model also grows linearly with absorbed light and lacks saturation and photoprotection, so its growth at the two light levels from literature (Weissman et al., 2018) is underestimated and its light response should be read with care. Our use of TurNuP k_cat_ values in an ecGEMs follows the *C. ohadii* model (Soleymani et al., 2025), and proteomics-based kinetic inference has also been applied to *C. reinhardtii* (Arend et al., 2023).

**Table 1.** Candidate experiments ranked by the strength of the model evidence. Targets are candidates, and the rankings depend on the predicted k_cat_ values.

| Rank | Target | Proposed experiment | Model prediction | Supporting evidence |
|---|---|---|---|---|
| 1 | Urea carboxylase | Overexpress in air and at elevated CO_ with urea as the sole nitrogen source and measure growth | Relaxing this step alone raises growth by 627% in air and 78% at elevated CO_ | Runs at 100% of capacity in both conditions, abundance rises 10.8-fold in step with growth, and it was the only one of 217 bounded reactions that changed growth when relaxed |
| 2 | UDP-glucose pyrophosphorylase | Overexpress in air and at elevated CO_ and compare growth | Growth gain in air but not at elevated CO_, since control falls from 0.151 to 0.012 | Largest control coefficient in air, and a bottleneck in 67% to 100% of the random k<sub>cat</sub> draws |
| 3 | Allophanate hydrolase together with urea carboxylase | Co-overexpress both enzymes and compare them with urea carboxylase alone | Allophanate hydrolase becomes limiting once urea carboxylase is relaxed, and relaxing both reaches 0.028 and 0.299 h<sup>−1</sup> | Uses 14% of its capacity in air and 56% at elevated CO_ |
| 4 | Urea supply matched to CO_ | Vary urea concentration across CO_ levels and measure growth and urea use | Urea beyond the carboxylation capacity at a given CO_ level is wasted | Urea carboxylase runs at full capacity while allophanate hydrolase keeps spare capacity |
| 5 | Starch metabolism | Reduce starch synthesis activity in air and at elevated CO_ and compare growth | Growth should respond in air but not at elevated CO_ | Control falls from 0.16 in air to 0.01 at elevated CO_ |

These results point to several biotechnological implications (Alsiyabi et al., 2022). Because growth is limited by the size of the protein pool and not by any single enzyme, strain engineering that targets individual enzymes is unlikely to give large gains, and CO_2_ supplementation is the more effective lever. The model points to UDP-glucose pyrophosphorylase in air and to nucleotide metabolism, photosynthetic electron transport and amino acid synthesis at elevated CO_2_, so strains with more active forms of these enzymes are a first test for further gains in growth, although the identity of these enzymes is sensitive to k_cat_ error. The nitrogen coupling suggests that urea feeding should be matched to carbon supply, since urea carboxylase runs at full capacity in both conditions while allophanate hydrolase keeps spare capacity. A urea supply that exceeds what the cell can carboxylate at a given CO_2_ level would then be wasted. More broadly, the calibration algorithm gives a route to ecGEMs for fast-growing algae, where predicted k_cat_ values alone do not fit the measured protein pool. It ranks the enzymes that most need experimental k_cat_ values, so measurement effort can be directed to the reactions that matter most for growth.

## MATERIALS AND METHODS

### Reconstruction and manual curation of iPiccel954

*Draft reconstruction.* The draft model was generated from the *P. celeri* genome (Becker et al., 2020) with the RAPS (Metcalf et al., 2020), using the iCre1355 as a template. RAPS adds a reaction when its best BLAST match has an E-value below 10^−20^ and flags weaker matches for manual review. The draft was then compared by homology with the iPicre1230. Reactions present in both models and reactions present only in the draft were kept, and reactions present only in iPicre1230 were added sparingly. The final model was named iPiccel954.

*Biomass reaction.* The composition of *P. celeri* TG2 under nutrient-replete conditions (LaPanse et al., 2024) provided the percent ash-free dry weight of protein, carbohydrate, fatty acid methyl esters, nucleotides and chlorophyll. These five fractions were divided by their sum, so that the modeled biomass totaling 1000 mg. The target mass of each pool was divided among its species, and each coefficient was calculated as mass divided by MW. Nucleotides were split into 90% RNA and 10% DNA. The nucleotide proportions followed the genomic G+C content of 46%, with G equal to C and A equal to U or T. Protein kept the amino acid ratios of the draft, except for methionine and valine, which were set to the reported values, and the other amino acids were scaled to keep the total mass. Carbohydrate used the measured proportions of glucose, mannose, galactose and ribose, with glucose assigned to starch. Chlorophyll a and b were set to a 3 to 1 mass ratio. Other pools, including the carotenoids, kept their draft coefficients. The growth-associated maintenance term was not changed from iCre1355. After all formula corrections, the non-GAM coefficients were scaled so that the total mass of the biomass reaction, including the maintenance term, equaled 1 gmmol^−1^, as the MEMOTE test requires (Lieven et al., 2020).

*Elemental balance.* For each reaction with a hydrogen-only imbalance, the coefficient of the proton species of the matching compartment was adjusted by the imbalance. The change was kept only if the optimal growth rate stayed the same within 10^−6^. Remaining imbalances were corrected by changing the formula of the metabolite whose identifier matched the reaction identifier. This was repeated in passes until no further correction was found, and a change was kept only if it balanced the reaction, did not unbalance other reactions sharing the metabolite, and did not reduce growth. Reactions with the photon element Z, generic R groups and the starch polymer formula were excluded.

*Annotation.* Cross-database annotations were transferred from the BiGG universal model (Norsigian et al., 2020) by matching, in order, the identifier, the original BiGG identifiers in the universal model, and for metabolites the identifier without the compartment suffix. SBO terms were assigned by reaction type. These were SBO:0000629 for biomass, SBO:0000627 for exchange, SBO:0000628 for demand, SBO:0000185 for transport (reactions with metabolites in more than one compartment) and SBO:0000176 for other reactions. Metabolites received SBO:0000247 and genes SBO:0000243.

*Energy-generating cycles.* Following previous work (Fritzemeier et al., 2017), as implemented in MEMOTE, we tested ten energy metabolites (ATP, GTP, CTP, UTP, ITP, NADPH, NADH, FADH2, acetyl-CoA and the glutamate and 2-oxoglutarate couple). For each, all boundary reactions were closed, a dissipation reaction was added, and its maximum flux was calculated. A positive maximum indicated a cycle. In each iteration, the reversible reaction without a gene association that carried the largest flux was restricted to one direction. The restriction was kept if growth was unchanged and the dissipation flux fell, and the procedure stopped if only gene-associated reactions remained.

*Unbounded flux.* Flux variability analysis at the default bounds flagged reactions whose minimum or maximum reached the bound of 1000 mmolgDW^−1^h^−1^. Reversible flagged reactions were restricted to one direction in two rounds. Round 1 removed the direction pinned at the bound (the direction of flux in the optimal solution when both were pinned), and round 2 removed the opposite direction for reactions that could not be restricted in round 1. Irreversible flagged reactions were then blocked entirely (bounds set to zero) in a third round. Each round was applied as a batch and reduced by bisection to the largest set that kept growth above 80% of its baseline. Reactions fixed at zero were removed, followed by metabolites and genes that no reaction used.

### Construction of the enzyme-constrained model eciPiccel954

We built the ecGEM with the standard formulation (Sanchez et al., 2017). Every reversible reaction of the curated model was split into a forward and a reverse reaction (suffixes _FWD and _REV) so that all fluxes are non-negative. We predicted k_cat_ for the enzymes of the model with TurNuP (Kroll et al., 2023), which supplied k_cat_ values for 1282 irreversible reactions. The protein cost of each reaction was its enzyme MW divided by its k_cat_.

Protein abundances in ggDW^−1^ came from the previous work of the authors (Torres-Tiji et al., 2026) for cells grown in air and at elevated CO_2_. For each condition, the protein pool was the summed abundance of the enzymes present in the model. The upper bound of each reaction was the k_cat_ multiplied by the measured abundance of the associated protein and divided by its MW. Reactions without a gene association kept the bounds of the curated model. A single constraint required the total protein cost of all fluxes to remain at or below the protein pool of the condition. The model maximized the biomass flux under this constraint, and flux distributions were obtained by parsimonious flux balance analysis. All models were built with COBRApy (Ebrahim et al., 2013) and solved with the Gurobi solver.

### Calibration of k_cat_ values against the measured protein pool

The ecGEMs could not reach the measured growth rates under the measured protein pools. Maximizing growth with the pool constraint and all reaction bounds at 1000 mmolgDW^−1^h^−1^, or with the predicted bounds and no pool constraint, also did not reach the measured growth rates. We therefore calibrated each condition in two stages (Algorithm 1).

Stage 1 relaxed reaction bounds. We solved the model without the pool constraint, once with the predicted bounds and once with all bounds at 1000 mmolgDW^−1^h^−1^. A reaction was flagged when its flux in the second solution exceeded its predicted bound by more than 20%. Flagged reactions were ranked by this relative excess, with the largest first, and their bounds were set to 1000 mmolgDW^−1^h^−1^ one at a time. Growth was recalculated after each relaxation, until the measured growth rate was reached or all flagged reactions had been relaxed. The relaxed reactions were excluded from the protein pool constraint in all later steps, including the final run, because their k_cat_ were judged unreliable.

Stage 2 corrected the k_cat_. We fixed the biomass flux at the growth rate reached in Stage 1 and minimized the total protein cost of the fluxes through all reactions that were not relaxed in Stage 1. This minimal cost is the protein demand at that growth rate. In each round, the reaction with the largest share of the demand was identified, its k_cat_ was raised 100-fold. A reaction could be selected in more than one round, and its corrections accumulated. The procedure stopped when the demand fell to the measured pool or below, or after 100 rounds.

Finally, we reran the ecGEM with the corrected protein costs and bounds and the measured pool, maximized growth, and obtained the flux distributions by parsimonious flux balance analysis. These fluxes were used for all later analyses. The settings of both stages (20%, 1000 mmolgDW^−1^h^−1^, a factor of 100 and 100 rounds) were the same in both conditions.

To test whether the corrections transfer between conditions, we applied the corrections of one condition to the other. For each target condition we used its own upper bounds and protein costs, raised the bounds and divided the protein costs of the reactions corrected in the other condition by the cumulative fold applied there, excluded the relaxed reactions from the pool, and maximized growth under the measured pool of the target condition. We also maximized growth without any correction.

### Simulation of carbon and nitrogen coupling with urea as sole nitrogen source

The coupling analysis used the calibrated ecGEMs of WT-Air and WT-CO_2_ with the revised reaction bounds, without the protein pool constraint, so that only the bounds of individual enzymes limited the fluxes. We made three changes to represent growth on urea. First, urea was the only nitrogen source, and every other import of a nitrogen-containing compound was closed. Second, the two urea-splitting reactions without a gene association (uahCv and ATAH_REV) were blocked, so that urea could be used only through urea carboxylase (URCB) and allophanate hydrolase (ALPH). Third, glutamate synthase (GOGAT) uses NADH and not NADPH in the curated model, and we also tested an NADPH-dependent version.

We rebuilt the bounds of urea carboxylase and allophanate hydrolase for each condition. The genes of the two enzymes were identified from the annotations of the protein sequences. The bound of each reaction was the TurNuP k_cat_ multiplied by the measured abundance of the enzyme and divided by its MW, calculated from the sequence. This enzyme capacity was converted to mmolgDW^−1^h^−1^. We maximized growth at these bounds and obtained flux distributions by parsimonious flux balance analysis. The use of each enzyme was its flux as a percentage of its capacity.

To test which capacity limits growth, we solved the model with the measured capacities, with the capacity of urea carboxylase relaxed to 1000 mmolgDW^−1^h^−1^, with the capacity of allophanate hydrolase relaxed, and with both relaxed. We also scaled each capacity by 0.5, 1, 2, 5 and 10, and removed the bound entirely. To test whether other enzymes could limit growth, we raised the upper bound of each active reaction with a bound ten-fold, one at a time, and recorded the change in growth. The fold change between conditions was calculated as the ratio of the value at elevated CO_2_ to the value in air, for the abundance of each enzyme and for the capacity-limited growth.

We examined three nodes where carbon and nitrogen metabolism meet. These were the cytosolic bicarbonate pool shared by urea carboxylase, pyruvate carboxylase and acetyl-CoA carboxylase, the recapture of CO_2_ by PEP carboxylase, and the 2-oxoglutarate node at glutamate synthase and the transaminases. The fluxes through these nodes were taken from the parsimonious solutions of the measured-capacity model

### K_cat_ control analysis of growth-limiting enzymes

We quantified the control of each enzyme over growth with a k_cat_ control coefficient, calculated separately for each condition on the calibrated enzyme-constrained model at the pool-limited optimum. We removed the growth target and the k_cat_-derived bounds, so that the protein pool constraint alone limits growth. We first maximized growth under the measured pool. Then, for each reaction with a protein cost, we raised the k_cat_ by 10%, which lowered the protein cost by the same factor, maximized growth again and restored the original value before the next reaction. The coefficient of a reaction was the relative change in the maximal growth rate divided by the relative increase in its k_cat_. Reactions with a coefficient of 0.01 or higher were called growth-limiting bottlenecks.

Reactions were assigned to pathways with the pathway groups of the curated model, and forward and reverse reactions took the pathway of their parent reaction. Reactions without a pathway were excluded from the pathway analysis. The control of a pathway was the sum of the coefficients of its reactions, and its share was this sum divided by the total over all reactions. For pathways with at least five reactions, we tested whether the control was higher than expected by chance. We drew 5000 random sets of reactions of the same size and compared their summed coefficients with the observed sum in a one-sided test. P values were calculated as one plus the number of random sets at or above the observed value, divided by 5001, and corrected for multiple testing by the Benjamini-Hochberg procedure.

We compared the results with the measured proteome. The abundance of the enzyme of each reaction was evaluated from the gene-protein-reaction rule, with isozymes summed and subunits of a complex represented by the lowest abundance. Reactions without a gene association were excluded. We compared the abundance of bottleneck enzymes with that of the other active enzymes with a Mann-Whitney test. We calculated the Spearman correlation between the predicted demand of each reaction (its protein cost multiplied by its flux) and its measured abundance in air. For each pathway, we compared the share of the summed abundance in air with the share at elevated CO_2_ and report their ratio.

### Sensitivity of bottlenecks to model parameters and k_cat_ error

We tested how the bottlenecks depend on the threshold, the step size, the protein pool and the k_cat_. To test the threshold, we counted bottleneck reactions for cutoffs of the control coefficient from 0.001 to 0.1. To test the step size, we repeated the control analysis in WT-Air with k_cat_ increases of 5%, 10% and 20%, compared the sets of bottleneck reactions, and calculated the Spearman correlation of the coefficients.

To test whether growth is limited by the cost of a single reaction, we lowered the protein cost of one reaction at a time in WT-Air by 10-fold, by 100-fold and to zero, and recorded the relative gain in growth. We did this for all reactions with a protein cost. To test the effect of the pool, we multiplied the measured pool by 0.25, 0.5, 0.75, 1, 1.5, 2 and 4, and also removed it, and maximized growth in both conditions.

To test the effect of errors in the k_cat_, we multiplied each k_cat_ in WT-Air by a random factor, the exponential of a normal variable with mean zero and standard deviation ln 2, which corresponds to a typical two-fold error. The factor changed the protein cost of each reaction. The model was not recalibrated, so each draw gave its own growth rate. We then recalculated the control coefficients of all reactions that carried flux, with the 10% step. The whiskers in Fig. 5C show the 10th to 90th percentile of the pathway shares over 6 draws. For Fig. S5 we used 6 draws for WT-Air and 6 draws for WT-CO_2_. We recorded how often each of the twelve reactions with the highest coefficients had a coefficient of 0.01 or more, and the Spearman correlation between the coefficients of each draw and those of the calibrated model, calculated over their shared reactions. We repeated the draws for WT-CO_2_, counted the bottleneck reactions in every draw of both conditions, and recorded how often each reaction was a bottleneck in WT-CO_2_.

## Supporting information

which brought protein to the dominant fraction and the total to 1000 gmol-1

The corrections therefore transfer only partly in both directions, and the calibration is a partial test and not a validation

## DATA AVAILABILITY

All the data and codes are available at https://github.com/pnnl/p_celeri_GEM.

## AUTHOR CONTRIBUTION STATEMENT

N.B.C. conceived the study, performed all modeling and analysis, and wrote the original draft. Y.T.T. conceived the study, contributed to the writing of the initial draft, and edited the manuscript. A.C. and M.C.P. supervised the work and edited the manuscript. All authors read and approved the final manuscript.

## ACKNOWLEDGEMENT

This work used proteomics data generated on project awards from the Environmental Molecular Sciences Laboratory (EMSL), a DOE Office of Science User Facility sponsored by the Biological and Environmental Research program under Contract No. DE-AC05-76RL01830. We thank Sarah Leichty and Rick Washburn of the EMSL User Program Services team for managing the projects that produced these data, and the Environmental Health and Safety office of the Colorado School of Mines for supporting them. We also thank the U.S. Department of Energy for its sustained support of algal research.

## SUPPLEMENTARY FILES

**Supplementary Fig. S1** Curation of iPiccel954. Shows the largest biomass components, residual elemental imbalance, reactions per pathway, gene association, gene and reaction multiplicity, and the most connected metabolites.

**Supplementary Fig. S2** Structural and functional comparison with other algal models. Compares gene association by pathway and the structural, active and essential reaction and gene counts, including iPiccel954 without fatty acid reactions.

**Supplementary Fig. S3** Calibration of the enzyme-constrained models. Shows the protein pool demand over the relaxation rounds, the boosted reactions, the range of k_cat_ values, the pool allocation among top consumers, and the constrained reactions.

**Supplementary Fig. S4** Coupling of nitrogen and carbon assimilation. Tests the sensitivity of growth to the capacities of urea carboxylase and allophanate hydrolase, the model assumptions, the growth fold change between conditions, and a ten-fold relaxation of single bounds.

**Supplementary Fig. S5** Robustness of the bottleneck analysis. Shows how the bottlenecks depend on the control coefficient threshold, on relaxing one reaction at a time, on pool size, on the k_cat_ step size, and on random k_cat_ errors.

**Supplementary Table S1.** Biomass composition of iPiccel954 before and after curation, by metabolite and class.

**Supplementary Table S2.** Structural properties of iPiccel954 and other algal models.

**Supplementary Table S3.** Calibration of the enzyme-constrained models, with the growth and pool summary, the relaxed bounds, the boosted kcat values and the leave-out test.

**Supplementary Table S4.** Bottleneck enzymes and pathway shares of k_cat_ control, with measured enzyme abundance in air and elevated CO_2_.

**Supplementary Table S5.** Curation log of iPiccel954, with the stage summary, the logged corrections and the remaining imbalances.

