## Supplementary material for "Enzyme-Constrained Metabolic Modeling Reveals CO_2_-Dependent Growth Control in *Picochlorum celeri*": which brought protein to the dominant fraction and the total to 1000 gmol-1

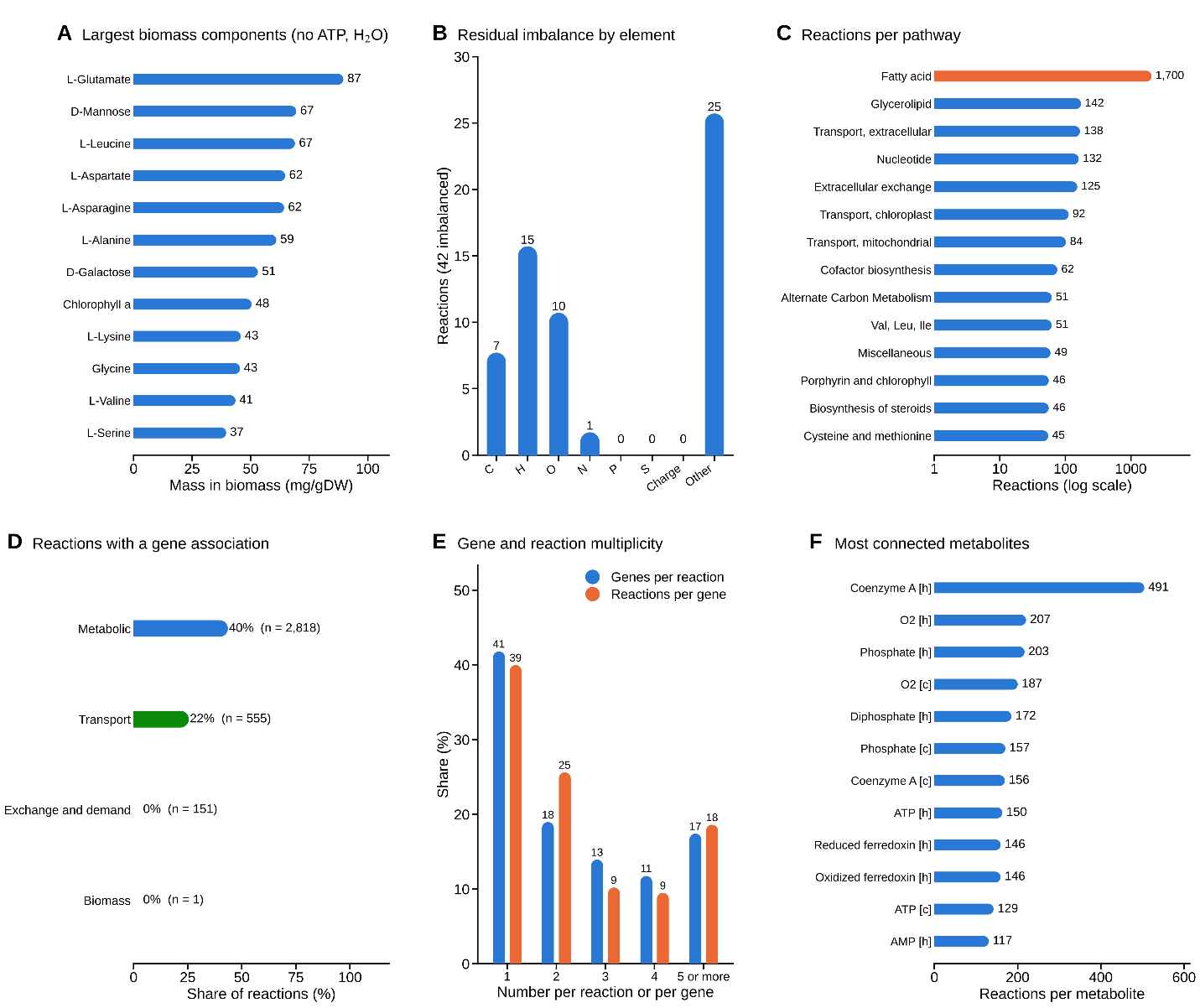

**Supplementary Figure S1. Content and quality of the iPiccel954 model.** (A) Largest biomass components by mass. (B) Residual imbalance of the final model by element. (C) Reactions per pathway. (D) Reactions with a gene association by reaction class. (E) Genes per reaction and reactions per gene. (F) Most connected metabolites.

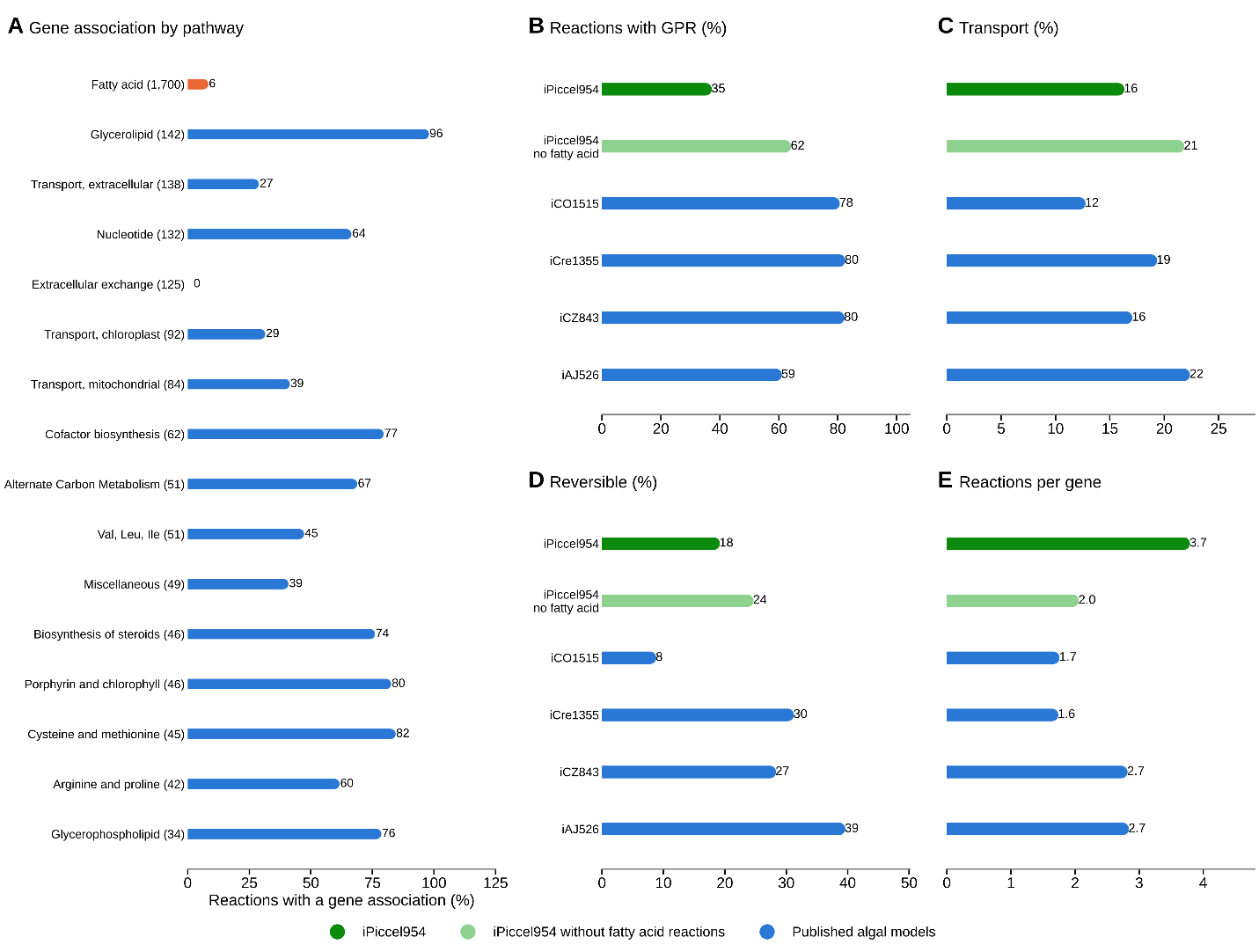

**Supplementary Figure S2. Gene association and the comparison without fatty acid reactions.** (A) Percentage of reactions with a gene association in the 16 largest pathway groups, with the number of reactions in brackets. Fatty acid metabolism is shown in orange. (B) Reactions with a gene-protein-reaction rule (%). (C) Transport reactions (%). (D) Reversible reactions (%). (E) Reactions per gene. In B to E, iPiccel954 is dark green, iPiccel954 without fatty acid reactions is light green, and published models are blue.

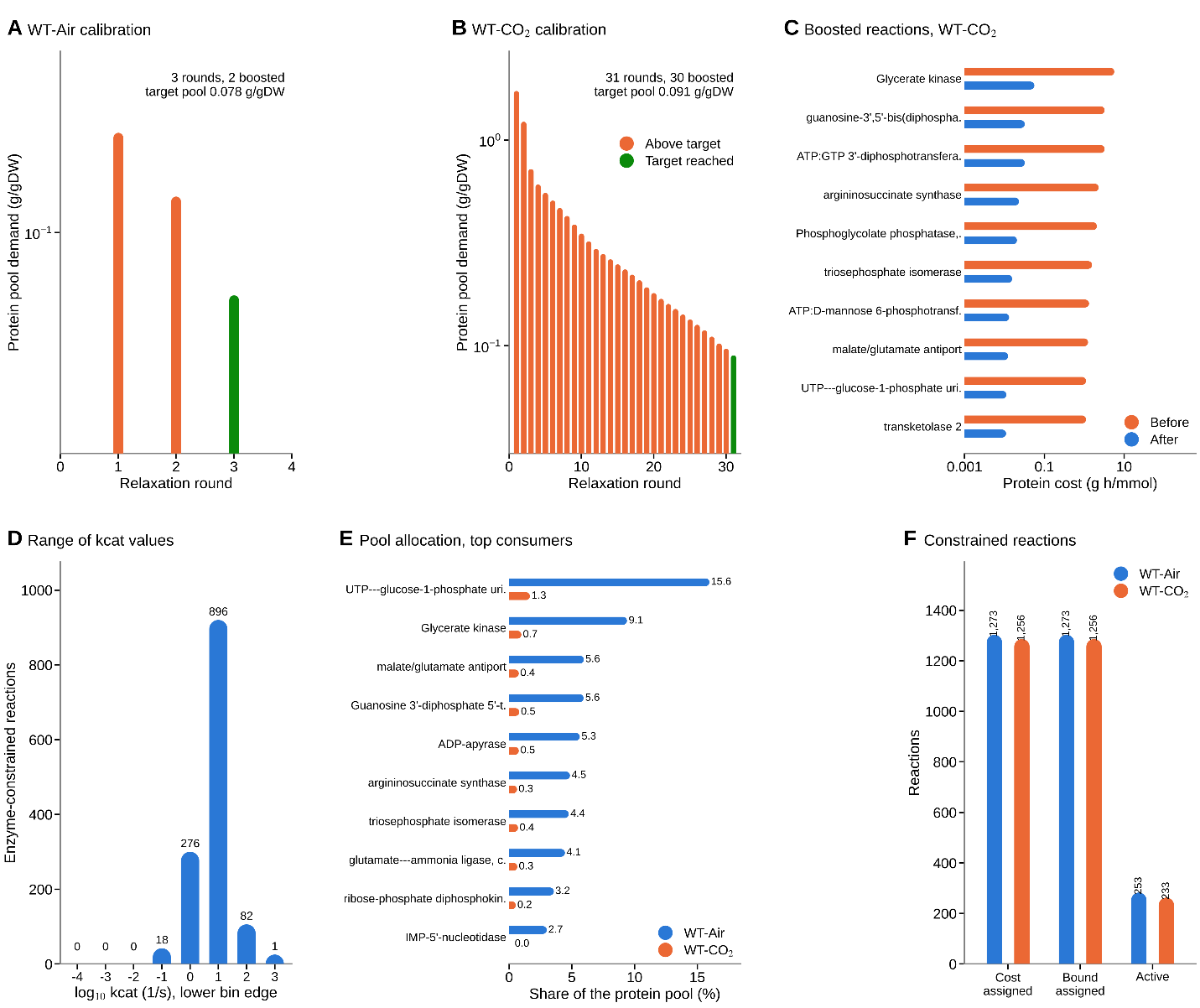

**Supplementary Figure S3. Calibration of the protein pool.** (A) Protein pool demand in each relaxation round for WT-Air. (B) The same for WT-CO2. (C) Protein cost of the ten most costly boosted reactions in WT-CO_2_, before and after the boost. (D) Range of the TurNuP k_cat_ values of the enzyme-constrained reactions. (E) The ten largest consumers of the protein pool in WT-Air and their share in WT-CO_2_. (F) Reactions with a protein cost, with a k_cat_-derived bound and carrying flux in the calibrated models.

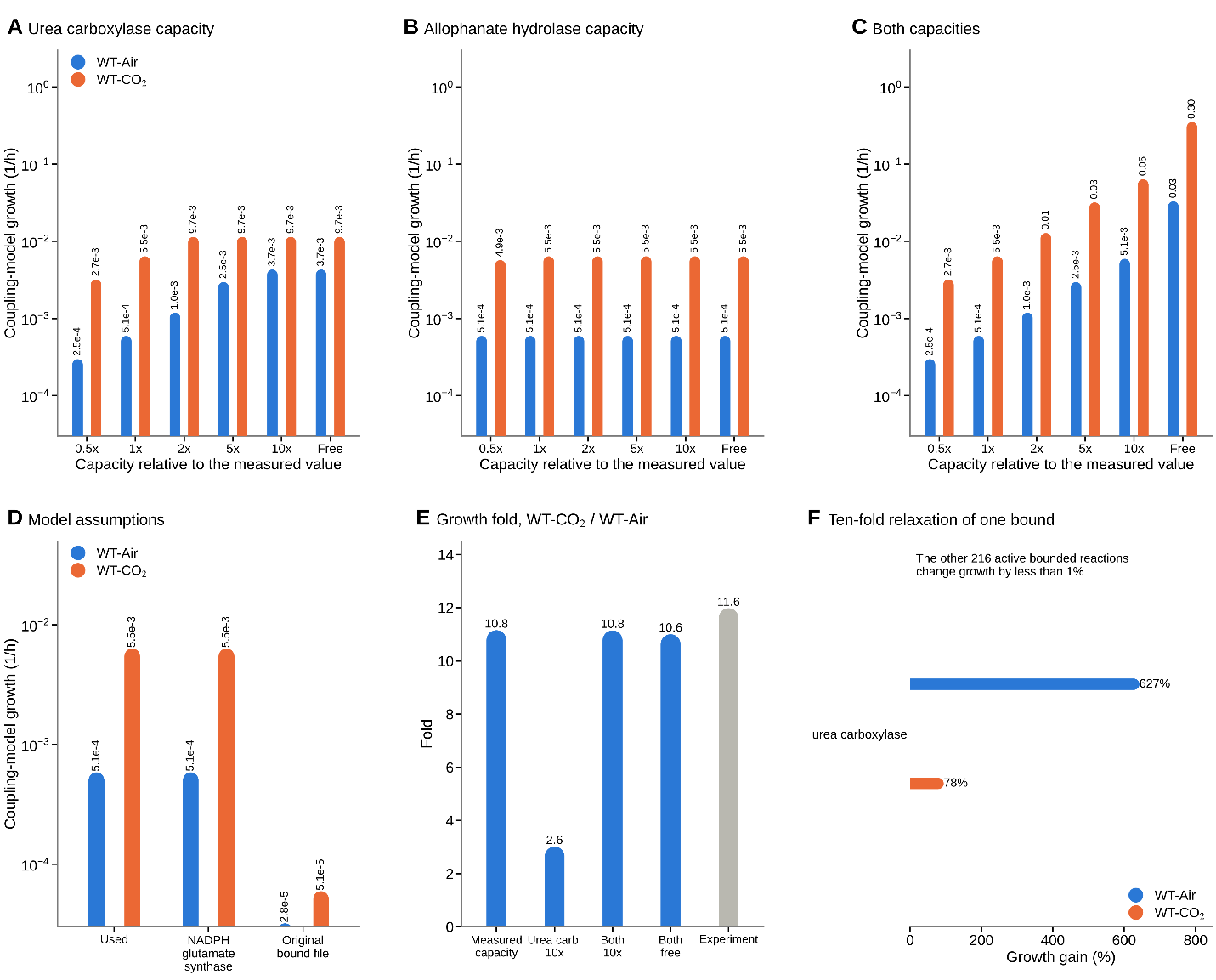

**Supplementary Figure S4. Sensitivity of the nitrogen and carbon coupling model.** (A) Coupling-model growth when the capacity of urea carboxylase is changed relative to the measured value. (B) The same for allophanate hydrolase. (C) The same for both enzymes together. (D) Effect of model assumptions, with an NADH or NADPH dependent glutamate synthase and with the original bound file. (E) Growth fold of WT-CO_2_ over WT-Air for several capacities, compared with the experiment. (F) Reactions whose ten-fold bound relaxation raises growth by more than 1%.

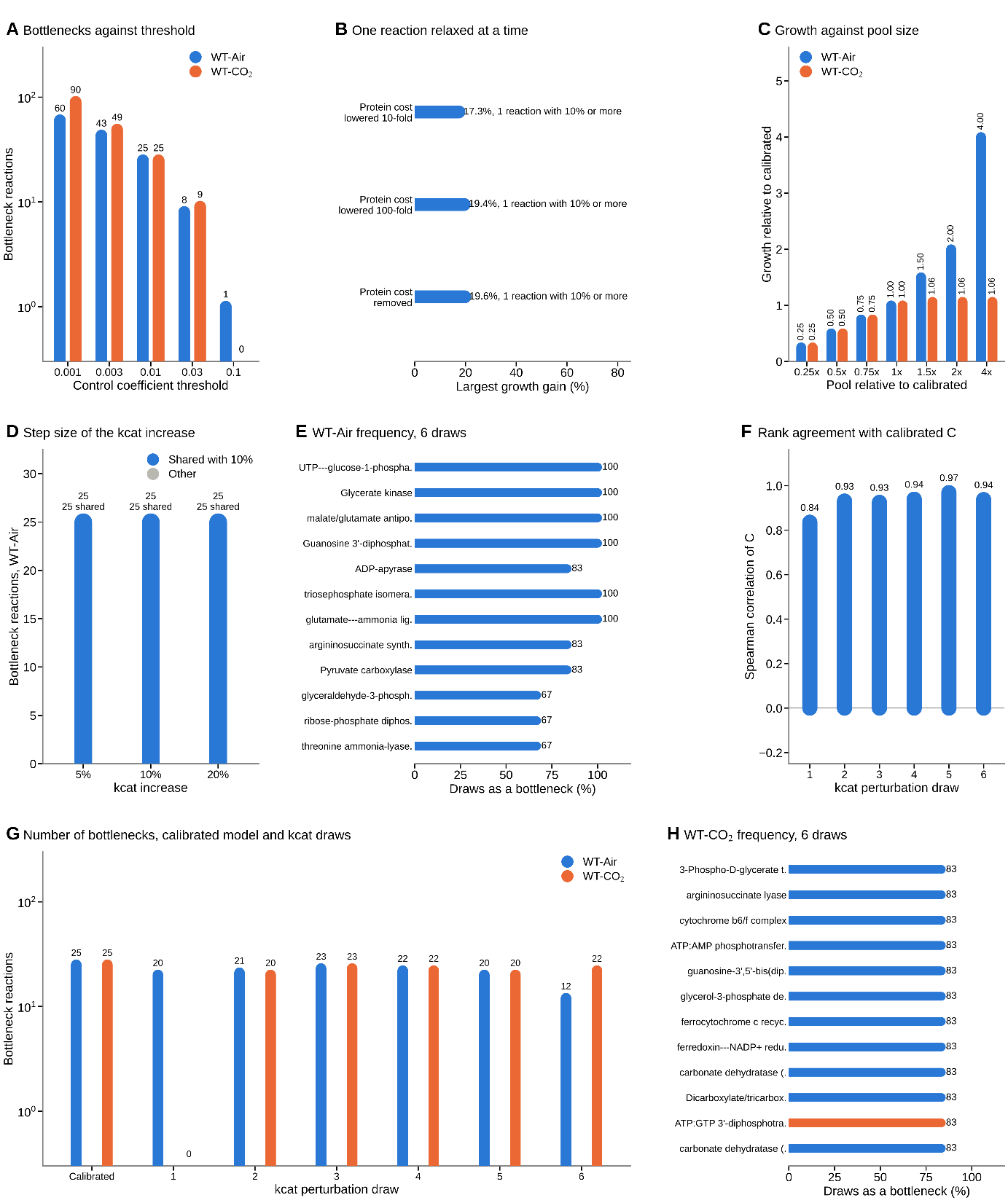

**Supplementary Figure S5. Sensitivity of the bottleneck analysis.** (A) Number of bottleneck reactions against the control coefficient threshold. (B) Largest growth gain when the protein cost of one reaction is lowered at a time in WT-Air. (C) Growth against the size of the protein pool, relative to the calibrated pool. (D) Bottleneck reactions in WT-Air for k_cat_ increases of 5%, 10% and 20%, and how many are shared with the 10% set. (E) Frequency of the WT-Air bottlenecks across six kcat perturbation draws. (F) Spearman correlation of the control coefficients of each draw with the calibrated ones, WT-Air. (G) Number of bottlenecks in the calibrated models and in every draw. (H) Frequency of the WT-CO_2_ bottlenecks across six draws.
