## Supplementary material for "Enzyme-Constrained Metabolic Modeling Reveals CO_2_-Dependent Growth Control in *Picochlorum celeri*": The corrections therefore transfer only partly in both directions, and the calibration is a partial test and not a validation

**Table S1** Biomass composition of iPiccel952 before and after curation. Coefficients in mmol/gDW and masses in mg per mmol of biomass.

| **Class** | **Metabolite ID** | **Name** | **Original coefficient** | **Curated coefficient** | **Original mass (mg)** | **Curated mass (mg)** |
| --- | --- | --- | --- | --- | --- | --- |
| Protein (amino acids) | glu__L_c | L-Glutamate | -0.122168 | -0.497801 | 17.975 | 73.241 |
| Protein (amino acids) | leu__L_c | L-Leucine | -0.104537 | -0.425958 | 13.712 | 55.874 |
| Protein (amino acids) | asp__L_c | L-Aspartate | -0.096538 | -0.393366 | 12.850 | 52.358 |
| Protein (amino acids) | asn__L_c | L-Asparagine | -0.096538 | -0.393366 | 12.754 | 51.971 |
| Protein (amino acids) | ala__L_c | L-Alanine | -0.135458 | -0.551952 | 12.068 | 49.175 |
| Protein (amino acids) | lys__L_c | L-Lysine | -0.061147 | -0.249156 | 8.939 | 36.424 |
| Protein (amino acids) | gly_c | Glycine | -0.118090 | -0.481182 | 8.865 | 36.121 |
| Protein (amino acids) | val__L_c | L-Valine | -0.077640 | -0.294415 | 9.095 | 34.490 |
| Protein (amino acids) | ser__L_c | L-Serine | -0.072732 | -0.296362 | 7.644 | 31.145 |
| Protein (amino acids) | phe__L_c | L-Phenylalanine | -0.045961 | -0.187278 | 7.592 | 30.936 |
| Protein (amino acids) | thr__L_c | L-Threonine | -0.061369 | -0.250060 | 7.310 | 29.787 |
| Protein (amino acids) | pro__L_c | L-Proline | -0.062612 | -0.255128 | 7.209 | 29.373 |
| Protein (amino acids) | tyr__L_c | L-Tyrosine | -0.034773 | -0.141689 | 6.300 | 25.672 |
| Protein (amino acids) | gln__L_c | L-Glutamine | -0.041309 | -0.168324 | 6.037 | 24.600 |
| Protein (amino acids) | ile__L_c | L-Isoleucine | -0.043412 | -0.176891 | 5.694 | 23.203 |
| Protein (amino acids) | his__L_c | L-Histidine | -0.018079 | -0.073667 | 2.805 | 11.430 |
| Protein (amino acids) | met__L_c | L-Methionine | -0.023720 | -0.071445 | 3.539 | 10.660 |
| Protein (amino acids) | trp__L_c | L-Tryptophan | -0.012536 | -0.051082 | 2.560 | 10.432 |
| Protein (amino acids) | cys__L_c | L-Cysteine | -0.019060 | -0.077666 | 2.309 | 9.410 |
| Protein (amino acids) | arg__L_c | L-Arginine | -0.001102 | -0.004490 | 0.192 | 0.782 |
| **Protein (amino acids)** | **Class total** | **20 species** |  |  | **155.4** | **627.1** |
| RNA (NTPs) | utp_c | UTP | -0.058616 | -0.033923 | 28.378 | 16.423 |
| RNA (NTPs) | gtp_c | GTP | -0.104206 | -0.028897 | 54.518 | 15.118 |
| RNA (NTPs) | ctp_c | CTP | -0.104206 | -0.028897 | 50.348 | 13.962 |
| **RNA (NTPs)** | **Class total** | **3 species** |  |  | **133.2** | **45.5** |
| DNA (dNTPs) | datp_c | dATP | -0.002180 | -0.003864 | 1.071 | 1.898 |
| DNA (dNTPs) | dttp_c | dTTP | -0.002180 | -0.003864 | 1.051 | 1.863 |
| DNA (dNTPs) | dgtp_c | dGTP | -0.003875 | -0.003291 | 1.965 | 1.669 |
| DNA (dNTPs) | dctp_c | dCTP | -0.003875 | -0.003291 | 1.810 | 1.538 |
| **DNA (dNTPs)** | **Class total** | **4 species** |  |  | **5.9** | **7.0** |
| Carbohydrate | man_c | D-Mannose | -0.109065 | -0.312533 | 19.649 | 56.305 |
| Carbohydrate | gal_c | D-Galactose | -0.160831 | -0.236511 | 28.975 | 42.609 |
| Carbohydrate | starch300_h | starch n=300 repeat units (80 repeat units amylose, 220 repeat units amylopectin, corresponds to maize starch) | -0.146004 | -0.000370 | 49.977 | 17.915 |
| Carbohydrate | rib-D_c | D-Ribose | 0.000000 | -0.081090 | - | 12.174 |
| Carbohydrate | arab__L_c | L-Arabinose | -0.057594 | 0.000000 | 8.647 | - |
| Carbohydrate | glc-A_c | alpha-D-glucose | -1.427343 | 0.000000 | 257.144 | - |
| **Carbohydrate** | **Class total** | **6 species** |  |  | **364.4** | **129.0** |
| Chlorophyll | chla_u | Chlorophyll a | -0.010084 | -0.045147 | 9.010 | 40.339 |
| Chlorophyll | chlb_u | Chlorophyll b | -0.016547 | -0.014817 | 15.016 | 13.446 |
| **Chlorophyll** | **Class total** | **2 species** |  |  | **24.0** | **53.8** |
| Carotenoid and xanthophyll | caro_u | beta-Carotene | -0.001412 | -0.001460 | 0.758 | 0.784 |
| Carotenoid and xanthophyll | lut_u | Lutein | -0.001260 | -0.001303 | 0.717 | 0.742 |
| Carotenoid and xanthophyll | loroxan_u | Loroxanthin | -0.000655 | -0.000678 | 0.383 | 0.396 |
| Carotenoid and xanthophyll | neoxan_u | Neoxanthin | -0.000555 | -0.000574 | 0.333 | 0.345 |
| Carotenoid and xanthophyll | acaro_h | alpha-Carotene | -0.000504 | -0.000521 | 0.271 | 0.280 |
| Carotenoid and xanthophyll | vioxan_u | Violaxanthin | -0.000353 | -0.000365 | 0.212 | 0.219 |
| Carotenoid and xanthophyll | zaxan_u | Zeaxanthin | -0.000303 | -0.000313 | 0.172 | 0.178 |
| Carotenoid and xanthophyll | anxan_u | Antheraxanthin | -0.000101 | -0.000104 | 0.059 | 0.061 |
| **Carotenoid and xanthophyll** | **Class total** | **8 species** |  |  | **2.9** | **3.0** |
| Lipid | pchol1601826Z9Z_c | pchol1601826Z9Z[c] | -0.066979 | -0.009402 | 50.774 | 7.127 |
| Lipid | pchol2226Z9Z1433Z6Z9Z_c | pchol2226Z9Z1433Z6Z9Z[c] | -0.054272 | -0.007618 | 42.336 | 5.943 |
| Lipid | 12dgr1601826Z9Z_h | 12dgr1601826Z9Z[h] | -0.064287 | -0.009024 | 38.118 | 5.351 |
| Lipid | pail16018212Z15Z_h | pail16018212Z15Z[h] | -0.042799 | -0.006008 | 35.739 | 5.017 |
| Lipid | tag1419Z1123Z6Z1123Z6Z_h | Triacylglycerol (14:1(9Z)/11:2(3Z,6Z)/11:2(3Z,6Z)) | -0.050705 | -0.007117 | 31.890 | 4.476 |
| Lipid | pchol1839Z12Z15Z1829Z12Z_c | pchol1839Z12Z15Z1829Z12Z[c] | -0.038537 | -0.005409 | 30.061 | 4.220 |
| Lipid | mgdg1836Z9Z12Z1636Z9Z12Z_h | mgdg1836Z9Z12Z1636Z9Z12Z[h] | -0.040223 | -0.005646 | 30.047 | 4.218 |
| Lipid | tag401001133Z6Z9Z_h | Triacylglycerol (4:0/10:0/11:3(3Z6Z9Z)) | -0.061929 | -0.008693 | 29.643 | 4.161 |
| Lipid | pchol1829Z12Z1826Z9Z_c | pchol1829Z12Z1826Z9Z[c] | -0.030781 | -0.004321 | 24.073 | 3.379 |
| Lipid | pchol1601836Z9Z12Z_c | pchol1601836Z9Z12Z[c] | -0.029551 | -0.004148 | 22.342 | 3.136 |
| Lipid | tag1419Z901123Z6Z_h | Triacylglycerol (14:1(9Z)/9:0/11:2(3Z,6Z)) | -0.029881 | -0.004194 | 18.075 | 2.537 |
| Lipid | mgdg1601826Z9Z_h | mgdg1601826Z9Z[h] | -0.023256 | -0.003264 | 17.560 | 2.465 |
| Lipid | pchol816Z1016Z_c | pchol816Z1016Z[c] | -0.030526 | -0.004285 | 16.289 | 2.287 |
| Lipid | mgdg1839Z12Z15Z1629Z12Z_h | mgdg1839Z12Z15Z1629Z12Z[h] | -0.017223 | -0.002418 | 12.900 | 1.811 |
| Lipid | pchol1839Z12Z15Z1839Z12Z15Z_c | pchol1839Z12Z15Z1839Z12Z15Z[c] | -0.016417 | -0.002304 | 12.773 | 1.793 |
| Lipid | tag1601601829Z12Z_h | Triacylglycerol (16:0/16:0/18:2(9Z,12Z)) | -0.015051 | -0.002113 | 12.513 | 1.756 |
| Lipid | tag1601119Z1123Z6Z_h | Triacylglycerol (16:0/11:1(9Z)/11:2(3Z,6Z)) | -0.018652 | -0.002618 | 12.329 | 1.731 |
| Lipid | pchol1829Z12Z1829Z12Z_c | pchol1829Z12Z1829Z12Z[c] | -0.015623 | -0.002193 | 12.218 | 1.715 |
| Lipid | 12dgr1619Z1829Z12Z_h | 12dgr1619Z1829Z12Z[h] | -0.020485 | -0.002875 | 12.105 | 1.699 |
| Lipid | pg1619Z1829Z12Z_h | pg1619Z1829Z12Z[h] | -0.015821 | -0.002221 | 11.786 | 1.654 |
| Lipid | 12dgr1616Z1839Z12Z15Z_h | 12dgr1616Z1839Z12Z15Z[h] | -0.019352 | -0.002716 | 11.396 | 1.600 |
| Lipid | pail190100_h | pail190100[h] | -0.013586 | -0.001907 | 10.447 | 1.466 |
| Lipid | mgdg1829Z12Z1626Z9Z_h | mgdg1829Z12Z1626Z9Z[h] | -0.013828 | -0.001941 | 10.385 | 1.458 |
| Lipid | pchol1601819Z_c | Phosphatidylcholine (16:0/18:1(9Z)) | -0.012873 | -0.001807 | 9.784 | 1.373 |
| Lipid | tag2901123Z6Z1233Z6Z9Z_h | Triacylglycerol (29:0/11:2(3Z6Z)/12:3(3Z6Z9Z)) | -0.011414 | -0.001602 | 9.740 | 1.367 |
| Lipid | pg1619Z1839Z12Z15Z_h | pg1619Z1839Z12Z15Z[h] | -0.012013 | -0.001686 | 8.925 | 1.253 |
| Lipid | tag1219Z801426Z9Z_h | Triacylglycerol (12:1(9Z)/8:0/14:2(6Z,9Z)) | -0.014377 | -0.002018 | 8.697 | 1.221 |
| Lipid | pg160160_h | pg160160[h] | -0.011845 | -0.001663 | 8.564 | 1.202 |
| Lipid | tag1216Z1101126Z9Z_h | Triacylglycerol (12:1(6Z)/11:0/11:2(6Z,9Z)) | -0.013971 | -0.001961 | 8.451 | 1.186 |
| Lipid | 12dgr1601836Z9Z12Z_h | 12dgr1601836Z9Z12Z[h] | -0.014302 | -0.002008 | 8.451 | 1.186 |
| Lipid | pe1702256Z9Z12Z15Z18Z_h | pe1702256Z9Z12Z15Z18Z[h] | -0.010673 | -0.001498 | 8.326 | 1.169 |
| Lipid | pchol1719Z1719Z_c | pchol1719Z1719Z[c] | -0.010783 | -0.001514 | 8.045 | 1.146 |
| Lipid | pail2219Z1829Z12Z_h | pail2219Z1829Z12Z[h] | -0.008735 | -0.001226 | 8.011 | 1.125 |
| Lipid | mgdg1601839Z12Z15Z_h | mgdg1601839Z12Z15Z[h] | -0.010453 | -0.001467 | 7.872 | 1.105 |
| Lipid | tag1419Z1101133Z6Z9Z_h | Triacylglycerol (14:1(9Z)/11:0/11:3(3Z,6Z,9Z)) | -0.012191 | -0.001711 | 7.691 | 1.080 |
| Lipid | tag2819Z1216Z1233Z6Z9Z_h | Triacylglycerol (28:1(9Z)/12:1(6Z)/11:3(3Z6Z9Z)) | -0.008372 | -0.001175 | 7.161 | 1.005 |
| Lipid | crm170250(O)_r | crm170250(O)[r] | -0.010040 | -0.001409 | 6.708 | 0.942 |
| Lipid | pchol2619Z80_c | pchol2619Z80[c] | -0.008453 | -0.001187 | 6.425 | 0.902 |
| Lipid | mgdg1601819Z_h | mgdg1601819Z[h] | -0.008306 | -0.001166 | 6.288 | 0.883 |
| Lipid | tag1601601836Z9Z12Z_h | Triacylglycerol (16:0/16:0/18:3(6Z,9Z,12Z)) | -0.007450 | -0.001046 | 6.179 | 0.867 |
| Lipid | pail1601836Z9Z12Z_h | pail1601836Z9Z12Z[h] | -0.007415 | -0.001041 | 6.177 | 0.867 |
| Lipid | pchol813Z1023Z6Z_c | pchol813Z1023Z6Z[c] | -0.011288 | -0.001584 | 6.001 | 0.842 |
| Lipid | pchol1801433Z6Z9Z_c | pchol1801433Z6Z9Z[c] | -0.008185 | -0.001149 | 5.958 | 0.836 |
| Lipid | tag2801226Z9Z1226Z9Z_h | Triacylglycerol (28:0/12:2(6Z9Z)/12:2(6Z9Z)) | -0.006778 | -0.000951 | 5.812 | 0.816 |
| Lipid | 12dgr1601819Z_h | 1,2-Diacyl-sn-glycerol (1-hexadecanoyl,2-(9Z)-octadecenoyl, 16:0/18:1(9Z)) | -0.009590 | -0.001346 | 5.706 | 0.801 |
| Lipid | pail1839Z12Z15Z2219Z_h | pail1839Z12Z15Z2219Z[h] | -0.006228 | -0.000874 | 5.700 | 0.800 |
| Lipid | tag2046Z9Z12Z15Z1123Z6Z2319Z_h | Triacylglycerol (20:4(6Z9Z12Z15Z)/11:2(3Z6Z)/23:1(9Z)) | -0.006464 | -0.000907 | 5.620 | 0.796 |
| Lipid | pchol1801836Z9Z12Z_c | pchol1801836Z9Z12Z[c] | -0.007102 | -0.000997 | 5.568 | 0.782 |
| Lipid | pg1719Z1829Z12Z_h | pg1719Z1829Z12Z[h] | -0.006907 | -0.000969 | 5.242 | 0.736 |
| Lipid | tag2801223Z6Z1226Z9Z_h | Triacylglycerol (28:0/12:2(3Z6Z)/12:2(6Z9Z)) | -0.006085 | -0.000854 | 5.217 | 0.732 |
| Lipid | pail2201839Z12Z15Z_h | pail2201839Z12Z15Z[h] | -0.005685 | -0.000798 | 5.215 | 0.732 |
| Lipid | tag40801133Z6Z9Z_h | Triacylglycerol (4:0/8:0/11:33Z6Z9Z) | -0.011521 | -0.001617 | 5.191 | 0.729 |
| Lipid | tag2053Z6Z9Z12Z15Z1123Z6Z2319Z_h | Triacylglycerol (20:5(3Z6Z9Z12Z15Z)/11:2(3Z6Z)/23:1(9Z)) | -0.005723 | -0.000803 | 5.032 | 0.706 |
| Lipid | pg1601619Z_h | pg1601619Z[h] | -0.006978 | -0.000979 | 5.030 | 0.706 |
| Lipid | tag1619Z1619Z2046Z9Z12Z15Z_h | Triacylglycerol (16:1(9Z)/16:1(9Z)/20:4(6Z9Z12Z15Z)) | -0.005892 | -0.000827 | 5.016 | 0.704 |
| Lipid | 12dgr180160_h | 12dgr180160[h] | -0.008260 | -0.001159 | 4.931 | 0.692 |
| Lipid | tag1216Z1123Z6Z1123Z6Z_h | Triacylglycerol (12:1(6Z)/11:2(3Z,6Z)/11:2(3Z,6Z)) | -0.007916 | -0.001111 | 4.757 | 0.668 |
| Lipid | tag2819Z1123Z6Z1123Z6Z_h | Triacylglycerol (28:1(9Z)/11:2(3Z6Z)/11:2(3Z6Z)) | -0.005480 | -0.000769 | 4.534 | 0.636 |
| Lipid | 12dgr1829Z12Z1839Z12Z15Z_h | 12dgr1829Z12Z1839Z12Z15Z[h] | -0.007325 | -0.001028 | 4.504 | 0.632 |
| Lipid | pchol816Z1019Z_c | pchol816Z1019Z[c] | -0.008409 | -0.001180 | 4.487 | 0.630 |
| Lipid | pchol2229Z12Z1433Z6Z9Z_c | pchol2229Z12Z1433Z6Z9Z[c] | -0.005648 | -0.000793 | 4.405 | 0.618 |
| Lipid | tag401123Z6Z1233Z6Z9Z_h | Triacylglycerol (4:0/11:2(3Z6Z)/12:3(3Z6Z9Z)) | -0.008603 | -0.001208 | 4.324 | 0.607 |
| Lipid | tag1901123Z6Z2263Z6Z9Z12Z15Z18Z_h | Triacylglycerol (19:0/11:2(3Z6Z)/22:6(3Z6Z9Z12Z15Z18Z) | -0.004811 | -0.000675 | 4.076 | 0.572 |
| Lipid | tag1829Z12Z1302119Z_h | Triacylglycerol (18:2(9Z,12Z)/13:0/21:1(9Z)) | -0.004718 | -0.000662 | 4.045 | 0.568 |
| Lipid | crm160160_r | crm160160[r] | -0.007855 | -0.001103 | 4.021 | 0.564 |
| Lipid | pail1619Z2246Z9Z12Z15Z_h | pail1619Z2246Z9Z12Z15Z[h] | -0.004461 | -0.000626 | 3.948 | 0.554 |
| Lipid | pchol18119Z1829Z12Z_c | pchol18119Z1829Z12Z[c] | -0.004912 | -0.000690 | 3.852 | 0.541 |
| Lipid | tag1106060_h | Triacylglycerol (11:0/6:0/6:0) | -0.008405 | -0.001180 | 3.838 | 0.539 |
| Lipid | pchol10060_c | pchol10060[c] | -0.007392 | -0.001038 | 3.767 | 0.529 |
| Lipid | tag28112Z1123Z6Z1123Z6Z_h | Triacylglycerol (28:1(12Z)/11:2(3Z6Z)/11:2(3Z6Z)) | -0.004552 | -0.000639 | 3.766 | 0.529 |
| Lipid | tag401001123Z6Z_h | Triacylglycerol (4:0/10:0/11:2(3Z6Z)) | -0.006814 | -0.000956 | 3.275 | 0.460 |
| Lipid | pg1601819Z_h | pg1601819Z[h] | -0.004051 | -0.000569 | 3.034 | 0.426 |
| Lipid | mgdg1839Z12Z15Z1839Z12Z15Z_h | mgdg1839Z12Z15Z1839Z12Z15Z[h] | -0.003758 | -0.000527 | 2.912 | 0.409 |
| Lipid | pchol1801829Z12Z_c | pchol1801829Z12Z_c | -0.003696 | -0.000519 | 2.906 | 0.408 |
| Lipid | tag1601836Z9Z12Z1836Z9Z12Z_h | Triacylglycerol (16:0/18:3(6Z9Z12Z)/18:3(6Z9Z12Z)) | -0.003359 | -0.000472 | 2.860 | 0.401 |
| Lipid | pail1601819Z_h | pail1601819Z[h] | -0.003371 | -0.000473 | 2.822 | 0.396 |
| Lipid | pe2301436Z9Z12Z_h | pe2301436Z9Z12Z[h] | -0.003714 | -0.000521 | 2.808 | 0.394 |
| Lipid | 12dgr1601839Z12Z15Z_h | 12dgr1601839Z12Z15Z[h] | -0.004748 | -0.000666 | 2.805 | 0.394 |
| Lipid | 12dgr160160_h | 12dgr160160[h] | -0.004843 | -0.000680 | 2.755 | 0.387 |
| Lipid | pail1839Z12Z15Z2249Z12Z15Z18Z_h | pail1839Z12Z15Z2249Z12Z15Z18Z[h] | -0.002981 | -0.000418 | 2.710 | 0.380 |
| Lipid | pg1619Z1819Z_h | pg1619Z1819Z[h] | -0.003610 | -0.000507 | 2.697 | 0.379 |
| Lipid | tag1501116Z240_h | Triacylglycerol (15:0/11:1(6Z)/24:0) | -0.003736 | -0.000524 | 2.694 | 0.437 |
| Lipid | tag2901023Z6Z1123Z6Z_h | Triacylglycerol (29:0/10:2(3Z6Z)/11:2(3Z6Z)) | -0.003143 | -0.000441 | 2.607 | 0.366 |
| Lipid | mgdg1829Z12Z1839Z12Z15Z_h | mgdg1829Z12Z1839Z12Z15Z[h] | -0.003348 | -0.000470 | 2.602 | 0.365 |
| Lipid | pchol2419Z1216Z_c | pchol2419Z1216Z[c] | -0.003259 | -0.000457 | 2.562 | 0.360 |
| Lipid | tag1201123Z6Z1123Z6Z_h | Triacylglycerol (12:0/11:2(3Z,6Z)/11:2(3Z,6Z)) | -0.004035 | -0.000566 | 2.433 | 0.341 |
| Lipid | tag1201126Z9Z1126Z9Z_h | Triacylglycerol (12:0/11:2(6Z,9Z)/11:2(6Z,9Z)) | -0.004028 | -0.000565 | 2.428 | 0.341 |
| Lipid | pe1919Z2039Z12Z15Z_h | pe1919Z2039Z12Z15Z[h] | -0.002851 | -0.000400 | 2.230 | 0.313 |
| Lipid | pg1601829Z12Z_h | pg1601829Z12Z[h] | -0.002907 | -0.000408 | 2.172 | 0.305 |
| Lipid | pchol1826Z9Z1826Z9Z_c | pchol1826Z9Z1826Z9Z[c] | -0.002759 | -0.000387 | 2.158 | 0.303 |
| Lipid | crm200160_r | crm200160[r] | -0.003484 | -0.000489 | 1.979 | 0.278 |
| Lipid | tag290801123Z6Z_h | Triacylglycerol (29:0/8:0/11:2(3Z6Z)) | -0.002425 | -0.000340 | 1.948 | 0.273 |
| Lipid | tag180160160_h | Triacylglycerol (18:0/16:0/16:0) | -0.002195 | -0.000308 | 1.834 | 0.257 |
| Lipid | 12dgr1819Z1829Z12Z_h | 12dgr1819Z1829Z12Z[h] | -0.002920 | -0.000410 | 1.807 | 0.254 |
| Lipid | pail1819Z2219Z_h | pail1819Z2219Z[h] | -0.001944 | -0.000273 | 1.787 | 0.251 |
| Lipid | 12dgr1839Z12Z15Z1839Z12Z15Z_h | 12dgr1839Z12Z15Z1839Z12Z15Z[h] | -0.002842 | -0.000399 | 1.742 | 0.245 |
| Lipid | mgdg1619Z1839Z12Z15Z_h | mgdg1619Z1839Z12Z15Z[h] | -0.002309 | -0.000324 | 1.734 | 0.243 |
| Lipid | tag40801123Z6Z_h | Triacylglycerol (4:0/8:0/11:23Z6Z) | -0.003680 | -0.000516 | 1.665 | 0.234 |
| Lipid | tag1401001233Z6Z9Z_h | Triacylglycerol (14:0/10:0/12:3(3Z,6Z,9Z)) | -0.003179 | -0.000446 | 1.656 | 0.282 |
| Lipid | tag1601602256Z9Z12Z15Z18Z_h | Triacylglycerol (16:0/16:0/22:5(6Z,9Z,12Z,15Z,18Z)) | -0.001828 | -0.000257 | 1.630 | 0.229 |
| Lipid | tag1216Z1223Z6Z1436Z9Z12Z_h | Triacylglycerol (12:1(6Z)/12:2(3Z,6Z)/14:3(6Z,9Z,12Z)) | -0.002469 | -0.000347 | 1.617 | 0.227 |
| Lipid | pchol1419Z2119Z_c | pchol1419Z2119Z[c] | -0.002126 | -0.000298 | 1.616 | 0.230 |
| Lipid | pchol1402046Z9Z12Z15Z_c | pchol1402046Z9Z12Z15Z[c] | -0.002134 | -0.000300 | 1.609 | 0.226 |
| Lipid | tag180160180_h | Triacylglycerol (18:0/16:0/18:0) | -0.001798 | -0.000252 | 1.553 | 0.218 |
| Lipid | mgdg1619Z1829Z12Z_h | mgdg1619Z1829Z12Z[h] | -0.002021 | -0.000284 | 1.522 | 0.214 |
| Lipid | tag3001226Z9Z1226Z9Z_h | Triacylglycerol (30:0/12:2(6Z9Z)/12:2(6Z9Z)) | -0.001683 | -0.000236 | 1.490 | 0.209 |
| Lipid | pchol2119Z1119Z_c | pchol2119Z1119Z[c] | -0.002037 | -0.000286 | 1.462 | 0.208 |
| Lipid | tag402419Z2419Z_h | Triacylglycerol (4:0/24:19Z/24:19Z) | -0.001643 | -0.000231 | 1.416 | 0.199 |
| Lipid | pe1801829Z12Z_c | phosphatidylethanolamine (18:0/18:2(9Z,12Z)) | -0.001824 | -0.000256 | 1.357 | 0.191 |
| Lipid | tag402302319Z_h | Triacylglycerol (4:0/23:0/23:19Z) | -0.001600 | -0.000225 | 1.336 | 0.188 |
| Lipid | 12dgr1619Z1839Z12Z15Z_h | 12dgr1619Z1839Z12Z15Z[h] | -0.002233 | -0.000313 | 1.315 | 0.185 |
| Lipid | tag2801223Z6Z1223Z6Z_h | Triacylglycerol (28:0/12:2(3Z6Z)/12:2(3Z6Z)) | -0.001502 | -0.000211 | 1.288 | 0.181 |
| Lipid | tag402036Z9Z12Z240_h | Triacylglycerol (4:0/20:36Z9Z12Z/24:0) | -0.001588 | -0.000223 | 1.276 | 0.179 |
| Lipid | tag1701419Z170_h | Triacylglycerol (17:0/14:1(9Z)/17:0) | -0.001559 | -0.000219 | 1.256 | 0.176 |
| Lipid | tag402219Z2319Z_h | Triacylglycerol (4:0/22:19Z/23:19Z) | -0.001505 | -0.000211 | 1.233 | 0.173 |
| Lipid | tag1101102419Z_h | Triacylglycerol (11:0/11:0/24:1(9Z)) | -0.001586 | -0.000223 | 1.233 | 0.173 |
| Lipid | pail1839Z12Z15Z2239Z12Z15Z_h | pail1839Z12Z15Z2239Z12Z15Z[h] | -0.001319 | -0.000185 | 1.202 | 0.169 |
| Lipid | tag1801433Z6Z9Z2246Z9Z12Z15Z_h | Triacylglycerol (18:0/14:3(3Z6Z9Z)/22:4(6Z9Z12Z15Z)) | -0.001333 | -0.000187 | 1.170 | 0.164 |
| Lipid | pchol819Z100_c | pchol819Z100[c] | -0.002153 | -0.000302 | 1.153 | 0.162 |
| Lipid | mgdg1601636Z9Z12Z_h | mgdg1601636Z9Z12Z[h] | -0.001581 | -0.000222 | 1.146 | 0.161 |
| Lipid | tag2901123Z6Z1219Z_h | Triacylglycerol (29:0/11:2(3Z6Z)/12:1(9Z)) | -0.001270 | -0.000178 | 1.089 | 0.153 |
| Lipid | mag1839Z12Z15Z_c | mag1839Z12Z15Z[c] | -0.003084 | -0.000433 | 1.087 | 0.153 |
| Lipid | tag2236Z9Z12Z1123Z6Z2119Z_h | Triacylglycerol (22:3(6Z9Z12Z)/11:2(3Z6Z)/21:1(9Z)) | -0.001433 | -0.000201 | 1.085 | 0.177 |
| Lipid | tag2901123Z6Z1419Z_h | Triacylglycerol (29:0/11:2(3Z6Z)/14:1(9Z)) | -0.001198 | -0.000168 | 1.063 | 0.149 |
| Lipid | tag40150150_h | Triacylglycerol (4:0/15:0/15:0) | -0.001691 | -0.000237 | 1.033 | 0.145 |
| Lipid | tag402256Z9Z12Z15Z18Z2419Z_h | Triacylglycerol (4:0/22:56Z9Z12Z15Z18Z/24:19Z) | -0.001248 | -0.000175 | 1.030 | 0.145 |
| Lipid | pail1601829Z12Z_h | pail1601829Z12Z[h] | -0.001213 | -0.000170 | 1.013 | 0.142 |
| Lipid | tag160160160_h | Triacylglycerol (16:0/16:0/16:0) | -0.001187 | -0.000167 | 0.958 | 0.134 |
| Lipid | pchol816Z1023Z6Z_c | pchol816Z1023Z6Z[c] | -0.001696 | -0.000238 | 0.902 | 0.127 |
| Lipid | tag2901123Z6Z1416Z_h | Triacylglycerol (29:0/11:2(3Z6Z)/14:1(6Z)) | -0.001003 | -0.000141 | 0.890 | 0.125 |
| Lipid | 12dgr1601829Z12Z_h | 12dgr1601829Z12Z[h] | -0.001494 | -0.000210 | 0.886 | 0.124 |
| Lipid | 12dgr1801829Z12Z_c | 12dgr1801829Z12Z[c] | -0.001393 | -0.000195 | 0.865 | 0.121 |
| Lipid | pail1601826Z9Z_h | pail1601826Z9Z[h] | -0.001031 | -0.000145 | 0.861 | 0.121 |
| Lipid | tag2901123Z6Z1133Z6Z9Z_h | Triacylglycerol (29:0/11:2(3Z6Z)/11:3(3Z6Z9Z)) | -0.001014 | -0.000142 | 0.853 | 0.120 |
| Lipid | tag2701123Z6Z1123Z6Z_h | Triacylglycerol (27:0/11:2(3Z6Z)/11:2(3Z6Z)) | -0.001021 | -0.000143 | 0.832 | 0.117 |
| Lipid | mgdg1601626Z9Z_h | mgdg1601626Z9Z[h] | -0.001074 | -0.000151 | 0.781 | 0.110 |
| Lipid | pchol1719Z1829Z12Z_c | pchol1719Z1829Z12Z[c] | -0.001021 | -0.000143 | 0.774 | 0.110 |
| Lipid | pe1801819Z_c | phosphatidylethanolamine (18:0/18:1(9Z)) | -0.001028 | -0.000144 | 0.767 | 0.108 |
| Lipid | pg1801416Z_h | pg1801416Z[h] | -0.001044 | -0.000147 | 0.753 | 0.106 |
| Lipid | pail1601823Z6Z_h | pail1601823Z6Z[h] | -0.000896 | -0.000126 | 0.748 | 0.105 |
| Lipid | tag1801126Z9Z2219Z_h | Triacylglycerol (18:0/11:2(6Z9Z)/22:1(9Z)) | -0.000877 | -0.000123 | 0.739 | 0.104 |
| Lipid | pchol1801839Z12Z15Z_c | pchol1801839Z12Z15Z[c] | -0.000939 | -0.000132 | 0.736 | 0.103 |
| Lipid | tag2256Z9Z12Z15Z18Z1123Z6Z2119Z_h | Triacylglycerol (22:5(6Z9Z12Z15Z18Z)/11:2(3Z6Z)/21:1(9Z)) | -0.000835 | -0.000117 | 0.734 | 0.103 |
| Lipid | tag401019Z1123Z6Z_h | Triacylglycerol (4:0/10:1(9Z)/11:2(3Z6Z)) | -0.001514 | -0.000213 | 0.725 | 0.102 |
| Lipid | tag402029Z12Z220_h | Triacylglycerol (4:0/20:29Z12Z/22:0) | -0.000927 | -0.000130 | 0.720 | 0.101 |
| Lipid | tag1801123Z6Z2219Z_h | Triacylglycerol (18:0/11:2(3Z6Z)/22:1(9Z)) | -0.000806 | -0.000113 | 0.680 | 0.095 |
| Lipid | pe2419Z1126Z9Z_h | pe2419Z1126Z9Z[h] | -0.000919 | -0.000129 | 0.669 | 0.094 |
| Lipid | pchol1601619Z_c | pchol1601619Z[c] | -0.000910 | -0.000128 | 0.666 | 0.093 |
| Lipid | pchol1801435Z8Z11Z_c | pchol1801435Z8Z11Z[c] | -0.000909 | -0.000128 | 0.662 | 0.093 |
| Lipid | pchol2119Z1219Z_c | pchol2119Z1219Z[c] | -0.000889 | -0.000125 | 0.650 | 0.093 |
| Lipid | tag16060230_h | Triacylglycerol (16:0/6:0/23:0) | -0.000842 | -0.000118 | 0.644 | 0.090 |
| Lipid | tag2301219Z1219Z_h | Triacylglycerol (23:0/12:1(9Z)/21:1(9Z)) | -0.000787 | -0.000111 | 0.625 | 0.088 |
| Lipid | tag2819Z1123Z6Z1419Z_h | Triacylglycerol (28:1(9Z)/11:2(3Z6Z)/14:1(9Z)) | -0.000717 | -0.000101 | 0.624 | 0.088 |
| Lipid | pe200200_h | pe200200[h] | -0.000771 | -0.000108 | 0.620 | 0.087 |
| Lipid | mgdg1819Z1829Z12Z_h | mgdg1819Z1829Z12Z[h] | -0.000786 | -0.000110 | 0.614 | 0.086 |
| Lipid | pchol2419Z1219Z_c | pchol2419Z1219Z[c] | -0.000774 | -0.000109 | 0.609 | 0.085 |
| Lipid | pe2056Z9Z12Z15Z18Z210_h | pe2056Z9Z12Z15Z18Z210[h] | -0.000751 | -0.000105 | 0.607 | 0.085 |
| Lipid | pe2719Z1219Z_h | pe2719Z1219Z[h] | -0.000771 | -0.000108 | 0.606 | 0.085 |
| Lipid | tag1846Z9Z12Z15Z1123Z6Z240_h | Triacylglycerol (18:4(6Z,9Z,12Z,15Z)/11:2(3Z6Z)/24:0) | -0.000701 | -0.000098 | 0.606 | 0.085 |
| Lipid | tag2001123Z6Z1719Z_h | Triacylglycerol (20:0/11:2(3Z6Z)/17:1(9Z)) | -0.000738 | -0.000104 | 0.588 | 0.083 |
| Lipid | crm180260(O)_r | crm180260(O)[r] | -0.000840 | -0.000118 | 0.585 | 0.082 |
| Lipid | pg1202239Z12Z15Z_h | pg1202239Z12Z15Z[h] | -0.000772 | -0.000108 | 0.575 | 0.081 |
| Lipid | tag2901123Z6Z140_h | Triacylglycerol (29:0/11:2(3Z6Z)/14:0) | -0.000646 | -0.000091 | 0.574 | 0.081 |
| Lipid | tag402002119Z_h | Triacylglycerol (4:0/20:0/21:19Z) | -0.000747 | -0.000105 | 0.572 | 0.080 |
| Lipid | tag1819Z1116Z1819Z_h | Triacylglycerol (18:1(9Z)/11:1(6Z)/18:1(9Z)) | -0.000705 | -0.000099 | 0.555 | 0.078 |
| Lipid | tag40901016Z_h | Triacylglycerol (4:0/9:0/10:16Z) | -0.001214 | -0.000170 | 0.552 | 0.077 |
| Lipid | crm200220(O)_r | crm200220(O)[r] | -0.000822 | -0.000115 | 0.549 | 0.077 |
| Lipid | tag2619Z1123Z6Z1123Z6Z_h | Triacylglycerol (26:1(9Z)/11:2(3Z6Z)/11:2(3Z6Z)) | -0.000689 | -0.000097 | 0.549 | 0.077 |
| Lipid | mgdg160160_h | mgdg160160[h] | -0.000748 | -0.000105 | 0.547 | 0.077 |
| Lipid | tag2429Z12Z1123Z6Z1419Z_h | Triacylglycerol (24:2(9Z12Z)/11:2(3Z6Z)/14:1(9Z)) | -0.000661 | -0.000093 | 0.543 | 0.076 |
| Lipid | mag160_c | 1-monoacylglycerol (16:0) | -0.001610 | -0.000226 | 0.532 | 0.075 |
| Lipid | pchol2119Z1226Z9Z_c | pchol2119Z1226Z9Z[c] | -0.000724 | -0.000102 | 0.529 | 0.075 |
| Lipid | tag1919Z1123Z6Z2053Z6Z9Z12Z15Z_h | Triacylglycerol (19:1(9Z)/11:2(3Z6Z)/20:5(3Z6Z9Z12Z15Z)) | -0.000633 | -0.000089 | 0.519 | 0.073 |
| Lipid | pe1502256Z9Z12Z15Z18Z_h | pe1502256Z9Z12Z15Z18Z[h] | -0.000685 | -0.000096 | 0.515 | 0.072 |
| Lipid | pchol2401433Z6Z9Z_c | pchol2401433Z6Z9Z[c] | -0.000633 | -0.000089 | 0.514 | 0.072 |
| Lipid | pchol1801426Z9Z_c | pchol1801426Z9Z[c] | -0.000702 | -0.000098 | 0.512 | 0.072 |
| Lipid | tag1419Z1116Z1123Z6Z_h | Triacylglycerol (14:1(9Z)/11:1(6Z)/11:2(3Z,6Z)) | -0.000808 | -0.000113 | 0.510 | 0.072 |
| Lipid | mag1829Z12Z_c | mag1829Z12Z[c] | -0.001404 | -0.000197 | 0.498 | 0.070 |
| Lipid | pg1616Z1839Z12Z15Z_h | pg1616Z1839Z12Z15Z[h] | -0.000669 | -0.000094 | 0.497 | 0.070 |
| Lipid | crm180260_r | crm180260[r] | -0.000727 | -0.000102 | 0.494 | 0.069 |
| Lipid | tag2501123Z6Z1123Z6Z_h | Triacylglycerol (25:0/11:2(3Z6Z)/11:2(3Z6Z)) | -0.000621 | -0.000087 | 0.488 | 0.068 |
| Lipid | tag1919Z1202119Z_h | Triacylglycerol (19:1(9Z)/12:0/21:1(9Z)) | -0.000561 | -0.000079 | 0.482 | 0.068 |
| Lipid | tag1626Z9Z1123Z6Z1819Z_h | Triacylglycerol (16:2(6Z9Z)/11:2(3Z6Z)/18:1(9Z)) | -0.000621 | -0.000087 | 0.469 | 0.066 |
| Lipid | tag1419Z1001133Z6Z9Z_h | Triacylglycerol (14:1(9Z)/10:0/11:3(3Z,6Z,9Z)) | -0.000928 | -0.000130 | 0.468 | 0.080 |
| Lipid | pchol1402119Z_c | pchol1402119Z[c] | -0.000605 | -0.000085 | 0.461 | 0.066 |
| Lipid | tag2501016Z170_h | Triacylglycerol (25:0/10:1(6Z)/17:0) | -0.000526 | -0.000074 | 0.453 | 0.064 |
| Lipid | tag2901123Z6Z1433Z6Z9Z_h | Triacylglycerol (29:0/11:2(3Z6Z)/14:3(3Z6Z9Z)) | -0.000510 | -0.000072 | 0.451 | 0.063 |
| Lipid | pchol1719Z2039Z12Z15Z_c | pchol1719Z2039Z12Z15Z[c] | -0.000570 | -0.000080 | 0.447 | 0.064 |
| Lipid | pg1801413Z_h | pg1801413Z[h] | -0.000603 | -0.000085 | 0.435 | 0.061 |
| Lipid | pg1839Z12Z15Z1839Z12Z15Z_h | pg1839Z12Z15Z1839Z12Z15Z[h] | -0.000562 | -0.000079 | 0.431 | 0.060 |
| Lipid | pchol1801819Z_c | pchol1801819Z_c | -0.000545 | -0.000076 | 0.429 | 0.060 |
| Lipid | tag2901123Z6Z1426Z9Z_h | Triacylglycerol (29:0/11:2(3Z6Z)/14:2(6Z9Z)) | -0.000476 | -0.000067 | 0.421 | 0.059 |
| Lipid | tag1619Z1123Z6Z220_h | Triacylglycerol (16:1(9Z)/11:2(3Z6Z)/22:0)) | -0.000513 | -0.000072 | 0.418 | 0.059 |
| Lipid | tag1619Z1123Z6Z190_h | Triacylglycerol (16:1(9Z)/11:2(3Z6Z)/19:0)) | -0.000537 | -0.000075 | 0.415 | 0.058 |
| Lipid | tag1839Z12Z15Z801123Z6Z_h | Triacylglycerol (18:3(9Z12Z15Z)/8:0/11:2(3Z6Z)) | -0.000622 | -0.000087 | 0.400 | 0.056 |
| Lipid | pg100220_h | pg100220[h] | -0.000541 | -0.000076 | 0.391 | 0.055 |
| Lipid | tag2819Z1123Z6Z150_h | Triacylglycerol (28:1(9Z)/11:2(3Z6Z)/15:0) | -0.000432 | -0.000061 | 0.389 | 0.054 |
| Lipid | pail160160_h | pail160160[h] | -0.000478 | -0.000067 | 0.387 | 0.054 |
| Lipid | tag2901016Z1123Z6Z_h | Triacylglycerol (29:0/10:1(6Z)/11:2(3Z6Z)) | -0.000465 | -0.000065 | 0.387 | 0.054 |
| Lipid | pg16018312Z15Z18Z_h | pg16018312Z15Z18Z[h] | -0.000512 | -0.000072 | 0.382 | 0.054 |
| Lipid | pchol1219Z60_c | pchol1219Z60[c] | -0.000708 | -0.000099 | 0.379 | 0.053 |
| Lipid | pchol1719Z160_c | pchol1719Z160[c] | -0.000516 | -0.000072 | 0.379 | 0.054 |
| Lipid | tag1839Z12Z15Z1123Z6Z2239Z12Z15Z_h | Triacylglycerol (18:3(9Z12Z15Z)/11:2(3Z6Z)/22:3(9Z,12Z,15Z)) | -0.000441 | -0.000062 | 0.374 | 0.052 |
| Lipid | pchol2419Z1433Z6Z9Z_c | pchol2419Z1433Z6Z9Z[c] | -0.000458 | -0.000064 | 0.371 | 0.052 |
| Lipid | tag2039Z12Z15Z1123Z6Z2263Z6Z9Z12Z15Z18Z_h | Triacylglycerol (20:3(9Z12Z15Z)/11:2(3Z6Z)/22:6(3Z6Z9Z12Z15Z18Z)) | -0.000431 | -0.000061 | 0.371 | 0.052 |
| Lipid | pchol180140_c | pchol180140[c] | -0.000501 | -0.000070 | 0.368 | 0.052 |
| Lipid | tag2901123Z6Z1126Z9Z_h | Triacylglycerol (29:0/11:2(3Z6Z)/11:2(6Z9Z)) | -0.000427 | -0.000060 | 0.360 | 0.051 |
| Lipid | tag801123Z6Z240_h | Triacylglycerol (8:0/11:23Z6Z/24:0) | -0.000487 | -0.000068 | 0.357 | 0.050 |
| Lipid | pg1719Z1716Z_h | pg1719Z1716Z[h] | -0.000473 | -0.000066 | 0.353 | 0.050 |
| Lipid | pchol2501233Z6Z9Z_c | pchol2501233Z6Z9Z[c] | -0.000437 | -0.000061 | 0.349 | 0.049 |
| Lipid | tag2901123Z6Z1123Z6Z_h | Triacylglycerol (29:0/11:2(3Z6Z)/11:2(3Z6Z)) | -0.000408 | -0.000057 | 0.344 | 0.048 |
| Lipid | pe1601819Z_h | pe1601819Z[h] | -0.000475 | -0.000067 | 0.341 | 0.048 |
| Lipid | tag3019Z1226Z9Z1226Z9Z_h | Triacylglycerol (30:1(9Z)/12:2(6Z9Z)/12:2(6Z9Z)) | -0.000383 | -0.000054 | 0.338 | 0.047 |
| Lipid | pe18111Z1819Z_c | phosphatidylethanolamine (18:1(11Z)/18:1(9Z)) | -0.000442 | -0.000062 | 0.329 | 0.046 |
| Lipid | whttdca_h | omega hydroxy tetradecanoate (n-C14:0) | -0.001339 | -0.000188 | 0.327 | 0.046 |
| Lipid | pg1202236Z9Z12Z_h | pg1202236Z9Z12Z[h] | -0.000430 | -0.000060 | 0.321 | 0.045 |
| Lipid | pg1839Z12Z15Z1829Z12Z_h | pg1839Z12Z15Z1829Z12Z[h] | -0.000408 | -0.000057 | 0.314 | 0.044 |
| Lipid | pe2039Z12Z15Z2056Z9Z12Z15Z18Z_h | pe2039Z12Z15Z2056Z9Z12Z15Z18Z[h] | -0.000397 | -0.000056 | 0.313 | 0.044 |
| Lipid | tag1219Z901123Z6Z_h | Triacylglycerol (12:1(9Z)/9:0/11:1(3Z,6Z)) | -0.000531 | -0.000075 | 0.307 | 0.043 |
| Lipid | ttdca_h | tetradecanoate (n-C14:0) | -0.001339 | -0.000188 | 0.306 | 0.043 |
| Lipid | tag401019Z2246Z9Z12Z15Z_h | Triacylglycerol (4:0/10:1(9Z)/22:4(6Z9Z12Z15Z)) | -0.000478 | -0.000067 | 0.301 | 0.042 |
| Lipid | tag2401123Z6Z1123Z6Z_h | Triacylglycerol (24:0/11:2(3Z6Z)/11:2(3Z6Z)) | -0.000388 | -0.000054 | 0.299 | 0.042 |
| Lipid | pg180160_h | pg180160[h] | -0.000396 | -0.000056 | 0.298 | 0.042 |
| Lipid | tag2719Z60110_h | Triacylglycerol (27:1(9Z)/6:0/11:0) | -0.000396 | -0.000056 | 0.295 | 0.041 |
| Lipid | pe1601829Z12Z_h | pe1601829Z12Z[h] | -0.000410 | -0.000058 | 0.294 | 0.041 |
| Lipid | pg1801419Z_h | pg1801419Z[h] | -0.000404 | -0.000057 | 0.291 | 0.041 |
| Lipid | tag401123Z6Z21112Z_h | Triacylglycerol (4:0/11:2(3Z6Z)/21:1(12Z)) | -0.000457 | -0.000064 | 0.289 | 0.041 |
| Lipid | tag40901123Z6Z_h | Triacylglycerol (4:0/9:0/11:23Z6Z) | -0.000616 | -0.000086 | 0.288 | 0.040 |
| Lipid | tag160901123Z6Z_h | Triacylglycerol (16:0/9:0/11:2(3Z6Z)) | -0.000452 | -0.000063 | 0.287 | 0.040 |
| Lipid | mgdg1829Z12Z140_h | mgdg1829Z12Z140[h] | -0.000389 | -0.000055 | 0.283 | 0.040 |
| Lipid | tag2501123Z6Z1233Z6Z9Z_h | Triacylglycerol (25:0/11:2(3Z6Z)/12:3(3Z6Z9Z)) | -0.000351 | -0.000049 | 0.280 | 0.039 |
| Lipid | tag401123Z6Z210_h | Triacylglycerol (4:0/11:2(3Z6Z)/21:0) | -0.000441 | -0.000062 | 0.280 | 0.039 |
| Lipid | tag170601123Z6Z_h | Triacylglycerol (17:0/6:0/11:2(3Z6Z)) | -0.000455 | -0.000064 | 0.276 | 0.039 |
| Lipid | tag2608090_h | Triacylglycerol (26:0/8:0/9:0) | -0.000356 | -0.000050 | 0.271 | 0.037 |
| Lipid | pail1802219Z_h | pail1802219Z[h] | -0.000294 | -0.000041 | 0.271 | 0.038 |
| Lipid | tag2601426Z9Z1433Z6Z9Z_h | Triacylglycerol (26:0/14:2(6Z9Z)/14:3(3Z6Z9Z)) | -0.000304 | -0.000043 | 0.263 | 0.038 |
| Lipid | crm180240_r | crm180240[r] | -0.000403 | -0.000057 | 0.263 | 0.037 |
| Lipid | tag1216Z1223Z6Z1223Z6Z_h | Triacylglycerol (12:1(6Z)/12:2(3Z,6Z)/12:2(3Z,6Z)) | -0.000412 | -0.000058 | 0.259 | 0.036 |
| Lipid | tag1216Z1101123Z6Z_h | Triacylglycerol (12:1(6Z)/11:0/11:2(3Z,6Z)) | -0.000426 | -0.000060 | 0.258 | 0.036 |
| Lipid | mgdg2801426Z9Z_h | mgdg2801426Z9Z[h] | -0.000290 | -0.000041 | 0.252 | 0.035 |
| Lipid | tag1419Z1223Z6Z1223Z6Z_h | Triacylglycerol (14:1(9Z)/12:2(3Z,6Z)/12:2(3Z,6Z)) | -0.000371 | -0.000052 | 0.244 | 0.034 |
| Lipid | tag2046Z9Z12Z15Z1302029Z12Z_h | Triacylglycerol (20:4(6Z9Z12Z15Z)/13:0/20:2(9Z,12Z)) | -0.000278 | -0.000039 | 0.242 | 0.034 |
| Lipid | tag2039Z12Z15Z1119Z2263Z6Z9Z12Z15Z18Z_h | Triacylglycerol (20:3(9Z12Z15Z)/11:1(9Z)/22:6(3Z6Z9Z12Z15Z18Z)) | -0.000281 | -0.000039 | 0.242 | 0.034 |
| Lipid | tag120601433Z6Z9Z_h | Triacylglycerol (12:0/6:0/14:3(3Z,6Z,9Z)) | -0.000408 | -0.000057 | 0.235 | 0.033 |
| Lipid | mgdg1839Z12Z15Z1636Z9Z12Z_h | mgdg1839Z12Z15Z1636Z9Z12Z[h] | -0.000313 | -0.000044 | 0.234 | 0.033 |
| Lipid | tag1419Z9090_h | Triacylglycerol (14:1(9Z)/9:0/9:0) | -0.000401 | -0.000056 | 0.233 | 0.033 |
| Lipid | tag160601123Z6Z_h | Triacylglycerol (16:0/6:0/11:2(3Z6Z)) | -0.000382 | -0.000054 | 0.227 | 0.032 |
| Lipid | pchol2019Z130_c | pchol2019Z130[c] | -0.000297 | -0.000042 | 0.218 | 0.031 |
| Lipid | tag12060160_h | Triacylglycerol (12:0/6:0/16:0) | -0.000351 | -0.000049 | 0.214 | 0.030 |
| Lipid | tag401001119Z_h | Triacylglycerol (4:0/10:0/11:1(9Z)) | -0.000437 | -0.000061 | 0.211 | 0.030 |
| Lipid | pg16112Z1839Z12Z15Z_h | pg16112Z1839Z12Z15Z[h] | -0.000284 | -0.000040 | 0.211 | 0.030 |
| Lipid | tag12060180_h | Triacylglycerol (12:0/6:0/18:0) | -0.000319 | -0.000045 | 0.204 | 0.029 |
| Lipid | tag1919Z1226Z9Z2029Z12Z_h | Triacylglycerol (19:1(9Z)/12:2(6Z9Z)/20:2(9Z,12Z)) | -0.000234 | -0.000033 | 0.197 | 0.028 |
| Lipid | tag2101219Z1219Z_h | Triacylglycerol (21:0/12:1(9Z)/12:1(9Z)) | -0.000255 | -0.000036 | 0.195 | 0.027 |
| Lipid | pg1719Z1719Z_h | pg1719Z1719Z[h] | -0.000259 | -0.000036 | 0.194 | 0.027 |
| Lipid | tag1101126Z9Z120_h | Triacylglycerol (11:0/11:2(6Z,9Z)/12:0) | -0.000313 | -0.000044 | 0.190 | 0.027 |
| Lipid | tag1213Z1101123Z6Z_h | Triacylglycerol (12:1(3Z)/11:0/11:2(3Z,6Z)) | -0.000308 | -0.000043 | 0.186 | 0.026 |
| Lipid | pchol1002239Z12Z15Z_c | pchol1002239Z12Z15Z[c] | -0.000256 | -0.000036 | 0.186 | 0.026 |
| Lipid | tag3001123Z6Z1123Z6Z_h | Triacylglycerol (30:0/11:2(3Z6Z)/11:2(3Z6Z)) | -0.000216 | -0.000030 | 0.185 | 0.026 |
| Lipid | tag1801123Z6Z150_h | Triacylglycerol (18:0/11:2(3Z6Z)/15:0) | -0.000238 | -0.000033 | 0.178 | 0.025 |
| Lipid | tag2519Z1023Z6Z1123Z6Z_h | Triacylglycerol (25:1(9Z)/10:2(3Z6Z)/11:2(3Z6Z)) | -0.000212 | -0.000030 | 0.178 | 0.023 |
| Lipid | pchol1802029Z12Z_c | pchol1802029Z12Z[c] | -0.000217 | -0.000030 | 0.176 | 0.025 |
| Lipid | crm2219Z1819Z(2O)_r | crm2219Z1819Z(2O)[r] | -0.000275 | -0.000039 | 0.175 | 0.025 |
| Lipid | tag801223Z6Z1419Z_h | Triacylglycerol (8:0/12:23Z6Z/14:19Z) | -0.000285 | -0.000040 | 0.173 | 0.024 |
| Lipid | tag801001829Z12Z_h | Triacylglycerol (8:0/10:0/18:29Z12Z) | -0.000267 | -0.000037 | 0.169 | 0.024 |
| Lipid | tag2801101426Z9Z_h | Triacylglycerol (28:0/11:0/14:2(6Z9Z)) | -0.000188 | -0.000026 | 0.167 | 0.023 |
| Lipid | tag2053Z6Z9Z12Z15Z1123Z6Z2219Z_h | Triacylglycerol (20:5(3Z6Z9Z12Z15Z)/11:2(3Z6Z)/22:1(9Z)) | -0.000192 | -0.000027 | 0.166 | 0.023 |
| Lipid | pail1601839Z12Z15Z_h | pail1601839Z12Z15Z[h] | -0.000198 | -0.000028 | 0.165 | 0.023 |
| Lipid | tag2619Z1119Z1123Z6Z_h | Triacylglycerol (26:1(9Z)/11:1(9Z)/11:2(3Z6Z)) | -0.000204 | -0.000029 | 0.163 | 0.023 |
| Lipid | tag2519Z1123Z6Z1219Z_h | Triacylglycerol (25:1(9Z)/11:2(3Z6Z)/12:1(9Z)) | -0.000203 | -0.000028 | 0.162 | 0.023 |
| Lipid | tag2819Z1001123Z6Z_h | Triacylglycerol (28:1(9Z)/10:0/11:2(3Z6Z)) | -0.000185 | -0.000026 | 0.151 | 0.021 |
| Lipid | tag120901119Z_h | Triacylglycerol (12:0/9:0/11:1(9Z)) | -0.000260 | -0.000036 | 0.151 | 0.021 |
| Lipid | pg1002419Z_h | pg1002419Z[h] | -0.000199 | -0.000028 | 0.149 | 0.021 |
| Lipid | tag110601023Z6Z_h | Triacylglycerol (11:0/6:0/10:2(3Z,6Z)) | -0.000281 | -0.000039 | 0.143 | 0.020 |
| Lipid | tag1601002429Z12Z_h | Triacylglycerol (16:0/10:0/24:2(9Z,12Z)) | -0.000191 | -0.000027 | 0.138 | 0.022 |
| Lipid | tag2039Z12Z15Z1123Z6Z2319Z_h | Triacylglycerol (20:3(9Z12Z15Z)/11:2(3Z6Z)/23:1(9Z)) | -0.000155 | -0.000022 | 0.135 | 0.019 |
| Lipid | tag60801123Z6Z_h | Triacylglycerol (6:0/8:0/11:23Z6Z) | -0.000279 | -0.000039 | 0.134 | 0.019 |
| Lipid | pchol2701133Z6Z9Z_c | pchol2701133Z6Z9Z[c] | -0.000159 | -0.000022 | 0.129 | 0.018 |
| Lipid | pchol2426Z9Z1433Z6Z9Z_c | pchol2426Z9Z1433Z6Z9Z[c] | -0.000155 | -0.000022 | 0.125 | 0.018 |
| Lipid | mgdg1601829Z12Z_h | mgdg1601829Z12Z[h] | -0.000162 | -0.000023 | 0.122 | 0.017 |
| Lipid | pe2046Z9Z12Z15Z2056Z9Z12Z15Z18Z_h | pe2046Z9Z12Z15Z2056Z9Z12Z15Z18Z[h] | -0.000151 | -0.000021 | 0.118 | 0.017 |
| Lipid | tag1101123Z6Z120_h | Triacylglycerol (11:0/11:2(3Z,6Z)/12:0) | -0.000193 | -0.000027 | 0.117 | 0.016 |
| Lipid | tag1819Z1123Z6Z1123Z6Z_h | Triacylglycerol (18:1(9Z)/11:2(3Z6Z)/11:2(3Z6Z)) | -0.000164 | -0.000023 | 0.112 | 0.016 |
| Lipid | pchol2001436Z9Z12Z_c | pchol2001436Z9Z12Z[c] | -0.000148 | -0.000021 | 0.112 | 0.016 |
| Lipid | tag401123Z6Z1219Z_h | Triacylglycerol (4:0/11:2(3Z6Z)/12:1(9Z)) | -0.000213 | -0.000030 | 0.108 | 0.015 |
| Lipid | tag1839Z12Z15Z601123Z6Z_h | Triacylglycerol (18:3(9Z12Z15Z)/6:0/11:2(3Z6Z)) | -0.000169 | -0.000024 | 0.104 | 0.015 |
| Lipid | tag401123Z6Z2119Z_h | Triacylglycerol (4:0/11:2(3Z6Z)/21:1(9Z)) | -0.000163 | -0.000023 | 0.103 | 0.015 |
| Lipid | tag601001829Z12Z_h | Triacylglycerol (6:0/10:0/18:29Z12Z) | -0.000169 | -0.000024 | 0.103 | 0.014 |
| Lipid | pchol1802053Z6Z9Z12Z15Z_c | pchol1802053Z6Z9Z12Z15Z[c] | -0.000125 | -0.000018 | 0.101 | 0.014 |
| Lipid | crm160180_r | crm160180[r] | -0.000178 | -0.000025 | 0.096 | 0.014 |
| Lipid | tag1501123Z6Z1226Z9Z_h | Triacylglycerol (15:0/11:2(3Z,6Z)/12:2(6Z,9Z) | -0.000171 | -0.000024 | 0.094 | 0.016 |
| Lipid | pchol2119Z1216Z_c | pchol2119Z1216Z[c] | -0.000126 | -0.000018 | 0.092 | 0.013 |
| Lipid | pail1839Z12Z15Z1839Z12Z15Z_h | pail1839Z12Z15Z1839Z12Z15Z[h] | -0.000108 | -0.000015 | 0.092 | 0.013 |
| Lipid | tag1501016Z1123Z6Z_h | Triacylglycerol (15:0/10:1(6Z)/11:2(3Z,6Z)) | -0.000165 | -0.000023 | 0.086 | 0.015 |
| Lipid | pchol12060_c | pchol12060[c] | -0.000159 | -0.000022 | 0.085 | 0.012 |
| Lipid | pchol1719Z1819Z_c | pchol1719Z1819Z[c] | -0.000084 | -0.000012 | 0.064 | 0.009 |
| Lipid | tag901123Z6Z1123Z6Z_h | Triacylglycerol (9:0/11:23Z6Z/11:23Z6Z) | -0.000094 | -0.000013 | 0.053 | 0.007 |
| Lipid | tag1829Z12Z1123Z6Z1123Z6Z_h | Triacylglycerol (18:2(9Z,12Z)/11:2(3Z6Z)/11:2(3Z6Z)) | -0.000076 | -0.000011 | 0.052 | 0.007 |
| Lipid | pchol1102229Z12Z_c | pchol1102229Z12Z[c] | -0.000062 | -0.000009 | 0.046 | 0.006 |
| Lipid | pchol17080_c | pchol17080[c] | -0.000071 | -0.000010 | 0.045 | 0.006 |
| Lipid | tag401123Z6Z120_h | Triacylglycerol (4:0/11:2(3Z6Z)/12:0) | -0.000082 | -0.000012 | 0.042 | 0.006 |
| Lipid | pchol1801443Z6Z9Z12Z_c | pchol1801443Z6Z9Z12Z[c] | -0.000055 | -0.000008 | 0.040 | 0.006 |
| Lipid | pchol1601829Z12Z_c | pchol1601829Z12Z[c] | -0.000045 | -0.000006 | 0.034 | 0.005 |
| Lipid | pchol2119Z1023Z6Z_c | pchol2119Z1023Z6Z[c] | -0.000046 | -0.000006 | 0.032 | 0.005 |
| Lipid | pchol1201839Z12Z15Z_c | pchol1201839Z12Z15Z[c] | -0.000038 | -0.000005 | 0.027 | 0.004 |
| Lipid | tag401016Z1116Z_h | Triacylglycerol (4:0/10:1(6Z)/11:1(6Z)) | -0.000046 | -0.000006 | 0.022 | 0.003 |
| Lipid | pe1201119Z_h | pe1201119Z[h] | -0.000035 | -0.000005 | 0.020 | 0.003 |
| Lipid | tag2501016Z150_h | Triacylglycerol (25:0/10:1(6Z)/15:0) | -0.000022 | -0.000003 | 0.018 | 0.003 |
| Lipid | tag901433Z6Z9Z1433Z6Z9Z_h | Triacylglycerol (9:0/14:33Z6Z9Z/14:33Z6Z9Z) | -0.000020 | -0.000003 | 0.013 | 0.002 |
| Lipid | pchol2001426Z9Z_c | pchol2001426Z9Z[c] | -0.000012 | -0.000002 | 0.009 | 0.001 |
| Lipid | pchol1846Z9Z12Z15Z1819Z_c | pchol1846Z9Z12Z15Z1819Z[c] | -0.000008 | -0.000001 | 0.006 | 0.001 |
| Lipid | tag3001123Z6Z1126Z9Z_h | Triacylglycerol (30:0/11:2(3Z6Z)/11:2(6Z9Z)) | -0.000004 | -0.000001 | 0.003 | 0.000 |
| **Lipid** | **Class total** | **312 species** |  |  | **957.9** | **134.7** |
| Growth-associated maintenance (unchanged) | atp_c | ATP | -39.240000 | -39.272708 | 19901.783 | 19918.372 |
| Growth-associated maintenance (unchanged) | h2o_c | H2O | -29.890000 | -29.890000 | 538.477 | 538.477 |
| Growth-associated maintenance (unchanged) | adp_c | ADP | 39.240000 | 39.240000 | - | - |
| Growth-associated maintenance (unchanged) | h_c | H+ | 39.240000 | 39.240000 | - | - |
| Growth-associated maintenance (unchanged) | pi_c | Phosphate | 39.240000 | 39.240000 | - | - |

**Table S2** Structural properties of iPiccel952 and four published algal models.

| **Property** | **iPiccel952** | **iPiccel952 without fatty acid reactions** | **iCO1515** | **iCre1355** | **iCZ843** | **iAJ526** |
| --- | --- | --- | --- | --- | --- | --- |
| Organism | P. celeri | P. celeri | C. ohadii | C. reinhardtii | C. vulgaris | C. variabilis |
| Genes | 954 | 931 | 1,515 | 1,460 | 843 | 531 |
| Reactions | 3,525 | 1,825 | 2,516 | 2,394 | 2,294 | 1,455 |
| Metabolite species | 2,973 | – | 3,517 | 1,845 | 1,770 | 1,236 |
| Unique metabolites | 2,074 (69.8%) | – | 2,539 (72.2%) | 1,132 (61.4%) | 457 (25.8%) | 770 (62.3%) |
| Compartments | 9 | – | 8 | 11 | 6 | 6 |
| Reactions with gene association | 1,238 (35.1%) | – | 1,975 (78.5%) | 1,924 (80.4%) | 1,838 (80.1%) | 856 (58.8%) |
| Transport reactions | 556 (15.8%) | – | 306 (12.2%) | 449 (18.8%) | 378 (16.5%) | 317 (21.8%) |
| Reactions allowing reverse flux | 640 (18.2%) | – | 196 (7.8%) | 723 (30.2%) | 626 (27.3%) | 561 (38.6%) |
| Reactions per gene | 3.7 | 2.0 | 1.7 | 1.6 | 2.7 | 2.7 |
| Photoautotrophic growth (1/h) | 0.34 | – | 1.71 | ~0.21 | ~0.03 | ~0.43 |
| Active reactions | 1,860 (52.8%) | 531 (29.1%) | 715 (28.4%) | 1,350 (56.4%) | 1,280 (55.8%) | 735 (50.5%) |
| Essential genes | 116 (12.2%) | 116 (12.5%) | 237 (15.6%) | 242 (16.6%) | 188 (22.3%) | 134 (25.2%) |
| Essential reactions | 1,258 (35.7%) | 274 (15.0%) | 670 (26.6%) | 437 (18.3%) | 388 (16.9%) | 268 (18.4%) |

**Table S3A** Growth, protein pool and size of the kcat calibration in the two analysed conditions.

| **Condition** | **Measured growth (1/h)** | **ecGEM growth (1/h)** | **Measured pool (g/gDCW)** | **ecGEM pool after calibration (g/gDCW)** | **Minimal pool demand before calibration (g/gDCW)** | **Stage 1 bound relaxations** | **Stage 2 boosts** | **Rounds** | **Distinct reactions boosted** |
| --- | --- | --- | --- | --- | --- | --- | --- | --- | --- |
| WT-Air | 0.028 | 0.028 | 0.0778 | 0.0771 | 0.666 | 0 | 16 | 17 | 16 |
| WT-CO2 | 0.325 | 0.325 | 0.0913 | 0.0887 | 7.492 | 9 | 48 | 49 | 46 |

**Table S3B** Reactions whose kcat-derived bound was relaxed to 1,000 mmol/gDW/h in Stage 1 of the calibration. WT-Air needed none.

| **Condition** | **Reaction** | **Name** | **Pathway** | **Original bound (mmol/gDW/h)** | **Relaxed bound (mmol/gDW/h)** |
| --- | --- | --- | --- | --- | --- |
| WT-CO2 | FA180ACPHi | fatty-acyl-ACP_hydrolase_(n-C18:0) | Fatty acid metabolism | 0.212 | 1,000 |
| WT-CO2 | AACPS6 | long-chain-fatty-acid-[acyl-carrier-protein]_ligase_(18:0) | Fatty acid metabolism | 0.119 | 1,000 |
| WT-CO2 | ACPCS | 1-aminocyclopropane-1-carboxylate_synthase | Cysteine and methionine metabolism | 0.0168 | 1,000 |
| WT-CO2 | M5TRPI | S-methyl-5-thioribose-1-phosphate_isomerase | Cysteine and methionine metabolism | 0.091 | 1,000 |
| WT-CO2 | PRAIS | phosphoribosylaminoimidazolesuccinocarboxamide_synthase,_cytosol | Nucleotide metabolism | 0.0226 | 1,000 |
| WT-CO2 | PDHe3mr | dihydrolipoamide_dehydrogenase_(E3),_mitochondrial | Glycolysis / Gluconeogenesis | 3.85 | 1,000 |
| WT-CO2 | PCr | Pyruvate carboxylase | Carbon fixation | 0.501 | 1,000 |
| WT-CO2 | NanoG0459 | glycolate dehydrogenase | Glyoxylate metabolism | 1.17 | 1,000 |
| WT-CO2 | NanoG0094 | acireductone synthase | Unassigned | 0.037 | 1,000 |

**Table S3C** Reactions whose kcat was raised in Stage 2 of the calibration. The kcat is given in 1/s and the protein cost in g per mmol/gDW/h.

| **Condition** | **Reaction** | **Name** | **Pathway** | **Gene** | **Rounds boosted** | **Total fold** | **Predicted kcat (1/s)** | **Calibrated kcat (1/s)** | **Predicted cost** | **Calibrated cost** |
| --- | --- | --- | --- | --- | --- | --- | --- | --- | --- | --- |
| WT-Air | GBDP_FWD | guanosine-3',5'-bis(diphosphate)_3'-diphosphatase (conv) | Nucleotide metabolism | GENE4065144 | 1 | 100 | 9.71 | 971 | 2.58 | 0.0258 |
| WT-Air | GTPDPK_FWD | ATP:GTP_3'-diphosphotransferase (conv) | Nucleotide metabolism | GENE4065144 | 2 | 100 | 9.71 | 971 | 2.58 | 0.0258 |
| WT-Air | UG1PUT | UTP---glucose-1-phosphate_uridylyltransferase | Starch metabolism | GENE4063900 | 3 | 100 | 16.6 | 1,661 | 0.881 | 0.00881 |
| WT-Air | GLYCK | Glycerate_kinase | Glyoxylate metabolism | GENE4065643 | 4 | 100 | 3.88 | 388 | 4.48 | 0.0448 |
| WT-Air | I5NT | IMP-5'-nucleotidase | Nucleotide metabolism | GENE4063844 | 5 | 100 | 24.9 | 2,493 | 0.74 | 0.0074 |
| WT-Air | GDTP_FWD | Guanosine_3'-diphosphate_5'-triphosphate_5'-phosphohydrolase (conv) | Nucleotide metabolism | GENE4066888 | 6 | 100 | 46.6 | 4,660 | 0.203 | 0.00203 |
| WT-Air | MALGLUth | malate/glutamate_antiport | Transport, chloroplast | GENE4068355 | 7 | 100 | 16.2 | 1,621 | 0.989 | 0.00989 |
| WT-Air | ADPA | ADP-apyrase | Nucleotide metabolism | GENE4062890 | 8 | 100 | 46.7 | 4,672 | 0.3 | 0.003 |
| WT-Air | ARGSS | argininosuccinate_synthase | Arginine and proline metabolism | GENE4064440 | 9 | 100 | 8.22 | 822 | 1.82 | 0.0182 |
| WT-Air | R5PDP | ribose-phosphate_diphosphokinase | Nucleotide metabolism | GENE4062717 | 10 | 100 | 20.7 | 2,069 | 0.601 | 0.00601 |
| WT-Air | TPIh_FWD | triosephosphate_isomerase (conv) | Glycolysis / Gluconeogenesis | GENE4065450 | 11 | 100 | 6.86 | 686 | 1.24 | 0.0124 |
| WT-Air | GAL | glutamate---ammonia_ligase,_cytosol | Glutamate metabolism | GENE4065958 | 12 | 100 | 16.4 | 1,642 | 0.715 | 0.00715 |
| WT-Air | NanoG0544 | pyruvate, phosphate dikinase | Carbon fixation | GENE4062286 | 13 | 100 | 38.8 | 3,879 | 0.754 | 0.00754 |
| WT-Air | NanoG0437 | glyceraldehyde-3-phosphate dehydrogenase (phosphorylating) | Glycolysis / Gluconeogenesis | GENE4066860 | 14 | 100 | 71.4 | 7,138 | 0.225 | 0.00225 |
| WT-Air | GLPThi | glucose-1-phosphate_adenylyltransferase_(chloroplast) | Starch metabolism | GENE4063059 | 15 | 100 | 45.6 | 4,559 | 0.339 | 0.00339 |
| WT-Air | NanoG1707 | phosphoenolpyruvate carboxylase | Carbon fixation | GENE4062117 | 16 | 100 | 45.9 | 4,588 | 0.663 | 0.00663 |
| WT-CO2 | GBDP_FWD | guanosine-3',5'-bis(diphosphate)_3'-diphosphatase (conv) | Nucleotide metabolism | GENE4065144 | 1, 32 | 10,000 | 9.71 | 97,112 | 2.58 | 0.000258 |
| WT-CO2 | GTPDPK_FWD | ATP:GTP_3'-diphosphotransferase (conv) | Nucleotide metabolism | GENE4065144 | 2, 33 | 10,000 | 9.71 | 97,112 | 2.58 | 0.000258 |
| WT-CO2 | UG1PUT | UTP---glucose-1-phosphate_uridylyltransferase | Starch metabolism | GENE4063900 | 3 | 100 | 16.6 | 1,661 | 0.881 | 0.00881 |
| WT-CO2 | GLYCK | Glycerate_kinase | Glyoxylate metabolism | GENE4065643 | 4 | 100 | 3.88 | 388 | 4.48 | 0.0448 |
| WT-CO2 | I5NT | IMP-5'-nucleotidase | Nucleotide metabolism | GENE4063844 | 5 | 100 | 24.9 | 2,493 | 0.74 | 0.0074 |
| WT-CO2 | GDTP_FWD | Guanosine_3'-diphosphate_5'-triphosphate_5'-phosphohydrolase (conv) | Nucleotide metabolism | GENE4066888 | 6 | 100 | 46.6 | 4,660 | 0.203 | 0.00203 |
| WT-CO2 | MALGLUth | malate/glutamate_antiport | Transport, chloroplast | GENE4068355 | 7 | 100 | 16.2 | 1,621 | 0.989 | 0.00989 |
| WT-CO2 | ADPA | ADP-apyrase | Nucleotide metabolism | GENE4062890 | 8 | 100 | 46.7 | 4,672 | 0.3 | 0.003 |
| WT-CO2 | ARGSS | argininosuccinate_synthase | Arginine and proline metabolism | GENE4064440 | 9 | 100 | 8.22 | 822 | 1.82 | 0.0182 |
| WT-CO2 | R5PDP | ribose-phosphate_diphosphokinase | Nucleotide metabolism | GENE4062717 | 10 | 100 | 20.7 | 2,069 | 0.601 | 0.00601 |
| WT-CO2 | TPIh_FWD | triosephosphate_isomerase (conv) | Glycolysis / Gluconeogenesis | GENE4065450 | 11 | 100 | 6.86 | 686 | 1.24 | 0.0124 |
| WT-CO2 | GAL | glutamate---ammonia_ligase,_cytosol | Glutamate metabolism | GENE4065958 | 12 | 100 | 16.4 | 1,642 | 0.715 | 0.00715 |
| WT-CO2 | NanoG0544 | pyruvate, phosphate dikinase | Carbon fixation | GENE4062286 | 13 | 100 | 38.8 | 3,879 | 0.754 | 0.00754 |
| WT-CO2 | NanoG0437 | glyceraldehyde-3-phosphate dehydrogenase (phosphorylating) | Glycolysis / Gluconeogenesis | GENE4066860 | 14 | 100 | 71.4 | 7,138 | 0.225 | 0.00225 |
| WT-CO2 | GLPThi | glucose-1-phosphate_adenylyltransferase_(chloroplast) | Starch metabolism | GENE4063059 | 15 | 100 | 45.6 | 4,559 | 0.339 | 0.00339 |
| WT-CO2 | NanoG1707 | phosphoenolpyruvate carboxylase | Carbon fixation | GENE4062117 | 16 | 100 | 45.9 | 4,588 | 0.663 | 0.00663 |
| WT-CO2 | PPC | phosphoenolpyruvate_carboxylase | Carbon fixation | GENE4062117 | 17 | 100 | 45.9 | 4,588 | 0.663 | 0.00663 |
| WT-CO2 | NanoG1457 | purine-nucleoside phosphorylase | Nucleotide metabolism | GENE4066817 | 18 | 100 | 42.1 | 4,215 | 0.225 | 0.00225 |
| WT-CO2 | THRAL_FWD | threonine_ammonia-lyase_(serine) (conv) | Glycine, serine and threonine metabolism | GENE4067892 | 19 | 100 | 25.3 | 2,533 | 0.643 | 0.00643 |
| WT-CO2 | NanoG0606 | F0F1-ATP synthase Complex V | Oxidative phosphorylation | GENE4064654 | 20 | 100 | 25.4 | 2,536 | 0.284 | 0.00284 |
| WT-CO2 | ATP_3h_tm | ADP/ATP transporter, mitochondrial | Transport, mitochondrial | GENE4068518 | 21 | 100 | 42.3 | 4,230 | 0.251 | 0.00251 |
| WT-CO2 | AM6PT | ATP:D-mannose_6-phosphotransferase | Carbohydrate metabolism | GENE4067601 | 22 | 100 | 14.9 | 1,488 | 1.05 | 0.0105 |
| WT-CO2 | 3PG_pi_thr | 3-Phospho-D-glycerate transport via triose phosphate translocator (from chloroplast) | Transport, chloroplast | GENE4066437 | 23 | 100 | 20.2 | 2,023 | 0.557 | 0.00557 |
| WT-CO2 | SERth_FWD | Amino_acid_transporter_(ser-L),_chloroplast (conv) | Transport, chloroplast | GENE4067440 | 24 | 100 | 45.2 | 4,517 | 0.499 | 0.00499 |
| WT-CO2 | PGKh_FWD | phosphoglycerate_kinase,_chloroplast (conv) | Glycolysis / Gluconeogenesis | GENE4066618 | 25 | 100 | 116 | 11,607 | 0.118 | 0.00118 |
| WT-CO2 | RPEh_REV | D-Ribulose-5-Phosphate_3-Epimerase (rev) | Pentose phosphate pathway | GENE4063580 | 26 | 100 | 48.9 | 4,886 | 0.458 | 0.00458 |
| WT-CO2 | TKT2h_REV | transketolase_2 (rev) | Pentose phosphate pathway | GENE4068845 | 27 | 100 | 24.4 | 2,439 | 0.88 | 0.0088 |
| WT-CO2 | TKT1h_REV | transketolase_1 (rev) | Pentose phosphate pathway | GENE4068845 | 28 | 100 | 24.4 | 2,439 | 0.88 | 0.0088 |
| WT-CO2 | ATUDm | ATP:UDP_phosphotransferase,_mitochondria | Nucleotide metabolism | GENE4067171 | 29 | 100 | 44 | 4,403 | 0.153 | 0.00153 |
| WT-CO2 | ARGSL | argininosuccinate_lyase | Arginine and proline metabolism | GENE4062108 | 30 | 100 | 45.7 | 4,575 | 0.319 | 0.00319 |
| WT-CO2 | FBA3hi | Sedoheptulose_1,7-bisphosphate_D-glyceraldehyde-3-phosphate-lyase | Carbon fixation | GENE4066500 | 31 | 100 | 60.2 | 6,024 | 0.2 | 0.002 |
| WT-CO2 | NanoG0405 | phosphopyruvate hydratase | Glycolysis / Gluconeogenesis | GENE4066860 | 34 | 100 | 71.4 | 7,138 | 0.225 | 0.00225 |
| WT-CO2 | CY_focytc_h | ferrocytochrome c recycle, chloroplast | Photosynthesis | GENE4064616 | 35 | 100 | 73.9 | 7,387 | 0.057 | 0.00057 |
| WT-CO2 | GLYDH | Glycerate_dehydrogenase | Glyoxylate metabolism | GENE4063220 | 36 | 100 | 97.4 | 9,737 | 0.184 | 0.00184 |
| WT-CO2 | ATAMh | ATP:AMP_phosphotransferase,_chloroplast | Nucleotide metabolism | GENE4064600 | 37 | 100 | 46.1 | 4,610 | 0.156 | 0.00156 |
| WT-CO2 | NanoG1229 | nad(p)+ transhydrogenase (si-specific) | Oxidative phosphorylation | GENE4066856 | 38 | 100 | 142 | 14,201 | 0.106 | 0.00106 |
| WT-CO2 | MPAKI_FWD | D-mannose-6-phosphate_aldose-ketose-isomerase (fwd) | Carbohydrate metabolism | GENE4064372 | 39 | 100 | 38.5 | 3,855 | 0.322 | 0.00322 |
| WT-CO2 | MALOAAtm | Dicarboxylate/tricarboxylate_carrier_(mal:oaa),_mitochondrial | Transport, mitochondrial | GENE4062844 | 40 | 100 | 78.8 | 7,878 | 0.108 | 0.00108 |
| WT-CO2 | SUCFUMtmr_FWD | Succinate/fumarate_antiporter,_mitochondrial (conv) | Transport, mitochondrial | GENE4062844 | 41 | 100 | 78.8 | 7,878 | 0.108 | 0.00108 |
| WT-CO2 | OCTm_FWD | ornithine_carbamoyltransferase,_mitochondrial (fwd) | Arginine and proline metabolism | GENE4067903 | 42 | 100 | 108 | 10,813 | 0.101 | 0.00101 |
| WT-CO2 | G6PIh_FWD | glucose-6-phosphate_isomerase,_chloroplast (conv) | Glycolysis / Gluconeogenesis | GENE4067753 | 43 | 100 | 113 | 11,336 | 0.153 | 0.00153 |
| WT-CO2 | PGIBh_FWD | glucose-6-phosphate_isomerase,_chloroplast_(g6p-B) (conv) | Glycolysis / Gluconeogenesis | GENE4067753 | 44 | 100 | 113 | 11,336 | 0.153 | 0.00153 |
| WT-CO2 | ASPATh_FWD | aspartate_aminotransferase,_chloroplast (conv) | Alanine and aspartate metabolism | GENE4066933 | 45 | 100 | 155 | 15,533 | 0.0869 | 0.000869 |
| WT-CO2 | HCO3Ehi | carbonate_dehydratase_(HCO3_equilibration_reaction),_irreversible,_chloroplast | Carbon fixation | GENE4067678 | 46 | 100 | 17.3 | 1,731 | 0.506 | 0.00506 |
| WT-CO2 | CBFC | cytochrome_b6/f_complex | Photosynthesis | GENE4063369 | 47 | 100 | 24.7 | 2,466 | 0.178 | 0.00178 |
| WT-CO2 | PSIred | photosystem_I_(red_light-activated) | Photosynthesis | GENE4064276 | 48 | 100 | 46.1 | 4,609 | 0.137 | 0.00137 |

**Table S3D** Leave-out test of the calibration. Growth under the measured protein pool of the target condition with no kcat correction, with the corrections found in the other condition, and with the corrections found in the condition itself.

| **Target condition** | **Measured growth (1/h)** | **No correction (1/h)** | **Corrections from the other condition (1/h)** | **Corrections from the condition itself (1/h)** | **Other-condition result, % of measured** |
| --- | --- | --- | --- | --- | --- |
| WT-Air | 0.0280 | 0.0033 | 0.0287 | 0.0282 | 103% |
| WT-CO2 | 0.3249 | 0.0039 | 0.0331 | 0.3249 | 10% |

**Table S4A** Bottleneck enzymes with a kcat control coefficient of 0.01 or higher, with their measured abundance (mg/gCDW).

| **Condition** | **Reaction** | **Name** | **Pathway** | **Control coefficient** | **Flux (mmol/gDW/h)** | **Protein cost** | **Abundance WT-Air** | **Abundance WT-CO2** | **Ratio CO2/Air** |
| --- | --- | --- | --- | --- | --- | --- | --- | --- | --- |
| WT-Air | PPC | phosphoenolpyruvate_carboxylase | Carbon fixation | 0.061 | 7.79 | 0.663 | 2.39 | 2.99 | 1.25 |
| WT-Air | NanoG1457 | purine-nucleoside phosphorylase | Nucleotide metabolism | 0.059 | 22.5 | 0.225 | 0.41 | 0.55 | 1.35 |
| WT-Air | THRAL_FWD | threonine_ammonia-lyase_(serine) (conv) | Glycine, serine and threonine metabolism | 0.055 | 7.26 | 0.643 | 0.06 | 0.09 | 1.57 |
| WT-Air | NanoG0606 | F0F1-ATP synthase Complex V | Oxidative phosphorylation | 0.053 | 15.9 | 0.284 | 25.33 | 40.15 | 1.58 |
| WT-Air | ATP_3h_tm | ADP/ATP transporter, mitochondrial | Transport, mitochondrial | 0.046 | 15.7 | 0.251 | 7.08 | 8.74 | 1.23 |
| WT-Air | AM6PT | ATP:D-mannose_6-phosphotransferase | Carbohydrate metabolism | 0.044 | 3.6 | 1.05 | 0.08 | 0.07 | 0.84 |
| WT-Air | 3PG_pi_thr | 3-Phospho-D-glycerate transport via triose phosphate translocator (from chloroplast) | Transport, chloroplast | 0.043 | 6.58 | 0.557 | 0.03 | 0.05 | 1.92 |
| WT-Air | SERth_FWD | Amino_acid_transporter_(ser-L),_chloroplast (conv) | Transport, chloroplast | 0.042 | 7.25 | 0.499 | 0.01 | 0.01 | 1.16 |
| WT-Air | PGKh_FWD | phosphoglycerate_kinase,_chloroplast (conv) | Glycolysis / Gluconeogenesis | 0.033 | 24.1 | 0.118 | 4.54 | 5.25 | 1.16 |
| WT-Air | RPEh_REV | D-Ribulose-5-Phosphate_3-Epimerase (rev) | Pentose phosphate pathway | 0.031 | 5.85 | 0.458 | 0.70 | 0.91 | 1.30 |
| WT-Air | TKT2h_REV | transketolase_2 (rev) | Pentose phosphate pathway | 0.030 | 2.93 | 0.88 | 2.27 | 2.60 | 1.14 |
| WT-Air | TKT1h_REV | transketolase_1 (rev) | Pentose phosphate pathway | 0.030 | 2.92 | 0.88 | 2.27 | 2.60 | 1.14 |
| WT-Air | ATUDm | ATP:UDP_phosphotransferase,_mitochondria | Nucleotide metabolism | 0.030 | 16.6 | 0.153 | 0.75 | 1.37 | 1.82 |
| WT-Air | ARGSL | argininosuccinate_lyase | Arginine and proline metabolism | 0.028 | 7.51 | 0.319 | 0.23 | 0.34 | 1.46 |
| WT-Air | FBA3hi | Sedoheptulose_1,7-bisphosphate_D-glyceraldehyde-3-phosphate-lyase | Carbon fixation | 0.025 | 10.6 | 0.2 | 5.56 | 9.11 | 1.64 |
| WT-Air | GBDP_FWD | guanosine-3',5'-bis(diphosphate)_3'-diphosphatase (conv) | Nucleotide metabolism | 0.023 | 76.7 | 0.0258 | 0.12 | 0.13 | 1.15 |
| WT-Air | GTPDPK_FWD | ATP:GTP_3'-diphosphotransferase (conv) | Nucleotide metabolism | 0.023 | 76.7 | 0.0258 | 0.12 | 0.13 | 1.15 |
| WT-Air | NanoG0405 | phosphopyruvate hydratase | Glycolysis / Gluconeogenesis | 0.021 | 7.82 | 0.225 | 4.41 | 6.68 | 1.52 |
| WT-Air | CY_focytc_h | ferrocytochrome c recycle, chloroplast | Photosynthesis | 0.017 | 25.4 | 0.057 | 0.04 | 0.04 | 0.96 |
| WT-Air | GLYDH | Glycerate_dehydrogenase | Glyoxylate metabolism | 0.015 | 7.19 | 0.184 | 3.17 | 3.54 | 1.11 |
| WT-Air | ATAMh | ATP:AMP_phosphotransferase,_chloroplast | Nucleotide metabolism | 0.015 | 8.01 | 0.156 | 1.58 | 2.54 | 1.61 |
| WT-Air | NanoG1229 | nad(p)+ transhydrogenase (si-specific) | Oxidative phosphorylation | 0.014 | 11.4 | 0.106 | 1.59 | 1.92 | 1.21 |
| WT-Air | MPAKI_FWD | D-mannose-6-phosphate_aldose-ketose-isomerase (fwd) | Carbohydrate metabolism | 0.014 | 3.6 | 0.322 | 0.09 | 0.08 | 0.87 |
| WT-CO2 | PRAI_FWD | phosphoribosylanthranilate_isomerase (fwd) | Phenylalanine, tyrosine and tryptophan biosynthesis | 0.253 | 0.0194 | 0.287 | 0.00 | 0.01 | 1.65 |

**Table S4B** Share of kcat control and of measured enzyme abundance by pathway (pathways with at least five reactions). Shares are percent of the total over all pathways in each condition.

| **Pathway** | **Reactions** | **Control share WT-Air (%)** | **Control share WT-CO2 (%)** | **Abundance share WT-Air (%)** | **Abundance share WT-CO2 (%)** | **Ratio of shares CO2/Air** |
| --- | --- | --- | --- | --- | --- | --- |
| Nucleotide metabolism | 109 | 17.5 | 0.0 | 5.4 | 5.9 | 1.09 |
| Carbon fixation | 12 | 10.9 | 0.0 | 6.2 | 3.9 | 0.64 |
| Pentose phosphate pathway | 22 | 10.4 | 0.0 | 2.8 | 2.4 | 0.84 |
| Transport, chloroplast | 27 | 9.8 | 0.0 | 0.6 | 0.9 | 1.49 |
| Glycolysis / Gluconeogenesis | 25 | 8.7 | 0.0 | 11.3 | 12.1 | 1.07 |
| Oxidative phosphorylation | 8 | 7.9 | 0.0 | 7.2 | 7.5 | 1.04 |
| Transport, mitochondrial | 19 | 7.4 | 0.0 | 1.7 | 1.6 | 0.98 |
| Carbohydrate metabolism | 9 | 6.5 | 0.0 | 0.2 | 0.2 | 0.94 |
| Glycine, serine and threonine metabolism | 23 | 6.3 | 0.0 | 6.0 | 3.8 | 0.63 |
| Photosynthesis | 13 | 5.6 | 0.0 | 0.2 | 0.2 | 0.94 |
| Arginine and proline metabolism | 26 | 4.3 | 0.0 | 1.5 | 1.5 | 1.04 |
| Glyoxylate metabolism | 11 | 2.2 | 0.0 | 1.4 | 0.8 | 0.58 |
| Alanine and aspartate metabolism | 10 | 0.9 | 0.0 | 2.1 | 1.3 | 0.62 |
| Glutamate metabolism | 10 | 0.8 | 0.0 | 2.9 | 2.8 | 0.96 |
| Transport, extracellular | 50 | 0.7 | 0.0 | 1.3 | 3.0 | 2.24 |
| Glycerophospholipid metabolism | 25 | 0.6 | 0.0 | 0.3 | 0.3 | 1.04 |
| Starch metabolism | 19 | 0.6 | 0.0 | 1.2 | 1.1 | 0.93 |
| Pantothenate and CoA biosynthesis | 12 | 0.3 | 0.0 | 0.5 | 0.6 | 1.21 |
| Pyruvate metabolism | 29 | 0.1 | 0.0 | 2.8 | 2.7 | 0.98 |
| Phenylalanine, tyrosine and tryptophan biosynthesis | 38 | 0.1 | 100.0 | 2.2 | 2.3 | 1.05 |
| Lysine metabolism | 8 | 0.0 | 0.0 | 0.2 | 0.2 | 0.94 |
| Cysteine and methionine metabolism | 42 | 0.0 | 0.0 | 4.5 | 4.9 | 1.09 |
| Amino sugar and nucleotide sugar metabolism | 5 | -0.0 | 0.0 | 0.0 | 0.0 | 0.98 |
| Porphyrin and chlorophyll metabolism | 51 | -0.0 | 0.0 | 3.2 | 4.2 | 1.34 |
| Transport, glyoxysome | 12 | -0.0 | 0.0 | 0.2 | 0.1 | 0.59 |
| beta-Alanine metabolism | 11 | -0.0 | 0.0 | 0.3 | 0.3 | 0.77 |
| Glutathione metabolism | 10 | -0.0 | 0.0 | 2.0 | 1.7 | 0.88 |
| Sphingolipid metabolism | 6 | -0.0 | 0.0 | 0.0 | 0.0 | 0.69 |
| TCA cycle | 10 | -0.0 | 0.0 | 1.2 | 1.0 | 0.86 |
| Valine, leucine and isoleucine metabolism | 41 | -0.0 | 0.0 | 4.0 | 4.4 | 1.09 |
| Nitrogen metabolism | 12 | -0.0 | 0.0 | 2.5 | 2.3 | 0.92 |
| Miscellaneous | 27 | -0.1 | 0.0 | 2.2 | 2.4 | 1.08 |
| Retinol metabolism | 12 | -0.1 | 0.0 | 0.3 | 0.3 | 1.00 |
| Histidine metabolism | 19 | -0.1 | 0.0 | 0.3 | 0.3 | 1.00 |
| Selenoamino acid metabolism | 18 | -0.1 | 0.0 | 1.6 | 1.6 | 1.05 |
| glycan metabolism | 21 | -0.1 | 0.0 | 0.3 | 0.3 | 0.90 |
| Biosynthesis of steroids | 52 | -0.1 | 0.0 | 1.5 | 1.9 | 1.25 |
| Carotenoid biosynthesis | 25 | -0.1 | 0.0 | 1.1 | 1.6 | 1.47 |
| Alternate Carbon Metabolism | 35 | -0.1 | 0.0 | 1.0 | 0.8 | 0.84 |
| Cofactor and Prosthetic Group Biosynthesis | 63 | -0.2 | 0.0 | 1.0 | 1.0 | 0.94 |
| Fatty acid metabolism | 144 | -0.2 | 0.0 | 9.9 | 8.2 | 0.83 |
| Glycerolipid metabolism | 136 | -0.3 | 0.0 | 4.7 | 7.2 | 1.54 |

**Table S5A** Curation log of iPiccel952 by stage. Counts from the curation report are compared with counts in the shipped model.

| **Stage** | **Action** | **Items in curation report** | **Items in shipped model** | **Detail** |
| --- | --- | --- | --- | --- |
| 1 | Biomass objective function rebuilt from LaPanse et al. 2024 | 360 species | – | Table S1 |
| 2 | Elemental imbalance before curation (non-boundary, non-biomass reactions) | 613 of 4,344 | – |  |
| 2 | Hydrogen-only imbalances corrected with a growth check | 485 of 486 | 352 present and balanced | Table S5B |
| 2 | Proton stoichiometry of ATP synthase, cytochrome b6f, cyclic electron flow, photosystem I and ferredoxin-NADP reductase made consistent across the thylakoid, and the stromal-lumenal proton exchange removed | 7 reactions | 7 present | Table S5G |
| 3 | Metabolites with cross-database annotation | 1,702 of 3,235 (52.6%) | – |  |
| 3 | Reactions with cross-database annotation | 2,030 of 4,344 (46.7%) | – |  |
| 3 | SBO terms assigned to all reactions, metabolites and genes | 4,344, 3,235 and 1,084 | – |  |
| 4 | Energy-generating cycles found (NADPH, NADH, FADH2, glutamate and 2-oxoglutarate) | 4 | – |  |
| 4 | Reactions without gene association restricted to one direction | 44 | 14 present | Table S5C |
| 5 | Reactions left with a single direction | 199 | 183 present | Table S5D |
| 5 | Reactions removed (bounds fixed at zero) | 823 | – |  |
| 5 | Metabolites and genes removed after reaction removal | 250 and 132 | – |  |
| 6 | Metabolite formula corrections | 37 | 37 present | Table S5F |
| 7 | Non-GAM biomass coefficients rescaled to 1 g/mmol (MEMOTE convention), factor | 1.191406 | – |  |
|  | Unbounded flux under default conditions, before and after curation | 36.6% to 0.27% | – |  |
|  | Reactions in stoichiometrically balanced cycles (MEMOTE), before and after curation | 1,437 to 2 | – |  |
|  | MEMOTE total score of the final model | 71.5% | – |  |
|  | Reactions, metabolites and genes in the final model | 3,521, 2,985 and 952 | 3,525, 2,973 and 954 |  |
|  | Non-boundary, non-biomass reactions with elemental imbalance in the final model | – | 42 | Table S5E |

**Table S5B** Hydrogen balance corrections present in the shipped model. The equation before is from the curation report and the equation after is from the shipped model.

| **Reaction** | **Equation before** | **Equation in shipped model** | **Proton species** | **Change (mol H)** |
| --- | --- | --- | --- | --- |
| PAIL18111Z160t | atp_c + h2o_c + pail18111Z160_c --> adp_c + h_c + pail18111Z160_e + pi_c | atp_c + h2o_c + pail18111Z160_c --> adp_c + pail18111Z160_e + pi_c | h_c | -1.000000 |
| PAIL1819Z160t | atp_c + h2o_c + pail1819Z160_c --> adp_c + h_c + pail1819Z160_e + pi_c | atp_c + h2o_c + pail1819Z160_c --> adp_c + pail1819Z160_e + pi_c | h_c | -1.000000 |
| PE1801819Zt | atp_c + h2o_c + pe1801819Z_c --> adp_c + h_c + pe1801819Z_e + pi_c | atp_c + h2o_c + pe1801819Z_c --> adp_c + pe1801819Z_e + pi_c | h_c | -1.000000 |
| PE1801829Z12Zt | atp_c + h2o_c + pe1801829Z12Z_c --> adp_c + h_c + pe1801829Z12Z_e + pi_c | atp_c + h2o_c + pe1801829Z12Z_c --> adp_c + pe1801829Z12Z_e + pi_c | h_c | -1.000000 |
| PE1801835Z9Z12Zt | atp_c + h2o_c + pe1801835Z9Z12Z_c --> adp_c + h_c + pe1801835Z9Z12Z_e + pi_c | atp_c + h2o_c + pe1801835Z9Z12Z_c --> adp_c + pe1801835Z9Z12Z_e + pi_c | h_c | -1.000000 |
| PE1801845Z9Z12Z15Zt | atp_c + h2o_c + pe1801845Z9Z12Z15Z_c --> adp_c + h_c + pe1801845Z9Z12Z15Z_e + pi_c | atp_c + h2o_c + pe1801845Z9Z12Z15Z_c --> adp_c + pe1801845Z9Z12Z15Z_e + pi_c | h_c | -1.000000 |
| PE18111Z1819Zt | atp_c + h2o_c + pe18111Z1819Z_c --> adp_c + h_c + pe18111Z1819Z_e + pi_c | atp_c + h2o_c + pe18111Z1819Z_c --> adp_c + pe18111Z1819Z_e + pi_c | h_c | -1.000000 |
| PE18111Z1829Z12Zt | atp_c + h2o_c + pe18111Z1829Z12Z_c --> adp_c + h_c + pe18111Z1829Z12Z_e + pi_c | atp_c + h2o_c + pe18111Z1829Z12Z_c --> adp_c + pe18111Z1829Z12Z_e + pi_c | h_c | -1.000000 |
| PE18111Z1835Z9Z12Zt | atp_c + h2o_c + pe18111Z1835Z9Z12Z_c --> adp_c + h_c + pe18111Z1835Z9Z12Z_e + pi_c | atp_c + h2o_c + pe18111Z1835Z9Z12Z_c --> adp_c + pe18111Z1835Z9Z12Z_e + pi_c | h_c | -1.000000 |
| PE18111Z1845Z9Z12Z15Zt | atp_c + h2o_c + pe18111Z1845Z9Z12Z15Z_c --> adp_c + h_c + pe18111Z1845Z9Z12Z15Z_e + pi_c | atp_c + h2o_c + pe18111Z1845Z9Z12Z15Z_c --> adp_c + pe18111Z1845Z9Z12Z15Z_e + pi_c | h_c | -1.000000 |
| PE1819Z1819Zt | atp_c + h2o_c + pe1819Z1819Z_c --> adp_c + h_c + pe1819Z1819Z_e + pi_c | atp_c + h2o_c + pe1819Z1819Z_c --> adp_c + pe1819Z1819Z_e + pi_c | h_c | -1.000000 |
| PE1819Z1829Z12Zt | atp_c + h2o_c + pe1819Z1829Z12Z_c --> adp_c + h_c + pe1819Z1829Z12Z_e + pi_c | atp_c + h2o_c + pe1819Z1829Z12Z_c --> adp_c + pe1819Z1829Z12Z_e + pi_c | h_c | -1.000000 |
| PE1819Z1835Z9Z12Zt | atp_c + h2o_c + pe1819Z1835Z9Z12Z_c --> adp_c + h_c + pe1819Z1835Z9Z12Z_e + pi_c | atp_c + h2o_c + pe1819Z1835Z9Z12Z_c --> adp_c + pe1819Z1835Z9Z12Z_e + pi_c | h_c | -1.000000 |
| PE1819Z1845Z9Z12Z15Zt | atp_c + h2o_c + pe1819Z1845Z9Z12Z15Z_c --> adp_c + h_c + pe1819Z1845Z9Z12Z15Z_e + pi_c | atp_c + h2o_c + pe1819Z1845Z9Z12Z15Z_c --> adp_c + pe1819Z1845Z9Z12Z15Z_e + pi_c | h_c | -1.000000 |
| PE1829Z12Z1835Z9Z12Zt | atp_c + h2o_c + pe1829Z12Z1835Z9Z12Z_c --> adp_c + h_c + pe1829Z12Z1835Z9Z12Z_e + pi_c | atp_c + h2o_c + pe1829Z12Z1835Z9Z12Z_c --> adp_c + pe1829Z12Z1835Z9Z12Z_e + pi_c | h_c | -1.000000 |
| PG18111Z160t | atp_c + h2o_c + pg18111Z160_c --> adp_c + h_c + pg18111Z160_e + pi_c | atp_c + h2o_c + pg18111Z160_c --> adp_c + pg18111Z160_e + pi_c | h_c | -1.000000 |
| PG1819Z160t | atp_c + h2o_c + pg1819Z160_c --> adp_c + h_c + pg1819Z160_e + pi_c | atp_c + h2o_c + pg1819Z160_c --> adp_c + pg1819Z160_e + pi_c | h_c | -1.000000 |
| PGP18111Z160t | atp_c + h2o_c + pgp18111Z160_c --> adp_c + h_c + pgp18111Z160_e + pi_c | atp_c + h2o_c + pgp18111Z160_c --> adp_c + pgp18111Z160_e + pi_c | h_c | -1.000000 |
| PGP1819Z160t | atp_c + h2o_c + pgp1819Z160_c --> adp_c + h_c + pgp1819Z160_e + pi_c | atp_c + h2o_c + pgp1819Z160_c --> adp_c + pgp1819Z160_e + pi_c | h_c | -1.000000 |
| NOS | 2.0 arg__L_c + 3.0 nadph_c + 4.0 o2_c --> 2.0 citr__L_c + 4.0 h2o_c + 2.0 h_c + 3.0 nadp_c + 2.0 no_c | 2.0 arg__L_c + 3.0 nadph_c + 4.0 o2_c --> 2.0 citr__L_c + 4.0 h2o_c + 3.0 nadp_c + 2.0 no_c | h_c | -2.000000 |
| NOS1 | arg__L_c + nadph_c + o2_c --> h2o_c + h_c + nadp_c + nwharg_c | arg__L_c + nadph_c + o2_c --> h2o_c + nadp_c + nwharg_c | h_c | -1.000000 |
| ORNTArm | akg_m + orn_m <=> glu5sa_m + glu__L_m + 3.0 h_m | akg_m + orn_m --> glu5sa_m + glu__L_m | h_m | -3.000000 |
| OCT | cbp_c + orn_c --> citr__L_c + 3.0 h_c + pi_c | cbp_c + orn_c --> citr__L_c + pi_c | h_c | -3.000000 |
| OCTh | cbp_h + orn_h <=> citr__L_h + 3.0 h_h + pi_h | cbp_h + orn_h <=> citr__L_h + pi_h | h_h | -3.000000 |
| OCTm | cbp_m + orn_m <=> citr__L_m + 3.0 h_m + pi_m | cbp_m + orn_m <=> citr__L_m + pi_m | h_m | -3.000000 |
| PPC | co2_c + h2o_c + pep_c --> h_c + oaa_c + pi_c | co2_c + h2o_c + pep_c --> oaa_c + pi_c | h_c | -1.000000 |
| PPDKh | atp_h + 3.0 h_h + pi_h + pyr_h --> amp_h + pep_h + ppi_h | atp_h + pi_h + pyr_h --> amp_h + pep_h + ppi_h | h_h | 3.000000 |
| SBP | h2o_h + s17bp_h <=> h_h + pi_h + s7p_h | h2o_h + s17bp_h --> pi_h + s7p_h | h_h | -1.000000 |
| SHSL4h | h2o_h + h_h + suchms_h --> 2obut_h + nh4_h + succ_h | h2o_h + suchms_h --> 2obut_h + nh4_h + succ_h | h_h | 1.000000 |
| PRDPARh | gln__L_h + h2o_h + prpp_h --> glu__L_h + h_h + ppi_h + pram_h | gln__L_h + h2o_h + prpp_h --> glu__L_h + ppi_h + pram_h | h_h | -1.000000 |
| PAPA18111Z1819Z | h2o_c + pa18111Z1819Z_c --> 12dgr18111Z1819Z_c + h_c + pi_c | h2o_c + pa18111Z1819Z_c --> 12dgr18111Z1819Z_c + pi_c | h_c | -1.000000 |
| SQDGW6DS1829Z12Z160 | 2.0 fdxrd_h + 4.0 h_h + o2_h + sqdg1819Z160_h --> 2.0 fdxox_h + 2.0 h2o_h + sqdg1829Z12Z160_h | 2.0 fdxrd_h + o2_h + sqdg1819Z160_h --> 2.0 fdxox_h + 2.0 h2o_h + sqdg1829Z12Z160_h | h_h | 4.000000 |
| PGPP18111Z160 | h2o_c + pgp18111Z160_c --> h_c + pg18111Z160_c + pi_c | h2o_c + pgp18111Z160_c --> pg18111Z160_c + pi_c | h_c | -1.000000 |
| PGPP18111Z160h | h2o_h + pgp18111Z160_h --> h_h + pg18111Z160_h + pi_h | h2o_h + pgp18111Z160_h --> pg18111Z160_h + pi_h | h_h | -1.000000 |
| PGPP1819Z160 | h2o_c + pgp1819Z160_c --> h_c + pg1819Z160_c + pi_c | h2o_c + pgp1819Z160_c --> pg1819Z160_c + pi_c | h_c | -1.000000 |
| PGPP1819Z160h | h2o_h + pgp1819Z160_h --> h_h + pg1819Z160_h + pi_h | h2o_h + pgp1819Z160_h --> pg1819Z160_h + pi_h | h_h | -1.000000 |
| PGLYCPh | 2pglyc_h + h2o_h --> glyclt_h + h_h + pi_h | 2pglyc_h + h2o_h --> glyclt_h + pi_h | h_h | -1.000000 |
| THRAL | h_h + ser__L_h <=> nh4_h + pyr_h | ser__L_h <-- nh4_h + pyr_h | h_h | 1.000000 |
| PGLYCPx | 2pglyc_x + h2o_x --> glyclt_x + h_x + pi_x | 2pglyc_x + h2o_x --> glyclt_x + pi_x | h_x | -1.000000 |
| MOTA | 2kmb_c + glu__L_c + h_c --> akg_c + met__L_c | 2kmb_c + glu__L_c --> akg_c + met__L_c | h_c | 1.000000 |
| MS | h_m + hcys__L_m + mhpglu_m --> hpglu_m + met__L_m | hcys__L_m + mhpglu_m --> hpglu_m + met__L_m | h_m | 1.000000 |
| NADS1 | atp_c + dnad_c + nh4_c --> amp_c + h_c + nad_c + ppi_c | atp_c + dnad_c + nh4_c --> amp_c + nad_c + ppi_c | h_c | -1.000000 |
| NITR | h_c + nadh_c + no3_c --> h2o_c + nad_c + no2_c | nadh_c + no3_c --> h2o_c + nad_c + no2_c | h_c | 1.000000 |
| NITR(NO) | focytc_h + 2.0 h_h + no2_h --> ficytc_h + h2o_h + no_h | focytc_h + no2_h --> ficytc_h + h2o_h + no_h | h_h | 2.000000 |
| NO2R | 6.0 fdxrd_h + 8.0 h_h + no2_h --> 6.0 fdxox_h + 2.0 h2o_h + nh4_h | 6.0 fdxrd_h + 6.0 h_h + no2_h --> 6.0 fdxox_h + 2.0 h2o_h + nh4_h | h_h | 2.000000 |
| NODOx | nadh_c + 2.0 no_c + 2.0 o2_c --> h_c + nad_c + 2.0 no3_c | nadh_c + 2.0 no_c + 2.0 o2_c --> nad_c + 2.0 no3_c | h_c | -1.000000 |
| NODOy | nadph_c + 2.0 no_c + 2.0 o2_c --> h_c + nadp_c + 2.0 no3_c | nadph_c + 2.0 no_c + 2.0 o2_c --> nadp_c + 2.0 no3_c | h_c | -1.000000 |
| OGMG | 4izp_c + 3.0 h2o_c + h_c + 2.0 nad_c --> 4ogm_c + for_c + 2.0 nadh_c + nh4_c | 4izp_c + 3.0 h2o_c + 2.0 h_c + 2.0 nad_c --> 4ogm_c + for_c + 2.0 nadh_c + nh4_c | h_c | -1.000000 |
| MTAM | 5fthf_m + 2.0 h_m --> h2o_m + methf_m | 5fthf_m --> h2o_m + methf_m | h_m | 2.000000 |
| MTHFO | h_m + mlthf_m + nadh_m --> 5mthf_m + nad_m | mlthf_m + nadh_m --> 5mthf_m + nad_m | h_m | 1.000000 |
| MTHFO(nadp) | h_m + mlthf_m + nadph_m --> 5mthf_m + nadp_m | mlthf_m + nadph_m --> 5mthf_m + nadp_m | h_m | 1.000000 |
| PBALm | ala-B_m + atp_m + pant-R_m --> amp_m + 3.0 h_m + pnto-R_m + ppi_m | ala-B_m + atp_m + pant-R_m --> amp_m + pnto-R_m + ppi_m | h_m | -3.000000 |
| PPATDh | h_h + 2.0 pyr_h --> alac-S_h + co2_h | 2.0 pyr_h --> alac-S_h + co2_h | h_h | 1.000000 |
| PPATDm | h_m + 2.0 pyr_m --> alac-S_m + co2_m | 2.0 pyr_m --> alac-S_m + co2_m | h_m | 1.000000 |
| PPCL | 4ppan_c + atp_c + cys__L_c --> 4ppcys_c + amp_c + h_c + ppi_c | 4ppan_c + atp_c + cys__L_c --> 4ppcys_c + amp_c + ppi_c | h_c | -1.000000 |
| PPCL(CTP) | 4ppan_c + ctp_c + cys__L_c --> 4ppcys_c + cmp_c + h_c + ppi_c | 4ppan_c + ctp_c + cys__L_c --> 4ppcys_c + cmp_c + ppi_c | h_c | -1.000000 |
| PGLh | 6pgl_h + h2o_h --> 6pgc_h + h_h | 6pgl_h + h2o_h --> 6pgc_h | h_h | -1.000000 |
| RBK | atp_c + rib-D_c --> adp_c + h_c + r5p_c | atp_c + rib-D_c --> adp_c + r5p_c | h_c | -1.000000 |
| RPDPK | atp_h + r5p_h <=> amp_h + h_h + prpp_h | atp_h + r5p_h <=> amp_h + prpp_h | h_h | -1.000000 |
| PHETL | atp_m + phe__L_m + trnaphe_m --> amp_m + h_m + phetrna_m + ppi_m | atp_m + phe__L_m + trnaphe_m --> amp_m + phetrna_m + ppi_m | h_m | -1.000000 |
| PPDH | h_c + pphn_c --> co2_c + h2o_c + phpyr_c | pphn_c --> co2_c + h2o_c + phpyr_c | h_c | 1.000000 |
| SKDH | 3dhsk_h + h_h + nadph_h <=> nadp_h + skm_h | 3dhsk_h + nadph_h <=> nadp_h + skm_h | h_h | 1.000000 |
| SKK | atp_h + skm_h --> adp_h + h_h + skm5p_h | atp_h + skm_h --> adp_h + skm5p_h | h_h | -1.000000 |
| TYRTAh | 34hpp_h + glu__L_h + h_h <=> akg_h + tyr__L_h | 34hpp_h + glu__L_h --> akg_h + tyr__L_h | h_h | 1.000000 |
| TYRTAm | 34hpp_m + glu__L_m + h_m --> akg_m + tyr__L_m | 34hpp_m + glu__L_m --> akg_m + tyr__L_m | h_m | 1.000000 |
| ORDC | h_c + orn_c --> co2_c + ptrc_c | orn_c --> co2_c + ptrc_c | h_c | 1.000000 |
| ORDCh | h_h + orn_h --> co2_h + ptrc_h | orn_h --> co2_h + ptrc_h | h_h | 1.000000 |
| MPML | atp_h + h2o_h + mg2_h + ppp9_h --> adp_h + h_h + mppp9_h + pi_h | atp_h + h2o_h + mg2_h + ppp9_h --> adp_h + 2.0 h_h + mppp9_h + pi_h | h_h | 1.000000 |
| MPMT | amet_h + mppp9_h <=> ahcys_h + h_h + mppp9me_h | amet_h + mppp9_h <=> ahcys_h + mppp9me_h | h_h | -1.000000 |
| PCHLDR | 2.0 atp_h + 2.0 fdxrd_h + 2.0 h2o_h + pchlda_h --> 2.0 adp_h + chlda_h + 2.0 fdxox_h + 2.0 pi_h | 2.0 atp_h + 2.0 fdxrd_h + 2.0 h2o_h + 2.0 h_h + pchlda_h --> 2.0 adp_h + chlda_h + 2.0 fdxox_h + 2.0 pi_h | h_h | -2.000000 |
| PHPHBDDMC | 3.0 h_h + phphbda_h <=> mfor_h + pyphphbda_h | phphbda_h <=> mfor_h + pyphphbda_h | h_h | 3.000000 |
| PPBNGS | 2.0 5aop_h + h_h --> 2.0 h2o_h + ppbng_h | 2.0 5aop_h --> 2.0 h2o_h + ppbng_h | h_h | 1.000000 |
| OCDOR | h_c + malcoa_c + nadph_c <=> 3opcoa_c + h2o_c + nadp_c | malcoa_c + nadph_c <-- 3opcoa_c + h2o_c + nadp_c | h_c | 1.000000 |
| PCNO | nadp_c + ppcoa_c <=> h_c + nadph_c + prpncoa_c | nadp_c + ppcoa_c --> nadph_c + prpncoa_c | h_c | -1.000000 |
| PPACOALh | atp_h + coa_h + ppa_h --> amp_h + h_h + ppcoa_h + ppi_h | atp_h + coa_h + ppa_h --> amp_h + ppcoa_h + ppi_h | h_h | -1.000000 |
| NTPD | h2o_c + xtp_c --> h_c + ppi_c + xmp_c | h2o_c + xtp_c --> ppi_c + xmp_c | h_c | -1.000000 |
| PPRGL | atp_c + gly_c + pram_c --> adp_c + gar_c + 2.0 h_c + pi_c | atp_c + gly_c + pram_c --> adp_c + gar_c + pi_c | h_c | -2.000000 |
| PPRGLh | atp_h + gly_h + pram_h --> adp_h + gar_h + h_h + pi_h | atp_h + gly_h + pram_h --> adp_h + gar_h + pi_h | h_h | -1.000000 |
| PRAIC | air_c + co2_c <=> 5aizc_c + 2.0 h_c | air_c + co2_c <=> 5aizc_c | h_c | -2.000000 |
| PRAICh | air_h + co2_h <=> 5aizc_h + h_h | air_h + co2_h <=> 5aizc_h | h_h | -1.000000 |
| PRAIS | 5aizc_c + asp__L_c + atp_c --> 25aics_c + adp_c + h_c + pi_c | 5aizc_c + asp__L_c + atp_c --> 25aics_c + adp_c + pi_c | h_c | -1.000000 |
| PRAISh | 5aizc_h + asp__L_h + atp_h --> 25aics_h + adp_h + h_h + pi_h | 5aizc_h + asp__L_h + atp_h --> 25aics_h + adp_h + pi_h | h_h | -1.000000 |
| PRFGCLh | atp_h + fpram_h --> adp_h + air_h + h_h + pi_h | atp_h + fpram_h --> adp_h + air_h + pi_h | h_h | -1.000000 |
| PRFGS | atp_c + fgam_c + gln__L_c + h2o_c --> adp_c + fpram_c + glu__L_c + h_c + pi_c | atp_c + fgam_c + gln__L_c + h2o_c --> adp_c + fpram_c + glu__L_c + pi_c | h_c | -1.000000 |
| PRFGSh | atp_h + fgam_h + gln__L_h + h2o_h --> adp_h + fpram_h + glu__L_h + h_h + pi_h | atp_h + fgam_h + gln__L_h + h2o_h --> adp_h + fpram_h + glu__L_h + pi_h | h_h | -1.000000 |
| R5PDP | atp_c + r5p_c --> amp_c + h_c + prpp_c | atp_c + r5p_c --> amp_c + prpp_c | h_c | -1.000000 |
| SUAT | atp_c + h_c + so4_c --> aps_c + ppi_c | atp_c + 2.0 h_c + so4_c --> aps_c + ppi_c | h_c | -1.000000 |
| UGAH | h2o_m + h_m + udg_m <=> nh4_m + urdglyc_m | h2o_m + udg_m <=> nh4_m + urdglyc_m | h_m | 1.000000 |
| ORPDC | h_c + orot5p_c --> co2_c + ump_c | orot5p_c --> co2_c + ump_c | h_c | 1.000000 |
| UTGAL | atp_c + gln__L_c + h2o_c + utp_c --> adp_c + ctp_c + glu__L_c + h_c + pi_c | atp_c + gln__L_c + h2o_c + utp_c --> adp_c + ctp_c + glu__L_c + pi_c | h_c | -1.000000 |
| LLDH(ferr)x | 2.0 ficytc_x + lac__L_x --> 2.0 focytc_x + 2.0 h_x + pyr_x | 2.0 ficytc_x + lac__L_x --> 2.0 focytc_x + pyr_x | h_x | -2.000000 |
| PYRDC | h_m + pyr_m --> acald_m + co2_m | pyr_m --> acald_m + co2_m | h_m | 1.000000 |
| PYRShi | coa_h + 2.0 fdxox_h + pyr_h --> accoa_h + co2_h + 2.0 fdxrd_h + h_h | coa_h + 2.0 fdxox_h + pyr_h --> accoa_h + co2_h + 2.0 fdxrd_h + 2.0 h_h | h_h | 1.000000 |
| R5PFL | r5p_h --> db4p_h + for_h + h_h | r5p_h --> db4p_h + for_h | h_h | -1.000000 |
| RIBFS | 2.0 dmlz_h + h_h --> 4r5au_h + ribflv_h | 2.0 dmlz_h --> 4r5au_h + ribflv_h | h_h | 1.000000 |
| PHOMSELCYSL | h_c + phom_c + selcys_c --> pi_c + selcyst_c | phom_c + selcys_c --> pi_c + selcyst_c | h_c | 1.000000 |
| PHOMSELCYSLh | h_h + phom_h + selcys_h --> pi_h + selcyst_h | phom_h + selcys_h --> pi_h + selcyst_h | h_h | 1.000000 |
| SEAHCYSHYD | h2o_c + seahcys_c <=> adn_c + h_c + selhcys_c | h2o_c + seahcys_c <=> adn_c + selhcys_c | h_c | -1.000000 |
| SELCYSLY2 | h_c + pdx5p_c + selcys_c --> ala__L_c + pydx5p_c + seln_c | pdx5p_c + selcys_c --> ala__L_c + pydx5p_c + seln_c | h_c | 1.000000 |
| SELCYSTL | h2o_c + selcyst_c --> h_c + nh4_c + pyr_c + selhcys_c | h2o_c + selcyst_c --> nh4_c + pyr_c + selhcys_c | h_c | -1.000000 |
| SELCYSTLh | h2o_h + selcyst_h --> h_h + nh4_h + pyr_h + selhcys_h | h2o_h + selcyst_h --> nh4_h + pyr_h + selhcys_h | h_h | -1.000000 |
| SECYSTL | atp_m + ser__L_m + trnasecys_m --> amp_m + h_m + ppi_m + sertrna-LPAREN-sec-RPAREN_m | atp_m + ser__L_m + trnasecys_m --> amp_m + ppi_m + sertrna-LPAREN-sec-RPAREN_m | h_m | -1.000000 |
| SLDK | atp_m + h2o_m + seln_m --> amp_m + 2.0 h_m + pi_m + selnp_m | atp_m + h2o_m + seln_m --> amp_m + pi_m + selnp_m | h_m | -2.000000 |
| SERPT | h_c + pmtcoa_c + ser__L_c --> 3dsphgn_c + co2_c + coa_c | pmtcoa_c + ser__L_c --> 3dsphgn_c + co2_c + coa_c | h_c | 1.000000 |
| SULO | h2o_c + hso3_c + o2_c --> h2o2_c + h_c + so4_c | h2o_c + hso3_c + o2_c --> h2o2_c + 2.0 h_c + so4_c | h_c | 1.000000 |
| SULOm | h2o_m + hso3_m + o2_m --> h2o2_m + h_m + so4_m | h2o_m + hso3_m + o2_m --> h2o2_m + 2.0 h_m + so4_m | h_m | 1.000000 |
| TAL | thr__L_h --> 2obut_h + h_c + nh4_h | thr__L_h --> 2obut_h + nh4_h | h_c | -1.000000 |
| PYDXDH | h2o_c + o2_c + pydx_c --> 4pyrdx_c + h2o2_c + h_c | h2o_c + o2_c + pydx_c --> 4pyrdx_c + h2o2_c | h_c | -1.000000 |
| PYDXOR | h_c + nadph_c + pydx_c <=> nadp_c + pydxn_c | nadph_c + pydx_c --> nadp_c + pydxn_c | h_c | 1.000000 |
| PYDXS | g3p_c + gln__L_c + ru5p-D_c --> glu__L_c + 3.0 h2o_c + h_c + pi_c + pydx5p_c | g3p_c + gln__L_c + ru5p-D_c --> glu__L_c + 3.0 h2o_c + pi_c + pydx5p_c | h_c | -1.000000 |
| uahCv | h2o_c + 2.0 h_c + urea_c --> co2_c + 2.0 nh4_c | h2o_c + urea_c --> co2_c + 2.0 nh4_c | h_c | 2.000000 |
| tooCv | h2o_h + h_h + o2_h + tyr__L_h --> 34hpp_h + h2o2_h + nh4_h | h2o_h + o2_h + tyr__L_h --> 34hpp_h + h2o2_h + nh4_h | h_h | 1.000000 |
| pfkCv | atp_c + f6p-B_c --> adp_c + fdp-B_c + h_c | atp_c + f6p-B_c --> adp_c + fdp-B_c | h_c | -1.000000 |
| sbtr2hCv | nad_h + sbt-D_h --> fru-B_h + h_h + nadh_h | nad_h + sbt-D_h --> fru-B_h + nadh_h | h_h | -1.000000 |
| RETACIH | h2o_h + retac_h --> ac_h + h_h + retinol-cis-11_h | h2o_h + retac_h --> ac_h + retinol-cis-11_h | h_h | -1.000000 |
| RETINOLSAT | h_h + nadph_h + retinol_h --> dhretinol_h + nadp_h | nadph_h + retinol_h --> dhretinol_h + nadp_h | h_h | 1.000000 |
| RETPALMIH | h2o_h + retpalm_h --> h_h + hdca_h + retinol-cis-11_h | h2o_h + retpalm_h --> hdca_h + retinol-cis-11_h | h_h | -1.000000 |
| PHCDCv | 1p3h5c_c + 2.0 h2o_c + nad_c --> e4hglu_c + h_c + nadh_c | 1p3h5c_c + 2.0 h2o_c + nad_c --> e4hglu_c + nadh_c | h_c | -1.000000 |
| PHCDbCv | 1p3h5c_c + 2.0 h2o_c + nadp_c --> e4hglu_c + h_c + nadph_c | 1p3h5c_c + 2.0 h2o_c + nadp_c --> e4hglu_c + nadph_c | h_c | -1.000000 |
| PHCHGSCv | 1p3h5c_c + h2o_c + h_c --> 4hglusa_c | 1p3h5c_c + h2o_c --> 4hglusa_c | h_c | 1.000000 |
| RBK_L1 | atp_c + rbl__L_c --> adp_c + h_c + ru5p__L_c | atp_c + rbl__L_c --> adp_c + ru5p__L_c | h_c | -1.000000 |
| FACOAL250(2OH)c | atp_c + coa_c + pentacosac(2OH)_c --> amp_c + pentacoscoa(2OH)_c + ppi_c | atp_c + coa_c + 5.0 h_c + pentacosac(2OH)_c --> amp_c + pentacoscoa(2OH)_c + ppi_c | h_c | -5.000000 |
| FACOAL250(2OH)h | atp_h + coa_h + pentacosac(2OH)_h --> amp_h + pentacoscoa(2OH)_h + ppi_h | atp_h + coa_h + 5.0 h_h + pentacosac(2OH)_h --> amp_h + pentacoscoa(2OH)_h + ppi_h | h_h | -5.000000 |
| MGDG160160_SYN | 12dgr160160_h + udpgal_h --> h_h + mgdg160160_h + udp_h | 12dgr160160_h + udpgal_h --> mgdg160160_h + udp_h | h_h | -1.000000 |
| MGDG1601626Z9Z_SYN | 12dgr1601626Z9Z_h + udpgal_h --> 2.0 h_h + mgdg1601626Z9Z_h + udp_h | 12dgr1601626Z9Z_h + udpgal_h --> mgdg1601626Z9Z_h + udp_h | h_h | -2.000000 |
| MGDG1601636Z9Z12Z_SYN | 12dgr1601636Z9Z12Z_h + udpgal_h --> 2.0 h_h + mgdg1601636Z9Z12Z_h + udp_h | 12dgr1601636Z9Z12Z_h + udpgal_h --> mgdg1601636Z9Z12Z_h + udp_h | h_h | -2.000000 |
| MGDG1601819Z_SYN | 12dgr1601819Z_h + udpgal_h --> h_h + mgdg1601819Z_h + udp_h | 12dgr1601819Z_h + udpgal_h --> mgdg1601819Z_h + udp_h | h_h | -1.000000 |
| MGDG1601826Z9Z_SYN | 12dgr1601826Z9Z_h + udpgal_h --> 2.0 h_h + mgdg1601826Z9Z_h + udp_h | 12dgr1601826Z9Z_h + udpgal_h --> mgdg1601826Z9Z_h + udp_h | h_h | -2.000000 |
| MGDG1601829Z12Z_SYN | 12dgr1601829Z12Z_h + udpgal_h --> h_h + mgdg1601829Z12Z_h + udp_h | 12dgr1601829Z12Z_h + udpgal_h --> mgdg1601829Z12Z_h + udp_h | h_h | -1.000000 |
| MGDG1601839Z12Z15Z_SYN | 12dgr1601839Z12Z15Z_h + udpgal_h --> 2.0 h_h + mgdg1601839Z12Z15Z_h + udp_h | 12dgr1601839Z12Z15Z_h + udpgal_h --> mgdg1601839Z12Z15Z_h + udp_h | h_h | -2.000000 |
| MGDG1619Z1829Z12Z_SYN | 12dgr1619Z1829Z12Z_h + udpgal_h --> 2.0 h_h + mgdg1619Z1829Z12Z_h + udp_h | 12dgr1619Z1829Z12Z_h + udpgal_h --> mgdg1619Z1829Z12Z_h + udp_h | h_h | -2.000000 |
| MGDG1619Z1839Z12Z15Z_SYN | 12dgr1619Z1839Z12Z15Z_h + udpgal_h --> h_h + mgdg1619Z1839Z12Z15Z_h + udp_h | 12dgr1619Z1839Z12Z15Z_h + udpgal_h --> mgdg1619Z1839Z12Z15Z_h + udp_h | h_h | -1.000000 |
| MGDG1819Z1829Z12Z_SYN | 12dgr1819Z1829Z12Z_h + udpgal_h --> h_h + mgdg1819Z1829Z12Z_h + udp_h | 12dgr1819Z1829Z12Z_h + udpgal_h --> mgdg1819Z1829Z12Z_h + udp_h | h_h | -1.000000 |
| MGDG1829Z12Z140_SYN | 12dgr1829Z12Z140_h + udpgal_h --> 2.0 h_h + mgdg1829Z12Z140_h + udp_h | 12dgr1829Z12Z140_h + udpgal_h --> mgdg1829Z12Z140_h + udp_h | h_h | -2.000000 |
| MGDG1829Z12Z1626Z9Z_SYN | 12dgr1829Z12Z1626Z9Z_h + udpgal_h --> 2.0 h_h + mgdg1829Z12Z1626Z9Z_h + udp_h | 12dgr1829Z12Z1626Z9Z_h + udpgal_h --> mgdg1829Z12Z1626Z9Z_h + udp_h | h_h | -2.000000 |
| MGDG1829Z12Z1839Z12Z15Z_SYN | 12dgr1829Z12Z1839Z12Z15Z_h + udpgal_h --> 2.0 h_h + mgdg1829Z12Z1839Z12Z15Z_h + udp_h | 12dgr1829Z12Z1839Z12Z15Z_h + udpgal_h --> mgdg1829Z12Z1839Z12Z15Z_h + udp_h | h_h | -2.000000 |
| MGDG1836Z9Z12Z1636Z9Z12Z_SYN | 12dgr1836Z9Z12Z1636Z9Z12Z_h + udpgal_h --> 2.0 h_h + mgdg1836Z9Z12Z1636Z9Z12Z_h + udp_h | 12dgr1836Z9Z12Z1636Z9Z12Z_h + udpgal_h --> mgdg1836Z9Z12Z1636Z9Z12Z_h + udp_h | h_h | -2.000000 |
| MGDG1839Z12Z15Z1629Z12Z_SYN | 12dgr1839Z12Z15Z1629Z12Z_h + udpgal_h --> 2.0 h_h + mgdg1839Z12Z15Z1629Z12Z_h + udp_h | 12dgr1839Z12Z15Z1629Z12Z_h + udpgal_h --> mgdg1839Z12Z15Z1629Z12Z_h + udp_h | h_h | -2.000000 |
| MGDG1839Z12Z15Z1636Z9Z12Z_SYN | 12dgr1839Z12Z15Z1636Z9Z12Z_h + udpgal_h --> 2.0 h_h + mgdg1839Z12Z15Z1636Z9Z12Z_h + udp_h | 12dgr1839Z12Z15Z1636Z9Z12Z_h + udpgal_h --> mgdg1839Z12Z15Z1636Z9Z12Z_h + udp_h | h_h | -2.000000 |
| MGDG1839Z12Z15Z1839Z12Z15Z_SYN | 12dgr1839Z12Z15Z1839Z12Z15Z_h + udpgal_h --> h_h + mgdg1839Z12Z15Z1839Z12Z15Z_h + udp_h | 12dgr1839Z12Z15Z1839Z12Z15Z_h + udpgal_h --> mgdg1839Z12Z15Z1839Z12Z15Z_h + udp_h | h_h | -1.000000 |
| MGDG2801426Z9Z_SYN | 12dgr2801426Z9Z_h + udpgal_h --> 2.0 h_h + mgdg2801426Z9Z_h + udp_h | 12dgr2801426Z9Z_h + udpgal_h --> mgdg2801426Z9Z_h + udp_h | h_h | -2.000000 |
| PCHOL1002239Z12Z15Z_SYN | 12dgr1002239Z12Z15Z_c + cdpchol_c --> cmp_c + h_c + pchol1002239Z12Z15Z_c | 12dgr1002239Z12Z15Z_c + cdpchol_c --> cmp_c + pchol1002239Z12Z15Z_c | h_c | -1.000000 |
| PCHOL10060_SYN | 12dgr10060_c + cdpchol_c --> cmp_c + h_c + pchol10060_c | 12dgr10060_c + cdpchol_c --> cmp_c + pchol10060_c | h_c | -1.000000 |
| PCHOL1102229Z12Z_SYN | 12dgr1102229Z12Z_c + cdpchol_c --> cmp_c + h_c + pchol1102229Z12Z_c | 12dgr1102229Z12Z_c + cdpchol_c --> cmp_c + pchol1102229Z12Z_c | h_c | -1.000000 |
| PCHOL1201839Z12Z15Z_SYN | 12dgr1201839Z12Z15Z_c + cdpchol_c --> cmp_c + h_c + pchol1201839Z12Z15Z_c | 12dgr1201839Z12Z15Z_c + cdpchol_c --> cmp_c + pchol1201839Z12Z15Z_c | h_c | -1.000000 |
| PCHOL12060_SYN | 12dgr12060_c + cdpchol_c --> cmp_c + h_c + pchol12060_c | 12dgr12060_c + cdpchol_c --> cmp_c + pchol12060_c | h_c | -1.000000 |
| PCHOL1219Z60_SYN | 12dgr1219Z60_c + cdpchol_c --> cmp_c + h_c + pchol1219Z60_c | 12dgr1219Z60_c + cdpchol_c --> cmp_c + pchol1219Z60_c | h_c | -1.000000 |
| PCHOL1402046Z9Z12Z15Z_SYN | 12dgr1402046Z9Z12Z15Z_c + cdpchol_c --> cmp_c + h_c + pchol1402046Z9Z12Z15Z_c | 12dgr1402046Z9Z12Z15Z_c + cdpchol_c --> cmp_c + pchol1402046Z9Z12Z15Z_c | h_c | -1.000000 |
| PCHOL1601619Z_SYN | 12dgr1601619Z_c + cdpchol_c --> cmp_c + h_c + pchol1601619Z_c | 12dgr1601619Z_c + cdpchol_c --> cmp_c + pchol1601619Z_c | h_c | -1.000000 |
| PCHOL1601819Z_SYN | 12dgr1601819Z_c + cdpchol_c --> cmp_c + h_c + pchol1601819Z_c | 12dgr1601819Z_c + cdpchol_c --> cmp_c + pchol1601819Z_c | h_c | -1.000000 |
| PCHOL1601826Z9Z_SYN | 12dgr1601826Z9Z_c + cdpchol_c --> cmp_c + h_c + pchol1601826Z9Z_c | 12dgr1601826Z9Z_c + cdpchol_c --> cmp_c + pchol1601826Z9Z_c | h_c | -1.000000 |
| PCHOL1601829Z12Z_SYN | 12dgr1601829Z12Z_c + cdpchol_c --> cmp_c + h_c + pchol1601829Z12Z_c | 12dgr1601829Z12Z_c + cdpchol_c --> cmp_c + pchol1601829Z12Z_c | h_c | -1.000000 |
| PCHOL1601836Z9Z12Z_SYN | 12dgr1601836Z9Z12Z_c + cdpchol_c --> cmp_c + h_c + pchol1601836Z9Z12Z_c | 12dgr1601836Z9Z12Z_c + cdpchol_c --> cmp_c + pchol1601836Z9Z12Z_c | h_c | -1.000000 |
| PCHOL17080_SYN | 12dgr17080_c + cdpchol_c --> cmp_c + h_c + pchol17080_c | 12dgr17080_c + cdpchol_c --> cmp_c + pchol17080_c | h_c | -1.000000 |
| PCHOL180140_SYN | 12dgr180140_c + cdpchol_c --> cmp_c + h_c + pchol180140_c | 12dgr180140_c + cdpchol_c --> cmp_c + pchol180140_c | h_c | -1.000000 |
| PCHOL1801426Z9Z_SYN | 12dgr1801426Z9Z_c + cdpchol_c --> cmp_c + h_c + pchol1801426Z9Z_c | 12dgr1801426Z9Z_c + cdpchol_c --> cmp_c + pchol1801426Z9Z_c | h_c | -1.000000 |
| PCHOL1801433Z6Z9Z_SYN | 12dgr1801433Z6Z9Z_c + cdpchol_c --> cmp_c + h_c + pchol1801433Z6Z9Z_c | 12dgr1801433Z6Z9Z_c + cdpchol_c --> cmp_c + pchol1801433Z6Z9Z_c | h_c | -1.000000 |
| PCHOL1801435Z8Z11Z_SYN | 12dgr1801435Z8Z11Z_c + cdpchol_c --> cmp_c + h_c + pchol1801435Z8Z11Z_c | 12dgr1801435Z8Z11Z_c + cdpchol_c --> cmp_c + pchol1801435Z8Z11Z_c | h_c | -1.000000 |
| PCHOL1801443Z6Z9Z12Z_SYN | 12dgr1801443Z6Z9Z12Z_c + cdpchol_c --> cmp_c + h_c + pchol1801443Z6Z9Z12Z_c | 12dgr1801443Z6Z9Z12Z_c + cdpchol_c --> cmp_c + pchol1801443Z6Z9Z12Z_c | h_c | -1.000000 |
| PCHOL1801819Z_SYN | 12dgr1801819Z_c + cdpchol_c --> cmp_c + h_c + pchol1801819Z_c | 12dgr1801819Z_c + cdpchol_c --> cmp_c + pchol1801819Z_c | h_c | -1.000000 |
| PCHOL1801829Z12Z_SYN | 12dgr1801829Z12Z_c + cdpchol_c --> cmp_c + h_c + pchol1801829Z12Z_c | 12dgr1801829Z12Z_c + cdpchol_c --> cmp_c + pchol1801829Z12Z_c | h_c | -1.000000 |
| PCHOL1801836Z9Z12Z_SYN | 12dgr1801836Z9Z12Z_c + cdpchol_c --> cmp_c + h_c + pchol1801836Z9Z12Z_c | 12dgr1801836Z9Z12Z_c + cdpchol_c --> cmp_c + pchol1801836Z9Z12Z_c | h_c | -1.000000 |
| PCHOL1801839Z12Z15Z_SYN | 12dgr1801839Z12Z15Z_c + cdpchol_c --> cmp_c + h_c + pchol1801839Z12Z15Z_c | 12dgr1801839Z12Z15Z_c + cdpchol_c --> cmp_c + pchol1801839Z12Z15Z_c | h_c | -1.000000 |
| PCHOL1802029Z12Z_SYN | 12dgr1802029Z12Z_c + cdpchol_c --> cmp_c + h_c + pchol1802029Z12Z_c | 12dgr1802029Z12Z_c + cdpchol_c --> cmp_c + pchol1802029Z12Z_c | h_c | -1.000000 |
| PCHOL1802053Z6Z9Z12Z15Z_SYN | 12dgr1802053Z6Z9Z12Z15Z_c + cdpchol_c --> cmp_c + h_c + pchol1802053Z6Z9Z12Z15Z_c | 12dgr1802053Z6Z9Z12Z15Z_c + cdpchol_c --> cmp_c + pchol1802053Z6Z9Z12Z15Z_c | h_c | -1.000000 |
| PCHOL18119Z1829Z12Z_SYN | 12dgr18119Z1829Z12Z_c + cdpchol_c --> cmp_c + h_c + pchol18119Z1829Z12Z_c | 12dgr18119Z1829Z12Z_c + cdpchol_c --> cmp_c + pchol18119Z1829Z12Z_c | h_c | -1.000000 |
| PCHOL1826Z9Z1826Z9Z_SYN | 12dgr1826Z9Z1826Z9Z_c + cdpchol_c --> cmp_c + h_c + pchol1826Z9Z1826Z9Z_c | 12dgr1826Z9Z1826Z9Z_c + cdpchol_c --> cmp_c + pchol1826Z9Z1826Z9Z_c | h_c | -1.000000 |
| PCHOL1829Z12Z1826Z9Z_SYN | 12dgr1829Z12Z1826Z9Z_c + cdpchol_c --> cmp_c + h_c + pchol1829Z12Z1826Z9Z_c | 12dgr1829Z12Z1826Z9Z_c + cdpchol_c --> cmp_c + pchol1829Z12Z1826Z9Z_c | h_c | -1.000000 |
| PCHOL1829Z12Z1829Z12Z_SYN | 12dgr1829Z12Z1829Z12Z_c + cdpchol_c --> cmp_c + h_c + pchol1829Z12Z1829Z12Z_c | 12dgr1829Z12Z1829Z12Z_c + cdpchol_c --> cmp_c + pchol1829Z12Z1829Z12Z_c | h_c | -1.000000 |
| PCHOL1839Z12Z15Z1829Z12Z_SYN | 12dgr1839Z12Z15Z1829Z12Z_c + cdpchol_c --> cmp_c + h_c + pchol1839Z12Z15Z1829Z12Z_c | 12dgr1839Z12Z15Z1829Z12Z_c + cdpchol_c --> cmp_c + pchol1839Z12Z15Z1829Z12Z_c | h_c | -1.000000 |
| PCHOL1839Z12Z15Z1839Z12Z15Z_SYN | 12dgr1839Z12Z15Z1839Z12Z15Z_c + cdpchol_c --> cmp_c + h_c + pchol1839Z12Z15Z1839Z12Z15Z_c | 12dgr1839Z12Z15Z1839Z12Z15Z_c + cdpchol_c --> cmp_c + pchol1839Z12Z15Z1839Z12Z15Z_c | h_c | -1.000000 |
| PCHOL1846Z9Z12Z15Z1819Z_SYN | 12dgr1846Z9Z12Z15Z1819Z_c + cdpchol_c --> cmp_c + h_c + pchol1846Z9Z12Z15Z1819Z_c | 12dgr1846Z9Z12Z15Z1819Z_c + cdpchol_c --> cmp_c + pchol1846Z9Z12Z15Z1819Z_c | h_c | -1.000000 |
| PCHOL2001426Z9Z_SYN | 12dgr2001426Z9Z_c + cdpchol_c --> cmp_c + h_c + pchol2001426Z9Z_c | 12dgr2001426Z9Z_c + cdpchol_c --> cmp_c + pchol2001426Z9Z_c | h_c | -1.000000 |
| PCHOL2001436Z9Z12Z_SYN | 12dgr2001436Z9Z12Z_c + cdpchol_c --> cmp_c + h_c + pchol2001436Z9Z12Z_c | 12dgr2001436Z9Z12Z_c + cdpchol_c --> cmp_c + pchol2001436Z9Z12Z_c | h_c | -1.000000 |
| PCHOL2226Z9Z1433Z6Z9Z_SYN | 12dgr2226Z9Z1433Z6Z9Z_c + cdpchol_c --> cmp_c + h_c + pchol2226Z9Z1433Z6Z9Z_c | 12dgr2226Z9Z1433Z6Z9Z_c + cdpchol_c --> cmp_c + pchol2226Z9Z1433Z6Z9Z_c | h_c | -1.000000 |
| PCHOL2229Z12Z1433Z6Z9Z_SYN | 12dgr2229Z12Z1433Z6Z9Z_c + cdpchol_c --> cmp_c + h_c + pchol2229Z12Z1433Z6Z9Z_c | 12dgr2229Z12Z1433Z6Z9Z_c + cdpchol_c --> cmp_c + pchol2229Z12Z1433Z6Z9Z_c | h_c | -1.000000 |
| PCHOL2401433Z6Z9Z_SYN | 12dgr2401433Z6Z9Z_c + cdpchol_c --> cmp_c + h_c + pchol2401433Z6Z9Z_c | 12dgr2401433Z6Z9Z_c + cdpchol_c --> cmp_c + pchol2401433Z6Z9Z_c | h_c | -1.000000 |
| PCHOL2419Z1216Z_SYN | 12dgr2419Z1216Z_c + cdpchol_c --> cmp_c + h_c + pchol2419Z1216Z_c | 12dgr2419Z1216Z_c + cdpchol_c --> cmp_c + pchol2419Z1216Z_c | h_c | -1.000000 |
| PCHOL2419Z1219Z_SYN | 12dgr2419Z1219Z_c + cdpchol_c --> cmp_c + h_c + pchol2419Z1219Z_c | 12dgr2419Z1219Z_c + cdpchol_c --> cmp_c + pchol2419Z1219Z_c | h_c | -1.000000 |
| PCHOL2419Z1433Z6Z9Z_SYN | 12dgr2419Z1433Z6Z9Z_c + cdpchol_c --> cmp_c + h_c + pchol2419Z1433Z6Z9Z_c | 12dgr2419Z1433Z6Z9Z_c + cdpchol_c --> cmp_c + pchol2419Z1433Z6Z9Z_c | h_c | -1.000000 |
| PCHOL2426Z9Z1433Z6Z9Z_SYN | 12dgr2426Z9Z1433Z6Z9Z_c + cdpchol_c --> cmp_c + h_c + pchol2426Z9Z1433Z6Z9Z_c | 12dgr2426Z9Z1433Z6Z9Z_c + cdpchol_c --> cmp_c + pchol2426Z9Z1433Z6Z9Z_c | h_c | -1.000000 |
| PCHOL2501233Z6Z9Z_SYN | 12dgr2501233Z6Z9Z_c + cdpchol_c --> cmp_c + h_c + pchol2501233Z6Z9Z_c | 12dgr2501233Z6Z9Z_c + cdpchol_c --> cmp_c + pchol2501233Z6Z9Z_c | h_c | -1.000000 |
| PCHOL2619Z80_SYN | 12dgr2619Z80_c + cdpchol_c --> cmp_c + h_c + pchol2619Z80_c | 12dgr2619Z80_c + cdpchol_c --> cmp_c + pchol2619Z80_c | h_c | -1.000000 |
| PCHOL2701133Z6Z9Z_SYN | 12dgr2701133Z6Z9Z_c + cdpchol_c --> cmp_c + h_c + pchol2701133Z6Z9Z_c | 12dgr2701133Z6Z9Z_c + cdpchol_c --> cmp_c + pchol2701133Z6Z9Z_c | h_c | -1.000000 |
| PCHOL813Z1023Z6Z_SYN | 12dgr813Z1023Z6Z_c + cdpchol_c --> cmp_c + h_c + pchol813Z1023Z6Z_c | 12dgr813Z1023Z6Z_c + cdpchol_c --> cmp_c + pchol813Z1023Z6Z_c | h_c | -1.000000 |
| PCHOL816Z1016Z_SYN | 12dgr816Z1016Z_c + cdpchol_c --> cmp_c + h_c + pchol816Z1016Z_c | 12dgr816Z1016Z_c + cdpchol_c --> cmp_c + pchol816Z1016Z_c | h_c | -1.000000 |
| PCHOL816Z1019Z_SYN | 12dgr816Z1019Z_c + cdpchol_c --> cmp_c + h_c + pchol816Z1019Z_c | 12dgr816Z1019Z_c + cdpchol_c --> cmp_c + pchol816Z1019Z_c | h_c | -1.000000 |
| PCHOL816Z1023Z6Z_SYN | 12dgr816Z1023Z6Z_c + cdpchol_c --> cmp_c + h_c + pchol816Z1023Z6Z_c | 12dgr816Z1023Z6Z_c + cdpchol_c --> cmp_c + pchol816Z1023Z6Z_c | h_c | -1.000000 |
| PCHOL819Z100_SYN | 12dgr819Z100_c + cdpchol_c --> cmp_c + h_c + pchol819Z100_c | 12dgr819Z100_c + cdpchol_c --> cmp_c + pchol819Z100_c | h_c | -1.000000 |
| PETHDMT[c] | amet_c + methamp_c <=> ahcys_c + ethadmp_c + h_c | amet_c + methamp_c <=> ahcys_c + ethadmp_c | h_c | -1.000000 |
| PETHMT[c] | amet_c + ethamp_c <=> ahcys_c + h_c + methamp_c | amet_c + ethamp_c <=> ahcys_c + methamp_c | h_c | -1.000000 |
| PETHTMT[c] | amet_c + ethadmp_c <=> ahcys_c + cholp_c + h_c | amet_c + ethadmp_c <=> ahcys_c + cholp_c | h_c | -1.000000 |
| PGLer | 6pgl_r + h2o_r --> 6pgc_r + h_r | 6pgl_r + h2o_r --> 6pgc_r | h_r | -1.000000 |
| PGPP100220H | h2o_h + pgp100220_h --> 2.0 h_h + pg100220_h + pi_h | h2o_h + pgp100220_h --> pg100220_h + pi_h | h_h | -2.000000 |
| PGPP1002419ZH | h2o_h + pgp1002419Z_h --> 2.0 h_h + pg1002419Z_h + pi_h | h2o_h + pgp1002419Z_h --> pg1002419Z_h + pi_h | h_h | -2.000000 |
| PGPP1202236Z9Z12ZH | h2o_h + pgp1202236Z9Z12Z_h --> 2.0 h_h + pg1202236Z9Z12Z_h + pi_h | h2o_h + pgp1202236Z9Z12Z_h --> 4.0 h_h + pg1202236Z9Z12Z_h + pi_h | h_h | 2.000000 |
| PGPP1202239Z12Z15ZH | h2o_h + pgp1202239Z12Z15Z_h --> 2.0 h_h + pg1202239Z12Z15Z_h + pi_h | h2o_h + pgp1202239Z12Z15Z_h --> 4.0 h_h + pg1202239Z12Z15Z_h + pi_h | h_h | 2.000000 |
| PGPP160160H | h2o_h + pgp160160_h --> 2.0 h_h + pg160160_h + pi_h | h2o_h + pgp160160_h --> pg160160_h + pi_h | h_h | -2.000000 |
| PGPP1601619ZH | h2o_h + pgp1601619Z_h --> 2.0 h_h + pg1601619Z_h + pi_h | h2o_h + pgp1601619Z_h --> pg1601619Z_h + pi_h | h_h | -2.000000 |
| PGPP1601819ZH | h2o_h + pgp1601819Z_h --> 2.0 h_h + pg1601819Z_h + pi_h | h2o_h + pgp1601819Z_h --> pg1601819Z_h + pi_h | h_h | -2.000000 |
| PGPP1601829Z12ZH | h2o_h + pgp1601829Z12Z_h --> 2.0 h_h + pg1601829Z12Z_h + pi_h | h2o_h + pgp1601829Z12Z_h --> pg1601829Z12Z_h + pi_h | h_h | -2.000000 |
| PGPP16018312Z15Z18ZH | h2o_h + pgp16018312Z15Z18Z_h --> 2.0 h_h + pg16018312Z15Z18Z_h + pi_h | h2o_h + pgp16018312Z15Z18Z_h --> pg16018312Z15Z18Z_h + pi_h | h_h | -2.000000 |
| PGPP16112Z1839Z12Z15ZH | h2o_h + pgp16112Z1839Z12Z15Z_h --> 2.0 h_h + pg16112Z1839Z12Z15Z_h + pi_h | h2o_h + pgp16112Z1839Z12Z15Z_h --> pg16112Z1839Z12Z15Z_h + pi_h | h_h | -2.000000 |
| PGPP1616Z1839Z12Z15ZH | h2o_h + pgp1616Z1839Z12Z15Z_h --> 2.0 h_h + pg1616Z1839Z12Z15Z_h + pi_h | h2o_h + pgp1616Z1839Z12Z15Z_h --> pg1616Z1839Z12Z15Z_h + pi_h | h_h | -2.000000 |
| PGPP1619Z1819ZH | h2o_h + pgp1619Z1819Z_h --> 2.0 h_h + pg1619Z1819Z_h + pi_h | h2o_h + pgp1619Z1819Z_h --> pg1619Z1819Z_h + pi_h | h_h | -2.000000 |
| PGPP1619Z1829Z12ZH | h2o_h + pgp1619Z1829Z12Z_h --> 2.0 h_h + pg1619Z1829Z12Z_h + pi_h | h2o_h + pgp1619Z1829Z12Z_h --> pg1619Z1829Z12Z_h + pi_h | h_h | -2.000000 |
| PGPP1619Z1839Z12Z15ZH | h2o_h + pgp1619Z1839Z12Z15Z_h --> 2.0 h_h + pg1619Z1839Z12Z15Z_h + pi_h | h2o_h + pgp1619Z1839Z12Z15Z_h --> pg1619Z1839Z12Z15Z_h + pi_h | h_h | -2.000000 |
| PGPP1719Z1716ZH | h2o_h + pgp1719Z1716Z_h --> 2.0 h_h + pg1719Z1716Z_h + pi_h | h2o_h + pgp1719Z1716Z_h --> pg1719Z1716Z_h + pi_h | h_h | -2.000000 |
| PGPP1719Z1719ZH | h2o_h + pgp1719Z1719Z_h --> 2.0 h_h + pg1719Z1719Z_h + pi_h | h2o_h + pgp1719Z1719Z_h --> pg1719Z1719Z_h + pi_h | h_h | -2.000000 |
| PGPP1801413ZH | h2o_h + pgp1801413Z_h --> 2.0 h_h + pg1801413Z_h + pi_h | h2o_h + pgp1801413Z_h --> pg1801413Z_h + pi_h | h_h | -2.000000 |
| PGPP1801416ZH | h2o_h + pgp1801416Z_h --> 2.0 h_h + pg1801416Z_h + pi_h | h2o_h + pgp1801416Z_h --> pg1801416Z_h + pi_h | h_h | -2.000000 |
| PGPP1801419ZH | h2o_h + pgp1801419Z_h --> 2.0 h_h + pg1801419Z_h + pi_h | h2o_h + pgp1801419Z_h --> pg1801419Z_h + pi_h | h_h | -2.000000 |
| PGPP180160H | h2o_h + pgp180160_h --> 2.0 h_h + pg180160_h + pi_h | h2o_h + pgp180160_h --> pg180160_h + pi_h | h_h | -2.000000 |
| PGPP1839Z12Z15Z1829Z12ZH | h2o_h + pgp1839Z12Z15Z1829Z12Z_h --> 2.0 h_h + pg1839Z12Z15Z1829Z12Z_h + pi_h | h2o_h + pgp1839Z12Z15Z1829Z12Z_h --> pg1839Z12Z15Z1829Z12Z_h + pi_h | h_h | -2.000000 |
| PGPP1839Z12Z15Z1839Z12Z15ZH | h2o_h + pgp1839Z12Z15Z1839Z12Z15Z_h --> 2.0 h_h + pg1839Z12Z15Z1839Z12Z15Z_h + pi_h | h2o_h + pgp1839Z12Z15Z1839Z12Z15Z_h --> pg1839Z12Z15Z1839Z12Z15Z_h + pi_h | h_h | -2.000000 |
| PGPS1202236Z9Z12ZH | cdp12dgr1202236Z9Z12Z_h + glyc3p_h --> cmp_h + pgp1202236Z9Z12Z_h | cdp12dgr1202236Z9Z12Z_h + glyc3p_h + 4.0 h_h --> cmp_h + pgp1202236Z9Z12Z_h | h_h | -4.000000 |
| PGPS1202239Z12Z15ZH | cdp12dgr1202239Z12Z15Z_h + glyc3p_h --> cmp_h + pgp1202239Z12Z15Z_h | cdp12dgr1202239Z12Z15Z_h + glyc3p_h + 4.0 h_h --> cmp_h + pgp1202239Z12Z15Z_h | h_h | -4.000000 |
| PGPS1719Z1829Z12ZH | cdp12dgr1719Z1829Z12Z_h + glyc3p_h --> cmp_h + pgp1719Z1829Z12Z_h | cdp12dgr1719Z1829Z12Z_h + glyc3p_h + 2.0 h_h --> cmp_h + pgp1719Z1829Z12Z_h | h_h | -2.000000 |
| TG120901119Z | 12dgr12090_h + acoa1119Z_h --> coa_h + tag120901119Z_h | 12dgr12090_h + acoa1119Z_h + 2.0 h_h --> coa_h + tag120901119Z_h | h_h | -2.000000 |
| TG1219Z901123Z6Z | 12dgr1219Z90_h + acoa1123Z6Z_h --> coa_h + tag1219Z901123Z6Z_h | 12dgr1219Z90_h + acoa1123Z6Z_h + 2.0 h_h --> coa_h + tag1219Z901123Z6Z_h | h_h | -2.000000 |
| TG1419Z9090 | 12dgr1419Z90_h + nonacoa_h --> coa_h + tag1419Z9090_h | 12dgr1419Z90_h + 2.0 h_h + nonacoa_h --> coa_h + tag1419Z9090_h | h_h | -2.000000 |
| TG1601602256Z9Z12Z15Z18Z | 12dgr160160_h + acoa2256Z9Z12Z15Z18Z_h --> coa_h + tag1601602256Z9Z12Z15Z18Z_h | 12dgr160160_h + acoa2256Z9Z12Z15Z18Z_h + 10.0 h_h --> coa_h + tag1601602256Z9Z12Z15Z18Z_h | h_h | -10.000000 |
| TG2001123Z6Z1719Z | 12dgr2001123Z6Z_h + acoa1719Z_h --> coa_h + tag2001123Z6Z1719Z_h | 12dgr2001123Z6Z_h + acoa1719Z_h --> coa_h + 4.0 h_h + tag2001123Z6Z1719Z_h | h_h | 4.000000 |
| TG2039Z12Z15Z1119Z2263Z6Z9Z12Z15Z18Z | 12dgr2039Z12Z15Z1119Z_h + acoa2263Z6Z9Z12Z15Z18Z_h --> coa_h + tag2039Z12Z15Z1119Z2263Z6Z9Z12Z15Z18Z_h | 12dgr2039Z12Z15Z1119Z_h + acoa2263Z6Z9Z12Z15Z18Z_h + 4.0 h_h --> coa_h + tag2039Z12Z15Z1119Z2263Z6Z9Z12Z15Z18Z_h | h_h | -4.000000 |
| TG2039Z12Z15Z1123Z6Z2263Z6Z9Z12Z15Z18Z | 12dgr2039Z12Z15Z1123Z6Z_h + acoa2263Z6Z9Z12Z15Z18Z_h --> coa_h + tag2039Z12Z15Z1123Z6Z2263Z6Z9Z12Z15Z18Z_h | 12dgr2039Z12Z15Z1123Z6Z_h + acoa2263Z6Z9Z12Z15Z18Z_h + 4.0 h_h --> coa_h + tag2039Z12Z15Z1123Z6Z2263Z6Z9Z12Z15Z18Z_h | h_h | -4.000000 |
| TG2039Z12Z15Z1123Z6Z2319Z | 12dgr2039Z12Z15Z1123Z6Z_h + acoa2319Z_h --> coa_h + tag2039Z12Z15Z1123Z6Z2319Z_h | 12dgr2039Z12Z15Z1123Z6Z_h + acoa2319Z_h --> coa_h + 4.0 h_h + tag2039Z12Z15Z1123Z6Z2319Z_h | h_h | 4.000000 |
| TG2046Z9Z12Z15Z1302029Z12Z | 12dgr2046Z9Z12Z15Z130_h + acoa2029Z12Z_h --> coa_h + tag2046Z9Z12Z15Z1302029Z12Z_h | 12dgr2046Z9Z12Z15Z130_h + acoa2029Z12Z_h + 4.0 h_h --> coa_h + tag2046Z9Z12Z15Z1302029Z12Z_h | h_h | -4.000000 |
| TG2053Z6Z9Z12Z15Z1123Z6Z2219Z | 12dgr2053Z6Z9Z12Z15Z1123Z6Z_h + acoa2219Z_h --> coa_h + tag2053Z6Z9Z12Z15Z1123Z6Z2219Z_h | 12dgr2053Z6Z9Z12Z15Z1123Z6Z_h + acoa2219Z_h + 4.0 h_h --> coa_h + tag2053Z6Z9Z12Z15Z1123Z6Z2219Z_h | h_h | -4.000000 |
| TG2053Z6Z9Z12Z15Z1123Z6Z2319Z | 12dgr2053Z6Z9Z12Z15Z1123Z6Z_h + acoa2319Z_h --> coa_h + tag2053Z6Z9Z12Z15Z1123Z6Z2319Z_h | 12dgr2053Z6Z9Z12Z15Z1123Z6Z_h + acoa2319Z_h + 4.0 h_h --> coa_h + tag2053Z6Z9Z12Z15Z1123Z6Z2319Z_h | h_h | -4.000000 |
| TG2101219Z1219Z | 12dgr2101219Z_h + acoa1219Z_h --> coa_h + tag2101219Z1219Z_h | 12dgr2101219Z_h + acoa1219Z_h + 4.0 h_h --> coa_h + tag2101219Z1219Z_h | h_h | -4.000000 |
| TG2256Z9Z12Z15Z18Z1123Z6Z2119Z | 12dgr2256Z9Z12Z15Z18Z1123Z6Z_h + acoa2119Z_h --> coa_h + tag2256Z9Z12Z15Z18Z1123Z6Z2119Z_h | 12dgr2256Z9Z12Z15Z18Z1123Z6Z_h + acoa2119Z_h + 4.0 h_h --> coa_h + tag2256Z9Z12Z15Z18Z1123Z6Z2119Z_h | h_h | -4.000000 |
| TG2301219Z1219Z | 12dgr2301219Z_h + acoa1219Z_h --> coa_h + tag2301219Z1219Z_h | 12dgr2301219Z_h + acoa1219Z_h + 4.0 h_h --> coa_h + tag2301219Z1219Z_h | h_h | -4.000000 |
| TG2429Z12Z1123Z6Z1419Z | 12dgr2429Z12Z1123Z6Z_h + acoa1419Z_h --> coa_h + tag2429Z12Z1123Z6Z1419Z_h | 12dgr2429Z12Z1123Z6Z_h + acoa1419Z_h + 10.0 h_h --> coa_h + tag2429Z12Z1123Z6Z1419Z_h | h_h | -10.000000 |
| TG2701123Z6Z1123Z6Z | 12dgr2701123Z6Z_h + acoa1123Z6Z_h --> coa_h + tag2701123Z6Z1123Z6Z_h | 12dgr2701123Z6Z_h + acoa1123Z6Z_h + 2.0 h_h --> coa_h + tag2701123Z6Z1123Z6Z_h | h_h | -2.000000 |
| TG2719Z60110 | 12dgr2719Z60_h + undeccoa_h --> coa_h + tag2719Z60110_h | 12dgr2719Z60_h + undeccoa_h --> coa_h + 4.0 h_h + tag2719Z60110_h | h_h | 4.000000 |
| TG2801223Z6Z1223Z6Z | 12dgr2801223Z6Z_h + acoa1223Z6Z_h --> coa_h + tag2801223Z6Z1223Z6Z_h | 12dgr2801223Z6Z_h + acoa1223Z6Z_h + 2.0 h_h --> coa_h + tag2801223Z6Z1223Z6Z_h | h_h | -2.000000 |
| TG2801223Z6Z1226Z9Z | 12dgr2801223Z6Z_h + acoa1226Z9Z_h --> coa_h + tag2801223Z6Z1226Z9Z_h | 12dgr2801223Z6Z_h + acoa1226Z9Z_h + 2.0 h_h --> coa_h + tag2801223Z6Z1226Z9Z_h | h_h | -2.000000 |
| TG2801226Z9Z1226Z9Z | 12dgr2801226Z9Z_h + acoa1226Z9Z_h --> coa_h + tag2801226Z9Z1226Z9Z_h | 12dgr2801226Z9Z_h + acoa1226Z9Z_h + 2.0 h_h --> coa_h + tag2801226Z9Z1226Z9Z_h | h_h | -2.000000 |
| TG28112Z1123Z6Z1123Z6Z | 12dgr28112Z1123Z6Z_h + acoa1123Z6Z_h --> coa_h + tag28112Z1123Z6Z1123Z6Z_h | 12dgr28112Z1123Z6Z_h + acoa1123Z6Z_h + 2.0 h_h --> coa_h + tag28112Z1123Z6Z1123Z6Z_h | h_h | -2.000000 |
| TG2819Z1001123Z6Z | 12dgr2819Z100_h + acoa1123Z6Z_h --> coa_h + tag2819Z1001123Z6Z_h | 12dgr2819Z100_h + acoa1123Z6Z_h + 2.0 h_h --> coa_h + tag2819Z1001123Z6Z_h | h_h | -2.000000 |
| TG2819Z1123Z6Z1123Z6Z | 12dgr2819Z1123Z6Z_h + acoa1123Z6Z_h --> coa_h + tag2819Z1123Z6Z1123Z6Z_h | 12dgr2819Z1123Z6Z_h + acoa1123Z6Z_h + 2.0 h_h --> coa_h + tag2819Z1123Z6Z1123Z6Z_h | h_h | -2.000000 |
| TG2819Z1216Z1233Z6Z9Z | 12dgr2819Z1216Z_h + acoa1233Z6Z9Z_h --> coa_h + tag2819Z1216Z1233Z6Z9Z_h | 12dgr2819Z1216Z_h + acoa1233Z6Z9Z_h + 2.0 h_h --> coa_h + tag2819Z1216Z1233Z6Z9Z_h | h_h | -2.000000 |
| TG2901016Z1123Z6Z | 12dgr2901016Z_h + acoa1123Z6Z_h --> coa_h + tag2901016Z1123Z6Z_h | 12dgr2901016Z_h + acoa1123Z6Z_h + 2.0 h_h --> coa_h + tag2901016Z1123Z6Z_h | h_h | -2.000000 |
| TG2901023Z6Z1123Z6Z | 12dgr2901023Z6Z_h + acoa1123Z6Z_h --> coa_h + tag2901023Z6Z1123Z6Z_h | 12dgr2901023Z6Z_h + acoa1123Z6Z_h + 2.0 h_h --> coa_h + tag2901023Z6Z1123Z6Z_h | h_h | -2.000000 |
| TG2901123Z6Z1123Z6Z | 12dgr2901123Z6Z_h + acoa1123Z6Z_h --> coa_h + tag2901123Z6Z1123Z6Z_h | 12dgr2901123Z6Z_h + acoa1123Z6Z_h + 2.0 h_h --> coa_h + tag2901123Z6Z1123Z6Z_h | h_h | -2.000000 |
| TG2901123Z6Z1126Z9Z | 12dgr2901123Z6Z_h + acoa1126Z9Z_h --> coa_h + tag2901123Z6Z1126Z9Z_h | 12dgr2901123Z6Z_h + acoa1126Z9Z_h + 2.0 h_h --> coa_h + tag2901123Z6Z1126Z9Z_h | h_h | -2.000000 |
| TG2901123Z6Z1133Z6Z9Z | 12dgr2901123Z6Z_h + acoa1133Z6Z9Z_h --> coa_h + tag2901123Z6Z1133Z6Z9Z_h | 12dgr2901123Z6Z_h + acoa1133Z6Z9Z_h + 2.0 h_h --> coa_h + tag2901123Z6Z1133Z6Z9Z_h | h_h | -2.000000 |
| TG2901123Z6Z140 | 12dgr2901123Z6Z_h + tdcoa_h --> coa_h + tag2901123Z6Z140_h | 12dgr2901123Z6Z_h + 2.0 h_h + tdcoa_h --> coa_h + tag2901123Z6Z140_h | h_h | -2.000000 |
| TG2901123Z6Z1416Z | 12dgr2901123Z6Z_h + acoa1416Z_h --> coa_h + tag2901123Z6Z1416Z_h | 12dgr2901123Z6Z_h + acoa1416Z_h + 2.0 h_h --> coa_h + tag2901123Z6Z1416Z_h | h_h | -2.000000 |
| TG2901123Z6Z1419Z | 12dgr2901123Z6Z_h + acoa1419Z_h --> coa_h + tag2901123Z6Z1419Z_h | 12dgr2901123Z6Z_h + acoa1419Z_h + 2.0 h_h --> coa_h + tag2901123Z6Z1419Z_h | h_h | -2.000000 |
| TG2901123Z6Z1426Z9Z | 12dgr2901123Z6Z_h + acoa1426Z9Z_h --> coa_h + tag2901123Z6Z1426Z9Z_h | 12dgr2901123Z6Z_h + acoa1426Z9Z_h + 2.0 h_h --> coa_h + tag2901123Z6Z1426Z9Z_h | h_h | -2.000000 |
| TG2901123Z6Z1433Z6Z9Z | 12dgr2901123Z6Z_h + acoa1433Z6Z9Z_h --> coa_h + tag2901123Z6Z1433Z6Z9Z_h | 12dgr2901123Z6Z_h + acoa1433Z6Z9Z_h + 2.0 h_h --> coa_h + tag2901123Z6Z1433Z6Z9Z_h | h_h | -2.000000 |
| TG3001123Z6Z1123Z6Z | 12dgr3001123Z6Z_h + acoa1123Z6Z_h --> coa_h + tag3001123Z6Z1123Z6Z_h | 12dgr3001123Z6Z_h + acoa1123Z6Z_h + 2.0 h_h --> coa_h + tag3001123Z6Z1123Z6Z_h | h_h | -2.000000 |
| TG3001123Z6Z1126Z9Z | 12dgr3001123Z6Z_h + acoa1126Z9Z_h --> coa_h + tag3001123Z6Z1126Z9Z_h | 12dgr3001123Z6Z_h + acoa1126Z9Z_h + 2.0 h_h --> coa_h + tag3001123Z6Z1126Z9Z_h | h_h | -2.000000 |
| TG3001226Z9Z1226Z9Z | 12dgr3001226Z9Z_h + acoa1226Z9Z_h --> coa_h + tag3001226Z9Z1226Z9Z_h | 12dgr3001226Z9Z_h + acoa1226Z9Z_h + 2.0 h_h --> coa_h + tag3001226Z9Z1226Z9Z_h | h_h | -2.000000 |
| TG3019Z1226Z9Z1226Z9Z | 12dgr3019Z1226Z9Z_h + acoa1226Z9Z_h --> coa_h + tag3019Z1226Z9Z1226Z9Z_h | 12dgr3019Z1226Z9Z_h + acoa1226Z9Z_h + 2.0 h_h --> coa_h + tag3019Z1226Z9Z1226Z9Z_h | h_h | -2.000000 |
| TG402002119Z | 12dgr40200_h + acoa2119Z_h --> coa_h + tag402002119Z_h | 12dgr40200_h + acoa2119Z_h + 2.0 h_h --> coa_h + tag402002119Z_h | h_h | -2.000000 |
| TG402029Z12Z220 | 12dgr402029Z12Z_h + docoscoa_h --> coa_h + tag402029Z12Z220_h | 12dgr402029Z12Z_h + docoscoa_h + 2.0 h_h --> coa_h + tag402029Z12Z220_h | h_h | -2.000000 |
| TG402036Z9Z12Z240 | 12dgr402036Z9Z12Z_h + ttccoa_h --> coa_h + tag402036Z9Z12Z240_h | 12dgr402036Z9Z12Z_h + 2.0 h_h + ttccoa_h --> coa_h + tag402036Z9Z12Z240_h | h_h | -2.000000 |
| TG402302319Z | 12dgr40230_h + acoa2319Z_h --> coa_h + tag402302319Z_h | 12dgr40230_h + acoa2319Z_h + 2.0 h_h --> coa_h + tag402302319Z_h | h_h | -2.000000 |
| TG402419Z2419Z | 12dgr402419Z_h + acoa2419Z_h --> coa_h + tag402419Z2419Z_h | 12dgr402419Z_h + acoa2419Z_h + 2.0 h_h --> coa_h + tag402419Z2419Z_h | h_h | -2.000000 |
| TG901433Z6Z9Z1433Z6Z9Z | 12dgr901433Z6Z9Z_h + acoa1433Z6Z9Z_h --> coa_h + tag901433Z6Z9Z1433Z6Z9Z_h | 12dgr901433Z6Z9Z_h + acoa1433Z6Z9Z_h + 6.0 h_h --> coa_h + tag901433Z6Z9Z1433Z6Z9Z_h | h_h | -6.000000 |
| SSALxm | h2o_m + nad_m + sucsal_m <=> 2.0 h_m + nadh_m + succ_m | h2o_m + nad_m + sucsal_m --> nadh_m + succ_m | h_m | -2.000000 |
| MTHFC | h2o_c + methf_c <=> 10fthf_c + h_c | h2o_c + methf_c --> 10fthf_c | h_c | -1.000000 |
| PDHe3mr | dhlam_m + nad_m --> h_m + lpam_m + nadh_m | dhlam_m + nad_m --> lpam_m + nadh_m | h_m | -1.000000 |
| ALCDH_nadp_i | acald_c + h_c + nadph_c --> etoh_c + nadp_c | acald_c + nadph_c --> etoh_c + nadp_c | h_c | 1.000000 |
| NanoG1180 | 3oodcoa_c + h_c + nadph_c --> 3hodcoa_c + nadp_c | 3oodcoa_c + nadph_c --> 3hodcoa_c + nadp_c | h_c | 1.000000 |
| NanoG1707 | co2_h + h2o_h + pep_h --> h_h + oaa_h + pi_h | co2_h + h2o_h + pep_h --> oaa_h + pi_h | h_h | -1.000000 |
| CY_focytc_h | ficytc_h --> focytc_h | ficytc_h + h_h --> focytc_h | h_h | -1.000000 |
| NanoG0278 | 3hib_m + nad_m --> h_m + mmsa_m + nadh_m | 3hib_m + nad_m --> mmsa_m + nadh_m | h_m | -1.000000 |
| PCr | atp_c + hco3_c + pyr_c --> adp_c + h_c + oaa_c + pi_c | atp_c + hco3_c + pyr_c --> adp_c + oaa_c + pi_c | h_c | -1.000000 |
| NanoG0263 | 2ahbut_h + h_h + nadph_h --> 23dhmp_h + nadp_h | 2ahbut_h + nadph_h --> 23dhmp_h + nadp_h | h_h | 1.000000 |
| NanoG1289 | glutrna_h + h_h + nadph_h --> glu1sa_h + nadp_h + trnaglu_h | glutrna_h + nadph_h --> glu1sa_h + nadp_h + trnaglu_h | h_h | 1.000000 |
| GLYCLTD | glx_c + h_c + nadh_c --> glyclt_c + nad_c | glx_c + nadh_c --> glyclt_c + nad_c | h_c | 1.000000 |
| NanoG1233 | h2o_c + methf_c --> 10fthf_c + h_c | h2o_c + methf_c --> 10fthf_c | h_c | -1.000000 |
| NanoG1034 | dhap_c + h_c + nadh_c --> glyc3p_c + nad_c | dhap_c + nadh_c --> glyc3p_c + nad_c | h_c | 1.000000 |
| NanoG0606 | adp_m + 3.0 h_c + pi_m --> atp_m + h2o_m + 2.0 h_m | adp_m + 2.0 h_c + pi_m --> atp_m + h2o_m + 2.0 h_m | h_c | 1.000000 |
| TMDPT | atp_c + thm_c --> amp_c + h_c + thmpp_c | atp_c + thm_c --> amp_c + thmpp_c | h_c | -1.000000 |
| NanoG1115 | amet_c + cyart_c --> ahcys_c + h_c + menecyart_c | amet_c + cyart_c --> ahcys_c + menecyart_c | h_c | -1.000000 |
| NanoG0609 | h2o_h + ppi_h --> h_h + 2.0 pi_h | h2o_h + ppi_h --> 2.0 pi_h | h_h | -1.000000 |
| NanoG1101 | 2.0 h_c + lanost_c + 3.0 nadph_c + 3.0 o2_c --> 44mctr_c + for_c + 4.0 h2o_c + 3.0 nadp_c | lanost_c + 3.0 nadph_c + 3.0 o2_c --> 44mctr_c + for_c + 4.0 h2o_c + 3.0 nadp_c | h_c | 2.000000 |
| NanoG0326 | acgam1p_c + h_c + utp_c --> ppi_c + uacgam_c | acgam1p_c + utp_c --> ppi_c + uacgam_c | h_c | 1.000000 |
| NanoG1582 | 2.0 focytc_m + h2o2_m + 2.0 h_m --> 2.0 ficytc_m + 2.0 h2o_m | 2.0 focytc_m + h2o2_m --> 2.0 ficytc_m + 2.0 h2o_m | h_m | 2.000000 |
| NanoG0832 | 3ohcoa_m + h_m + nadh_m --> 3hhcoa_m + nad_m | 3ohcoa_m + nadh_m --> 3hhcoa_m + nad_m | h_m | 1.000000 |
| NanoG0021 | h2o_h + nadp_h + sucsal_h --> 2.0 h_h + nadph_h + succ_h | h2o_h + nadp_h + sucsal_h --> nadph_h + succ_h | h_h | -2.000000 |
| NanoG0460 | o2_h + rb15bp_h --> 2pglyc_h + 3pg_h + 2.0 h_h | o2_h + rb15bp_h --> 2pglyc_h + 3pg_h + h_h | h_h | -1.000000 |
| NanoG0449 | acald_m + h2o_m + nadp_m --> ac_m + 2.0 h_m + nadph_m | acald_m + h2o_m + nadp_m --> ac_m + nadph_m | h_m | -2.000000 |
| NanoG0458 | glx_c + h_c + nadph_c --> glyclt_c + nadp_c | glx_c + nadph_c --> glyclt_c + nadp_c | h_c | 1.000000 |
| NanoG0405 | 2pg_c + h_c --> h2o_c + pep_c | 2pg_c --> h2o_c + pep_c | h_c | 1.000000 |
| NanoG1452 | h2o_c + nad_c + xan_c --> h_c + nadh_c + urate_c | h2o_c + nad_c + xan_c --> nadh_c + urate_c | h_c | -1.000000 |
| MTHFO_nadp | h_m + mlthf_m + nadph_m --> 5mthf_m + nadp_m | mlthf_m + nadph_m --> 5mthf_m + nadp_m | h_m | 1.000000 |
| GDR_nadp_m | gthox_m + h_m + nadph_m --> 2.0 gthrd_m + nadp_m | gthox_m + nadph_m --> 2.0 gthrd_m + nadp_m | h_m | 1.000000 |
| NanoG0052 | h_m + orn_m --> co2_m + ptrc_m | orn_m --> co2_m + ptrc_m | h_m | 1.000000 |
| NanoG0268 | 3mb2coa_m + atp_m + hco3_m --> 3mgcoa_m + adp_m + h_m + pi_m | 3mb2coa_m + atp_m + hco3_m --> 3mgcoa_m + adp_m + pi_m | h_m | -1.000000 |
| NanoG0332 | 2.0 ficytb5_c + nadh_c --> 2.0 focytb5_c + h_c + nad_c | 2.0 ficytb5_c + nadh_c --> 2.0 focytb5_c + 2.0 h_c + nad_c | h_c | 1.000000 |
| NanoG1109 | chlstol_c + h_c + nadph_c + o2_c --> ddsmsterol_c + 2.0 h2o_c + nadp_c | chlstol_c + nadph_c + o2_c --> ddsmsterol_c + 2.0 h2o_c + nadp_c | h_c | 1.000000 |
| ACP | 13dpg_c + h2o_c --> 3pg_c + h_c + pi_c | 13dpg_c + h2o_c --> 3pg_c + pi_c | h_c | -1.000000 |
| NanoG1345 | amet_h + yvite_h --> ahcys_h + avite1_h + h_h | amet_h + yvite_h --> ahcys_h + avite1_h | h_h | -1.000000 |
| NanoG0459 | glyclt_m + nad_m --> glx_m + h_m + nadh_m | glyclt_m + nad_m --> glx_m + nadh_m | h_m | -1.000000 |
| ATNS_nh4 | chor_h + nh4_h --> anth_h + h2o_h + h_h + pyr_h | chor_h + nh4_h --> anth_h + h2o_h + pyr_h | h_h | -1.000000 |
| NanoG0608 | 4.0 focytc_m + 4.0 h_m + o2_m --> 4.0 ficytc_m + 2.0 h2o_m | 4.0 focytc_m + o2_m --> 4.0 ficytc_m + 2.0 h2o_m | h_m | 4.000000 |
| HOXGh | 3.0 h_h + 3.0 nadph_h + 3.0 o2_h + pheme_h --> biliverd_h + co_h + fe2_h + 3.0 h2o_h + 3.0 nadp_h | 2.0 h_h + 3.0 nadph_h + 3.0 o2_h + pheme_h --> biliverd_h + co_h + fe2_h + 3.0 h2o_h + 3.0 nadp_h | h_h | 1.000000 |
| NanoG1104 | h_h + hgentis_h + pdp_h --> co2_h + mpq_h + ppi_h | hgentis_h + pdp_h --> co2_h + mpq_h + ppi_h | h_h | 1.000000 |
| NanoG1270 | h_h + hmppp9me_h + nadph_h + o2_h --> 2.0 h2o_h + nadp_h + omppp9me_h | hmppp9me_h + nadph_h + o2_h --> 2.0 h2o_h + nadp_h + omppp9me_h | h_h | 1.000000 |
| NanoG0544 | atp_h + pi_h + pyr_h --> amp_h + h_h + pep_h + ppi_h | atp_h + pi_h + pyr_h --> amp_h + pep_h + ppi_h | h_h | -1.000000 |
| NanoG1428 | 2mecdp_h + 2.0 fdxrd_h + h_h --> 2.0 fdxox_h + h2mb4p_h + h2o_h | 2mecdp_h + 2.0 fdxrd_h + 2.0 h_h --> 2.0 fdxox_h + h2mb4p_h + h2o_h | h_h | -1.000000 |
| NADHOR_2h | h_h + nadh_h + pq_h --> nad_h + pqh2_h | nadh_h + pq_h --> nad_h + pqh2_h | h_h | 1.000000 |
| NanoG0435 | cdp_m + h_m + pep_m --> ctp_m + pyr_m | cdp_m + pep_m --> ctp_m + pyr_m | h_m | 1.000000 |
| NanoG1111 | ddsmsterol_c + h_c + nadph_c --> dsmsterol_c + nadp_c | ddsmsterol_c + nadph_c --> dsmsterol_c + nadp_c | h_c | 1.000000 |
| MOD_4mop | 4mop_m + h_m + thmpp_m --> 3mhtpp_m + co2_m | 4mop_m + thmpp_m --> 3mhtpp_m + co2_m | h_m | 1.000000 |
| NanoG0261 | 2obut_h + h_h + pyr_h --> 2ahbut_h + co2_h | 2obut_h + pyr_h --> 2ahbut_h + co2_h | h_h | 1.000000 |
| NanoG0810 | 3hbcoa_m + nad_m --> aacoa_m + h_m + nadh_m | 3hbcoa_m + nad_m --> aacoa_m + nadh_m | h_m | -1.000000 |
| NanoG0450 | h_c + pyr_c --> acald_c + co2_c | pyr_c --> acald_c + co2_c | h_c | 1.000000 |
| NanoG0596 | fald_c + h2o_c + nad_c --> for_c + 2.0 h_c + nadh_c | fald_c + h2o_c + nad_c --> for_c + nadh_c | h_c | -2.000000 |
| NanoG0437 | 13dpg_h + h_h + nadh_h --> g3p_h + nad_h + pi_h | 13dpg_h + nadh_h --> g3p_h + nad_h + pi_h | h_h | 1.000000 |
| NanoG1501 | 5hiu_m + h2o_m --> 5houdic_m + h_m | 5hiu_m + h2o_m --> 5houdic_m | h_m | -1.000000 |
| ALDH_im4ac | h2o_c + im4act_c + nad_c --> 2.0 h_c + im4ac_c + nadh_c | h2o_c + im4act_c + nad_c --> im4ac_c + nadh_c | h_c | -2.000000 |
| NanoG1293 | cpppg3_h + 2.0 h_h + o2_h --> 2.0 co2_h + 2.0 h2o_h + pppg9_h | cpppg3_h + o2_h --> 2.0 co2_h + 2.0 h2o_h + pppg9_h | h_h | 2.000000 |
| GDR_nadp_h | gthox_h + h_h + nadph_h --> 2.0 gthrd_h + nadp_h | gthox_h + nadph_h --> 2.0 gthrd_h + nadp_h | h_h | 1.000000 |
| ACDO_co | 12dmpo_c + o2_c --> 3mtpa_c + co_c + for_c + h_c | 12dmpo_c + o2_c --> 3mtpa_c + co_c + for_c | h_c | -1.000000 |
| NanoG1224 | atp_h + nad_h --> adp_h + h_h + nadp_h | atp_h + nad_h --> adp_h + nadp_h | h_h | -1.000000 |
| NanoG1110 | 7dhchsterol_c + h_c + nadph_c --> chsterol_c + nadp_c | 7dhchsterol_c + nadph_c --> chsterol_c + nadp_c | h_c | 1.000000 |
| SULRi | 3.0 h_c + hso3_c + 3.0 nadph_c --> 3.0 h2o_c + h2s_c + 3.0 nadp_c | hso3_c + 3.0 nadph_c --> 3.0 h2o_c + h2s_c + 3.0 nadp_c | h_c | 3.000000 |
| HBCO_nadp_m | aacoa_m + h_m + nadph_m --> 3hbcoa_m + nadp_m | aacoa_m + nadph_m --> 3hbcoa_m + nadp_m | h_m | 1.000000 |
| GDR_nadp | gthox_c + h_c + nadph_c --> 2.0 gthrd_c + nadp_c | gthox_c + nadph_c --> 2.0 gthrd_c + nadp_c | h_c | 1.000000 |
| NanoG1598 | 2.0 fdxrd_m + 3.0 h_m + nadh_m --> 2.0 fdxox_m + 2.0 h2_m + nad_m | 2.0 fdxrd_m + 2.0 h_m + nadh_m --> 2.0 fdxox_m + 2.0 h2_m + nad_m | h_m | 1.000000 |
| NanoG1296 | chlda_h + h_h + pdp_h --> chla_h + ppi_h | chlda_h + pdp_h --> chla_h + ppi_h | h_h | 1.000000 |
| NanoG0613 | atp_c + h2o_c + h_c --> adp_c + 2.0 h_e + pi_c | atp_c + h2o_c + 2.0 h_c --> adp_c + 2.0 h_e + pi_c | h_c | -1.000000 |
| NanoG0431 | gdp_h + h_h + pep_h --> gtp_h + pyr_h | gdp_h + pep_h --> gtp_h + pyr_h | h_h | 1.000000 |
| NanoG1344 | amet_h + dvite_h --> ahcys_h + bvite_h + h_h | amet_h + dvite_h --> ahcys_h + bvite_h | h_h | -1.000000 |
| ALCDH_nadp_hi | acald_h + h_h + nadph_h --> etoh_h + nadp_h | acald_h + nadph_h --> etoh_h + nadp_h | h_h | 1.000000 |
| KARI_23dhmb | 3hmoa_h + h_h + nadph_h --> 23dhmb_h + nadp_h | 3hmoa_h + nadph_h --> 23dhmb_h + nadp_h | h_h | 1.000000 |
| NanoG1597 | 2.0 fdxrd_h + 3.0 h_h + nadh_h --> 2.0 fdxox_h + 2.0 h2_h + nad_h | 2.0 fdxrd_h + 2.0 h_h + nadh_h --> 2.0 fdxox_h + 2.0 h2_h + nad_h | h_h | 1.000000 |
| NanoG0600 | focytc_h + 2.0 h_h + no2_h --> ficytc_h + h2o_h + no_h | focytc_h + no2_h --> ficytc_h + h2o_h + no_h | h_h | 2.000000 |
| NanoG1172 | h_c + nadph_c + o2_c + ocdce9coa_c --> 2.0 h2o_c + lnlccoa_c + nadp_c | nadph_c + o2_c + ocdce9coa_c --> 2.0 h2o_c + lnlccoa_c + nadp_c | h_c | 1.000000 |
| NanoG0440 | acald_h + h_h + nadh_h --> etoh_h + nad_h | acald_h + nadh_h --> etoh_h + nad_h | h_h | 1.000000 |
| KARI_23dhmp | 3hmop_h + h_h + nadph_h --> 23dhmp_h + nadp_h | 3hmop_h + nadph_h --> 23dhmp_h + nadp_h | h_h | 1.000000 |
| NanoG0785 | acACP_h + h_h + malACP_h --> ACP_h + actACP_h + co2_h | acACP_h + malACP_h --> ACP_h + actACP_h + co2_h | h_h | 1.000000 |
| NanoG0030 | acg5p_c + h_c + nadph_c --> acg5sa_c + nadp_c + pi_c | acg5p_c + nadph_c --> acg5sa_c + nadp_c + pi_c | h_c | 1.000000 |
| MTHFOR_nadp | h_c + mlthf_c + nadp_c --> methf_c + nadph_c | mlthf_c + nadp_c --> methf_c + nadph_c | h_c | 1.000000 |
| NanoG0407 | 2pg_m + h_m --> h2o_m + pep_m | 2pg_m --> h2o_m + pep_m | h_m | 1.000000 |
| SSNOm | h2o_m + nad_m + sucsal_m --> 2.0 h_m + nadh_m + succ_m | h2o_m + nad_m + sucsal_m --> nadh_m + succ_m | h_m | -2.000000 |
| MOD_3mop | 3mop_m + h_m + thmpp_m --> 2mhob_m + co2_m | 3mop_m + thmpp_m --> 2mhob_m + co2_m | h_m | 1.000000 |
| DHDH | 56dura_c + nadp_c --> h_c + nadph_c + ura_c | 56dura_c + nadp_c --> nadph_c + ura_c | h_c | -1.000000 |
| NanoG1232 | 5fthf_m + h_m --> h2o_m + methf_m | 5fthf_m --> h2o_m + methf_m | h_m | 1.000000 |
| BTS_nadph | btal_c + h_c + nadph_c --> 1btol_c + nadp_c | btal_c + nadph_c --> 1btol_c + nadp_c | h_c | 1.000000 |
| NanoG1225 | atp_m + nad_m --> adp_m + h_m + nadp_m | atp_m + nad_m --> adp_m + nadp_m | h_m | -1.000000 |
| NanoG0433 | cdp_c + h_c + pep_c --> ctp_c + pyr_c | cdp_c + pep_c --> ctp_c + pyr_c | h_c | 1.000000 |
| NanoG1231 | dhf_c + h_c + nadph_c --> nadp_c + thf_c | dhf_c + nadph_c --> nadp_c + thf_c | h_c | 1.000000 |
| NanoG0448 | acald_c + h2o_c + nadp_c --> ac_c + 2.0 h_c + nadph_c | acald_c + h2o_c + nadp_c --> ac_c + nadph_c | h_c | -2.000000 |
| NanoG0094 | dkmpp_c + h2o_c --> 12dmpo_c + h_c + pi_c | dkmpp_c + h2o_c --> 12dmpo_c + pi_c | h_c | -1.000000 |
| ATAH_co2 | alltt_m + h2o_m + h_m --> co2_m + nh4_m + udg_m | alltt_m + h2o_m --> co2_m + nh4_m + udg_m | h_m | 1.000000 |

**Table S5C** Reactions restricted to one direction to remove energy-generating cycles that are present in the shipped model.

| **Reaction** | **Equation** | **Bounds before** | **Bounds in shipped model** |
| --- | --- | --- | --- |
| PPAth | ppa_c <-- ppa_h | [-1000, 1000] | [-1000, 0] |
| ACtm | ac_c <-- ac_m | [-1000, 1000] | [-1000, 0] |
| BCTA(val)m | 3mob_m + glu__L_m <-- akg_m + val__L_m | [-1000, 1000] | [-1000, 0] |
| GAPD | g3p_c + nad_c + pi_c --> 13dpg_c + nadh_c | [-1000, 1000] | [0, 1000] |
| UMPK4 | gtp_c + ump_c --> gdp_c + udp_c | [-1000, 1000] | [0, 1000] |
| 2PGtm | 2pg_c <-- 2pg_m | [-1000, 1000] | [-1000, 0] |
| CYTK13 | datp_c + dcmp_c --> dadp_c + dcdp_c | [-1000, 1000] | [0, 1000] |
| CYTK7 | cmp_c + utp_c <-- cdp_c + udp_c | [-1000, 1000] | [-1000, 0] |
| UMPK5 | datp_c + ump_c <-- dadp_c + udp_c | [-1000, 1000] | [-1000, 0] |
| BCTA(val) | akg_c + val__L_c <-- 3mob_c + glu__L_c | [-1000, 1000] | [-1000, 0] |
| ASPNA1th | asp__L_c + na1_c --> asp__L_h + na1_h | [-1000, 1000] | [0, 1000] |
| DICtm | mal__L_c + succ_m <-- mal__L_m + succ_c | [-1000, 1000] | [-1000, 0] |
| HYDA | 2.0 fdxrd_c + 2.0 h_c <-- 2.0 fdxox_c + h2_c | [-1000, 1000] | [-1000, 0] |
| CY_focytc(h) | 2.0 ficytc_h + nadh_h <-- 2.0 focytc_h + nad_h | [-1000, 1000] | [-1000, 0] |

**Table S5D** Reactions left with a single direction in Stage 5 that are present in the shipped model.

| **Reaction** | **Equation** | **Bounds before** | **Bounds in shipped model** | **Gene association** |
| --- | --- | --- | --- | --- |
| 1ACPCth | 1acpc_c --> 1acpc_h | [-1000, 1000] | [0, 1000] | No |
| 4ABUTth | 4abut_c --> 4abut_h | [-1000, 1000] | [0, 1000] | No |
| 6PGLthr | 6pgl_c --> 6pgl_h | [-1000, 1000] | [0, 1000] | No |
| CMPth | cmp_c <-- cmp_h | [-1000, 1000] | [-1000, 0] | No |
| COAth | coa_c <-- coa_h | [-1000, 1000] | [-1000, 0] | No |
| CTPth | ctp_c --> ctp_h | [-1000, 1000] | [0, 1000] | No |
| DMPPth | dmpp_h --> dmpp_c | [-1000, 1000] | [0, 1000] | No |
| G1P(pi)th | g1p_c + pi_h --> g1p_h + pi_c | [-1000, 1000] | [0, 1000] | No |
| G6PA(pi)th | g6p-A_c + pi_h --> g6p-A_h + pi_c | [-1000, 1000] | [0, 1000] | No |
| GLNth | gln__L_c + h_c --> gln__L_h + h_h | [-1000, 1000] | [0, 1000] | Yes |
| GLYNA1th | gly_c + na1_c --> gly_h + na1_h | [-1000, 1000] | [0, 1000] | Yes |
| GLYth | gly_c + h_c --> gly_h + h_h | [-1000, 1000] | [0, 1000] | Yes |
| IPDPth | ipdp_h --> ipdp_c | [-1000, 1000] | [0, 1000] | No |
| MANth | man_c <-- man_h | [-1000, 1000] | [-1000, 0] | No |
| PMTCOAth | pmtcoa_c <=> pmtcoa_h | [-1000, 1000] | [-1000, 1000] | No |
| PPCOAth | ppcoa_c <-- ppcoa_h | [-1000, 1000] | [-1000, 0] | No |
| SERNA1th | na1_c + ser__L_c --> na1_h + ser__L_h | [-1000, 1000] | [0, 1000] | Yes |
| SERth | h_c + ser__L_c <-- h_h + ser__L_h | [-1000, 1000] | [-1000, 0] | Yes |
| THRNA1th | na1_c + thr__L_c --> na1_h + thr__L_h | [-1000, 1000] | [0, 1000] | Yes |
| THRth | h_c + thr__L_c <-- h_h + thr__L_h | [-1000, 1000] | [-1000, 0] | Yes |
| UDPGALth | udpgal_c + ump_h --> udpgal_h + ump_c | [-1000, 1000] | [0, 1000] | Yes |
| UDPth | udp_c <-- udp_h | [-1000, 1000] | [-1000, 0] | No |
| UMPth | ump_c --> ump_h | [-1000, 1000] | [0, 1000] | No |
| VALth | h_c + val__L_c <=> h_h + val__L_h | [-1000, 1000] | [-1000, 1000] | Yes |
| SO4t | h_e + so4_e --> h_c + so4_c | [-1000, 1000] | [0, 1000] | Yes |
| H2Otx | h2o_c --> h2o_x | [-1000, 1000] | [0, 1000] | No |
| NH4tx | nh4_c --> nh4_x | [-1000, 1000] | [0, 1000] | Yes |
| 10FTHFtx | 10fthf_c --> 10fthf_x | [-1000, 1000] | [0, 1000] | No |
| METHFtx | methf_c --> methf_x | [-1000, 1000] | [0, 1000] | Yes |
| 1ACPCtm | 1acpc_c --> 1acpc_m | [-1000, 1000] | [0, 1000] | No |
| 2DHPtm | 2dhp_c --> 2dhp_m | [-1000, 1000] | [0, 1000] | No |
| 4ABUTtm | 4abut_c --> 4abut_m | [-1000, 1000] | [0, 1000] | No |
| ASNNA1tm | asn__L_c + na1_c --> asn__L_m + na1_m | [-1000, 1000] | [0, 1000] | Yes |
| ASNtm | asn__L_c + h_c --> asn__L_m + h_m | [-1000, 1000] | [0, 1000] | Yes |
| ASPNA1tm | asp__L_c + na1_c --> asp__L_m + na1_m | [-1000, 1000] | [0, 1000] | No |
| ASPtm | asp__L_c + h_c --> asp__L_m + h_m | [-1000, 1000] | [0, 1000] | Yes |
| GLYNA1tm | gly_c + na1_c <-- gly_m + na1_m | [-1000, 1000] | [-1000, 0] | Yes |
| GLYtm | gly_c + h_c --> gly_m + h_m | [-1000, 1000] | [0, 1000] | Yes |
| H2tm | h2_c <-- h2_m | [-1000, 1000] | [-1000, 0] | No |
| METHFtm | h_c + methf_c --> h_m + methf_m | [-1000, 1000] | [0, 1000] | Yes |
| MLTHFtm | h_c + mlthf_c <-- h_m + mlthf_m | [-1000, 1000] | [-1000, 0] | Yes |
| NADtm | nad_c --> nad_m | [-1000, 1000] | [0, 1000] | No |
| PRONA1tm | na1_c + pro__L_c --> na1_m + pro__L_m | [-1000, 1000] | [0, 1000] | Yes |
| PROtm | h_c + pro__L_c --> h_m + pro__L_m | [-1000, 1000] | [0, 1000] | Yes |
| SERNA1tm | na1_c + ser__L_c --> na1_m + ser__L_m | [-1000, 1000] | [0, 1000] | Yes |
| SERtm | h_c + ser__L_c --> h_m + ser__L_m | [-1000, 1000] | [0, 1000] | Yes |
| SPMDtmr | h_m + spmd_c --> h_c + spmd_m | [-1000, 1000] | [0, 1000] | No |
| SUCFUMtmr | fum_m + succ_c <-- fum_c + succ_m | [-1000, 1000] | [-1000, 0] | Yes |
| THFtm | h_c + thf_c --> h_m + thf_m | [-1000, 1000] | [0, 1000] | Yes |
| THRNA1tm | na1_c + thr__L_c --> na1_m + thr__L_m | [-1000, 1000] | [0, 1000] | Yes |
| THRtm | h_c + thr__L_c --> h_m + thr__L_m | [-1000, 1000] | [0, 1000] | Yes |
| VALtm | h_c + val__L_c --> h_m + val__L_m | [-1000, 1000] | [0, 1000] | Yes |
| CAROtu | caro_u --> caro_h | [-1000, 1000] | [0, 1000] | No |
| GCAROtu | gcaro_h --> gcaro_u | [-1000, 1000] | [0, 1000] | No |
| ASPATh | akg_h + asp__L_h <-- glu__L_h + oaa_h | [-1000, 1000] | [-1000, 0] | Yes |
| G6PDA | gam6p_c + h2o_c --> f6p-B_c + nh4_c | [-1000, 1000] | [0, 1000] | Yes |
| UA4E | udpxyl_c --> udparab_c | [-1000, 1000] | [0, 1000] | Yes |
| UDPACGLP | acgam1p_c + utp_c --> ppi_c + uacgam_c | [-1000, 1000] | [0, 1000] | Yes |
| AOTA | acg5sa_c + glu__L_c --> acorn_c + akg_c | [-1000, 1000] | [0, 1000] | Yes |
| ORNTArm | akg_m + orn_m --> glu5sa_m + glu__L_m | [-1000, 1000] | [0, 1000] | Yes |
| CYARTMT | amet_c + cyart_c --> ahcys_c + menecyart_c | [-1000, 1000] | [0, 1000] | Yes |
| DSMSTOLD | ddsmsterol_c + fadh2_c --> dsmsterol_c + fad_c | [-1000, 1000] | [0, 1000] | No |
| LNS14DM | lanost_c + 3.0 nadph_c + 3.0 o2_c --> 44mctr_c + for_c + 4.0 h2o_c + 3.0 nadp_c | [-1000, 1000] | [0, 1000] | Yes |
| ALAAT | akg_c + ala__L_c --> glu__L_c + pyr_c | [-1000, 1000] | [0, 1000] | Yes |
| HCO3E | hco3_c --> co2_c + h2o_c | [-1000, 1000] | [0, 1000] | Yes |
| HCO3Em | hco3_m <-- co2_m + h2o_m | [-1000, 1000] | [-1000, 0] | Yes |
| PRUK | atp_h + h_h + ru5p-D_h --> adp_h + rb15bp_h | [-1000, 1000] | [0, 1000] | Yes |
| RBPCh | co2_h + h2o_h + rb15bp_h --> 2.0 3pg_h + h_h | [-1000, 1000] | [0, 1000] | Yes |
| SBP | h2o_h + s17bp_h --> pi_h + s7p_h | [-1000, 1000] | [0, 1000] | Yes |
| ASPATm | akg_m + asp__L_m --> glu__L_m + oaa_m | [-1000, 1000] | [0, 1000] | Yes |
| LCYB | gcaro_u --> caro_u | [-1000, 1000] | [0, 1000] | Yes |
| LCYG | lyc_h --> gcaro_h | [-1000, 1000] | [0, 1000] | Yes |
| 3HAD40 | 3hbutACP_h --> but2eACP_h + h2o_h | [-1000, 1000] | [0, 1000] | Yes |
| 3HAD60 | 3hhexACP_h --> h2o_h + thex2eACP_h | [-1000, 1000] | [0, 1000] | Yes |
| 3HAD80 | 3hoctACP_h --> h2o_h + toct2eACP_h | [-1000, 1000] | [0, 1000] | Yes |
| ECOAH3m | 3hocoa_m --> h2o_m + oc2coa_m | [-1000, 1000] | [0, 1000] | No |
| ECOAH5m | 3hddcoa_m --> dd2coa_m + h2o_m | [-1000, 1000] | [0, 1000] | No |
| ECOAH6m | 3htdcoa_m --> h2o_m + td2coa_m | [-1000, 1000] | [0, 1000] | No |
| HACD2m | 3ohcoa_m + nadh_m --> 3hhcoa_m + nad_m | [-1000, 1000] | [0, 1000] | Yes |
| FPGPL | f1p_h --> dhap_h + glyald_h | [-1000, 1000] | [0, 1000] | Yes |
| MAKI | man_h <-- fru-B_h | [-1000, 1000] | [-1000, 0] | No |
| SBTRh | nadp_h + sbt-D_h --> glc-A_h + nadph_h | [-1000, 1000] | [0, 1000] | Yes |
| UG4E | udpg_c --> udpgal_c | [-1000, 1000] | [0, 1000] | Yes |
| GLYALDOR | glyc-R_c + nadh_c --> glyald_c + h2o_c + nad_c | [-1000, 1000] | [0, 1000] | Yes |
| G3PO | dhap_h + h2o2_h --> glyc3p_h + o2_h | [-1000, 1000] | [0, 1000] | No |
| SPTA | pyr_c + ser__L_c --> ala__L_c + hpyr_c | [-1000, 1000] | [0, 1000] | Yes |
| THRAL | ser__L_h <-- nh4_h + pyr_h | [-1000, 1000] | [-1000, 0] | Yes |
| A1E | glc-A_c --> glc-B_c | [-1000, 1000] | [0, 1000] | Yes |
| G6PI | g6p-A_c --> g6p-B_c | [-1000, 1000] | [0, 1000] | Yes |
| PGCM | g1p_c <-- g6p-A_c | [-1000, 1000] | [-1000, 0] | Yes |
| PGIA | g6p-A_c <-- f6p-B_c | [-1000, 1000] | [-1000, 0] | Yes |
| PGMTh | g1p_h <-- g6p-A_h | [-1000, 1000] | [-1000, 0] | Yes |
| PDHe2hr | adhlam_h + coa_h --> accoa_h + dhlam_h | [-1000, 1000] | [0, 1000] | Yes |
| PDHe2r | adhlam_m + coa_m --> accoa_m + dhlam_m | [-1000, 1000] | [0, 1000] | Yes |
| PDHe3hr | dhlam_h + nad_h --> lpam_h + nadh_h | [-1000, 1000] | [0, 1000] | No |
| PGKh | 13dpg_h + adp_h <-- 3pg_h + atp_h | [-1000, 1000] | [-1000, 0] | Yes |
| TPIh | dhap_h <-- g3p_h | [-1000, 1000] | [-1000, 0] | Yes |
| RBCh | o2_h + rb15bp_h --> 2pglyc_h + 3pg_h + h_h | [-1000, 1000] | [0, 1000] | No |
| MDUMT | dump_c + mlthf_c --> dhf_c + dtmp_c | [-1000, 1000] | [0, 1000] | Yes |
| APPAT | atp_c + pan4p_c --> dpcoa_c + ppi_c | [-1000, 1000] | [0, 1000] | Yes |
| PPM | r1p_c --> r5p_c | [-1000, 1000] | [0, 1000] | Yes |
| TAh | g3p_h + s7p_h --> e4p_h + f6p-B_h | [-1000, 1000] | [0, 1000] | Yes |
| TYRTAh | 34hpp_h + glu__L_h --> akg_h + tyr__L_h | [-1000, 1000] | [0, 1000] | Yes |
| ATPSh | adp_h + 4.0 h_u + pi_h --> atp_h + h2o_h + 4.0 h_h | [-1000, 1000] | [0, 1000] | Yes |
| ACCAHh | 1acpc_h + h2o_h --> 2obut_h + nh4_h | [-1000, 1000] | [0, 1000] | No |
| ACCAHm | 1acpc_m + h2o_m --> 2obut_m + nh4_m | [-1000, 1000] | [0, 1000] | No |
| APPRT | ade_c + prpp_c --> amp_c + ppi_c | [-1000, 1000] | [0, 1000] | Yes |
| GBDP | h2o_c + ppgpp_c <-- gdp_c + ppi_c | [-1000, 1000] | [-1000, 0] | Yes |
| GDTP | gdptp_c + h2o_c <-- pi_c + ppgpp_c | [-1000, 1000] | [-1000, 0] | Yes |
| GPAR | gua_c + prpp_c --> gmp_c + ppi_c | [-1000, 1000] | [0, 1000] | Yes |
| GTPDPK | atp_c + gtp_c <-- amp_c + gdptp_c | [-1000, 1000] | [-1000, 0] | Yes |
| HPRT | hxan_c + prpp_c --> imp_c + ppi_c | [-1000, 1000] | [0, 1000] | Yes |
| XPPRT | prpp_c + xan_c --> ppi_c + xmp_c | [-1000, 1000] | [0, 1000] | Yes |
| ACKrm | ac_m + atp_m --> actp_m + adp_m | [-1000, 1000] | [0, 1000] | Yes |
| ASPAT | akg_c + asp__L_c --> glu__L_c + oaa_c | [-1000, 1000] | [0, 1000] | Yes |
| PTArm | accoa_m + pi_m --> actp_m + coa_m | [-1000, 1000] | [0, 1000] | Yes |
| HYDAm | 2.0 fdxrd_m + 2.0 h_m --> 2.0 fdxox_m + h2_m | [-1000, 1000] | [0, 1000] | Yes |
| ACSERL | acser_c + seln_c --> ac_c + selcys_c | [-1000, 1000] | [0, 1000] | Yes |
| MDHm | nadh_m + oaa_m --> mal__L_m + nad_m | [-1000, 1000] | [0, 1000] | Yes |
| FUMm | fum_m + h2o_m --> mal__L_m | [-1000, 1000] | [0, 1000] | Yes |
| ICDHhr | icit_h + nadp_h --> akg_h + co2_h + nadph_h | [-1000, 1000] | [0, 1000] | No |
| IMDHT(3c2hmp) | 2ippm_c + h2o_c --> 3c2hmp_c | [-1000, 1000] | [0, 1000] | No |
| HMNO | 3hib_m + nad_m --> mmsa_m + nadh_m | [-1000, 1000] | [0, 1000] | Yes |
| PYDXOR | nadph_c + pydx_c --> nadp_c + pydxn_c | [-1000, 1000] | [0, 1000] | Yes |
| upCv | pi_c + uri_c --> r1p_c + ura_c | [-1000, 1000] | [0, 1000] | No |
| CRETINOLOR1 | nadh_h + retinal-11-cis_h --> nad_h + retinol-cis-11_h | [-1000, 1000] | [0, 1000] | No |
| TRETINOLOR1 | nadh_h + retinal_h --> nad_h + retinol_h | [-1000, 1000] | [0, 1000] | No |
| GLCt2h | glc-A_c + h_c --> glc-A_h + h_h | [-1000, 1000] | [0, 1000] | No |
| citt | cit_e --> cit_c | [-1000, 1000] | [0, 1000] | No |
| urat | ura_e --> ura_c | [-1000, 1000] | [0, 1000] | No |
| ARAI | arab__L_c --> rbl__L_c | [-1000, 1000] | [0, 1000] | No |
| RBP4E | ru5p__L_c --> xu5p-D_c | [-1000, 1000] | [0, 1000] | No |
| RPE | xu5p-D_c --> ru5p-D_c | [-1000, 1000] | [0, 1000] | Yes |
| Ru5PPi_th | pi_h + ru5p-D_c --> pi_c + ru5p-D_h | [-1000, 1000] | [0, 1000] | No |
| LACDt_c | lac-D_e --> lac-D_c | [-1000, 1000] | [0, 1000] | No |
| UDPGLCter | udpg_c + ump_r --> udpg_r + ump_c | [-1000, 1000] | [0, 1000] | No |
| UMPtC | ump_r --> ump_c | [-1000, 1000] | [0, 1000] | No |
| SSALxm | h2o_m + nad_m + sucsal_m --> nadh_m + succ_m | [-1000, 1000] | [0, 1000] | No |
| MTHFC | h2o_c + methf_c --> 10fthf_c | [-1000, 1000] | [0, 1000] | No |
| DURAD | 56dura_c + nadp_c --> nadph_c + ura_c | [-1000, 1000] | [0, 1000] | No |
| CYTK12n | dcmp_n + dctp_n --> 2.0 dcdp_n | [-1000, 1000] | [0, 1000] | No |
| CYTK13n | datp_n + dcmp_n --> dadp_n + dcdp_n | [-1000, 1000] | [0, 1000] | No |
| CYTK2n | atp_n + dcmp_n --> adp_n + dcdp_n | [-1000, 1000] | [0, 1000] | No |
| CYTK6n | cmp_n + ctp_n --> 2.0 cdp_n | [-1000, 1000] | [0, 1000] | No |
| CYTK7n | cmp_n + utp_n --> cdp_n + udp_n | [-1000, 1000] | [0, 1000] | No |
| CYTK8n | cmp_n + datp_n --> cdp_n + dadp_n | [-1000, 1000] | [0, 1000] | No |
| UMPK4n | gtp_n + ump_n --> gdp_n + udp_n | [-1000, 1000] | [0, 1000] | No |
| UMPK6n | dctp_n + ump_n --> dcdp_n + udp_n | [-1000, 1000] | [0, 1000] | No |
| UMPK7n | dgtp_n + ump_n --> dgdp_n + udp_n | [-1000, 1000] | [0, 1000] | No |
| UMPKn | atp_n + ump_n --> adp_n + udp_n | [-1000, 1000] | [0, 1000] | No |
| CYTK10n | cmp_n + dgtp_n --> cdp_n + dgdp_n | [-1000, 1000] | [0, 1000] | No |
| CYTK11n | dcmp_n + dgtp_n --> dcdp_n + dgdp_n | [-1000, 1000] | [0, 1000] | No |
| CYTK14n | dcmp_n + utp_n --> dcdp_n + udp_n | [-1000, 1000] | [0, 1000] | No |
| CYTK1n | atp_n + cmp_n --> adp_n + cdp_n | [-1000, 1000] | [0, 1000] | No |
| CYTK3n | ctp_n + dcmp_n --> cdp_n + dcdp_n | [-1000, 1000] | [0, 1000] | No |
| CYTK4n | dcmp_n + gtp_n --> dcdp_n + gdp_n | [-1000, 1000] | [0, 1000] | No |
| CYTK9n | cmp_n + dctp_n --> cdp_n + dcdp_n | [-1000, 1000] | [0, 1000] | No |
| UMPK2n | ctp_n + ump_n --> cdp_n + udp_n | [-1000, 1000] | [0, 1000] | No |
| UMPK3n | ump_n + utp_n --> 2.0 udp_n | [-1000, 1000] | [0, 1000] | No |
| UMPK5n | datp_n + ump_n --> dadp_n + udp_n | [-1000, 1000] | [0, 1000] | No |
| CITt4_4 | cit_e + 4.0 na1_e --> cit_c + 4.0 na1_c | [-1000, 1000] | [0, 1000] | No |
| CITt4_2 | cit_e + 2.0 na1_e --> cit_c + 2.0 na1_c | [-1000, 1000] | [0, 1000] | No |
| pmtcoa_TRANSFERASE3_x | pmtcoa_c --> pmtcoa_x | [-1000, 1000] | [0, 1000] | No |
| BCTA(val)h | akg_h + val__L_h <=> 3mob_h + glu__L_h | [-1000, 1000] | [-1000, 1000] | No |
| IVCDH | fad_m + ivcoa_m --> 3mb2coa_m + fadh2_m | [-1000, 1000] | [0, 1000] | Yes |
| MCDH(2mb2coa) | 2mbcoa_m + fad_m --> 2mb2coa_m + fadh2_m | [-1000, 1000] | [0, 1000] | No |
| EAR100x | nadh_h + tdec2eACP_h --> dcaACP_h + nad_h | [-1000, 1000] | [0, 1000] | Yes |
| EAR100y | nadph_h + tdec2eACP_h --> dcaACP_h + nadp_h | [-1000, 1000] | [0, 1000] | Yes |
| EAR120x | nadh_h + tddec2eACP_h --> ddcaACP_h + nad_h | [-1000, 1000] | [0, 1000] | Yes |
| EAR120y | nadph_h + tddec2eACP_h --> ddcaACP_h + nadp_h | [-1000, 1000] | [0, 1000] | Yes |
| EAR140x | nadh_h + tmrs2eACP_h --> myrsACP_h + nad_h | [-1000, 1000] | [0, 1000] | Yes |
| EAR140y | nadph_h + tmrs2eACP_h --> myrsACP_h + nadp_h | [-1000, 1000] | [0, 1000] | Yes |
| EAR160x | nadh_h + tpalm2eACP_h --> nad_h + palmACP_h | [-1000, 1000] | [0, 1000] | Yes |
| EAR160y | nadph_h + tpalm2eACP_h --> nadp_h + palmACP_h | [-1000, 1000] | [0, 1000] | Yes |
| EAR181x | nadh_h + t3c11vaceACP_h --> nad_h + octeACP_h | [-1000, 1000] | [0, 1000] | Yes |
| EAR181y | nadph_h + t3c11vaceACP_h --> nadp_h + octeACP_h | [-1000, 1000] | [0, 1000] | Yes |
| EAR40x | but2eACP_h + nadh_h --> butACP_h + nad_h | [-1000, 1000] | [0, 1000] | Yes |
| EAR40y | but2eACP_h + nadph_h --> butACP_h + nadp_h | [-1000, 1000] | [0, 1000] | Yes |
| EAR60x | nadh_h + thex2eACP_h --> hexACP_h + nad_h | [-1000, 1000] | [0, 1000] | Yes |
| EAR60y | nadph_h + thex2eACP_h --> hexACP_h + nadp_h | [-1000, 1000] | [0, 1000] | Yes |
| EAR80x | nadh_h + toct2eACP_h --> nad_h + ocACP_h | [-1000, 1000] | [0, 1000] | Yes |
| EAR80y | nadph_h + toct2eACP_h --> nadp_h + ocACP_h | [-1000, 1000] | [0, 1000] | Yes |
| 3MOBtm | 3mob_c <-- 3mob_m | [-1000, 1000] | [-1000, 0] | No |

**Table S5E** Reactions that remain elementally imbalanced in the shipped model.

| **Reaction** | **Name** | **Category** | **Imbalance** | **Equation** |
| --- | --- | --- | --- | --- |
| AC270_OXIDATION | AC270_OXIDATION | Ceramide and fatty acid formula mismatch | C +20, H +40 | atp_h + coa_h + heptac_h --> amp_h + heptcoscoa_h + ppi_h |
| CER160_160_SYN | CER160_160_SYN | Ceramide and fatty acid formula mismatch | C -2, H -3 | dhcrm160160_r + 2.0 h_r + nadph_r + o2_r --> crm160160_r + 2.0 h2o_r + nadp_r |
| CER170_250O_SYN | CER170_250O_SYN | Ceramide and fatty acid formula mismatch | C -1, H -1, O +1 | dhcrm170250(O)_r + 2.0 h_r + nadph_r + o2_r --> crm170250(O)_r + 2.0 h2o_r + nadp_r |
| CER180_260O_SYN | CER180_260O_SYN | Ceramide and fatty acid formula mismatch | O +1 | dhcrm180260(O)_r + 2.0 h_r + nadph_r + o2_r --> crm180260(O)_r + 2.0 h2o_r + nadp_r |
| CER200_160_SYN | CER200_160_SYN | Ceramide and fatty acid formula mismatch | C +2, H +5 | dhcrm200160_r + 2.0 h_r + nadph_r + o2_r --> crm200160_r + 2.0 h2o_r + nadp_r |
| CER200_220O_SYN | CER200_220O_SYN | Ceramide and fatty acid formula mismatch | C +2, H +1, O +1 | dhcrm200220(O)_r + 6.0 h_r + nadph_r + o2_r --> crm200220(O)_r + 2.0 h2o_r + nadp_r |
| CER2219Z_1819Z2O_SYN | CER2219Z_1819Z2O_SYN | Ceramide and fatty acid formula mismatch | H -1, O +1 | dhcrm2219Z1819Z(2O)_r + 2.0 h_r + nadph_r + o2_r --> crm2219Z1819Z(2O)_r + 2.0 h2o_r + nadp_r |
| FA230(2OH)ACPHi | FA230(2OH)ACPHi | Ceramide and fatty acid formula mismatch | O -1 | h2o_h + tricosaACP(2OH)_h --> ACP_h + tricosac(2OH)_h |
| FA230(2OH)ACPHiC | FA230(2OH)ACPHiC | Ceramide and fatty acid formula mismatch | O -1 | h2o_c + tricosaACP(2OH)_c --> ACP_c + tricosac(2OH)_c |
| FACOAL230(2OH)_ACP | FACOAL230(2OH)_ACP | Ceramide and fatty acid formula mismatch | O +1 | 4.0 h_h + henecosaACP_h + malACP_h --> ACP_h + co2_h + h2o_h + tricosaACP(2OH)_h |
| FACOAL230(2OH)_ACPc | FACOAL230(2OH)_ACPc | Ceramide and fatty acid formula mismatch | O +1 | 4.0 h_c + henecosaACP_c + malACP_c --> ACP_c + co2_c + h2o_c + tricosaACP(2OH)_c |
| FACOAL40_ACP | FACOAL40_ACP | Ceramide and fatty acid formula mismatch | C -1, H -2, O +1 | actACP_h + 4.0 h_h + malACP_h --> ACP_h + butACP_h + 2.0 co2_h + h2o_c |
| NDP7er | NDP7er | ER proton (NDP7er) | H +1 | h2o_r + udp_r --> h_r + pi_r + ump_r |
| METARHODOPSINH | metarhodopsin_hydrolase_(spontaneous) | R-group placeholder | R -1 | metarhodopsin_h --> retinal_h |
| PS_asn-L(c) | incorporation_of_asn-L_residue_into_a_protein | R-group placeholder | C +1, H +1, N +1, O +1, R +2 | asn__L_c --> Asn-X-Ser-FSLASH-Thr_c + h2o_c |
| RHODOPSINRT | rhodopsin_retinyltransferase_(spontaneous) | R-group placeholder | R +1 | retinal-11-cis_h --> rhodopsin_h |
| DVPCHLD450OR | divinylprotochlorophyllide_oxidoreductase_(light-dependent,_450_nm) | Z placeholder (photons) | Z -1 | dvpchlda_h + nadph_h + photon450_h --> dvchlda_h + nadp_h |
| DVPCHLD646OR | divinylprotochlorophyllide_oxidoreductase_(light-dependent,_648_nm) | Z placeholder (photons) | Z -1 | dvpchlda_h + nadph_h + photon646_h --> dvchlda_h + nadp_h |
| PCHLD450OR | protochlorophyllide_oxidoreductase_(light-dependent,_450_nm) | Z placeholder (photons) | H -1, Z -1 | h_h + nadph_h + pchlda_h + photon450_h --> chlda_h + nadp_h |
| PCHLD646OR | protochlorophyllide_oxidoreductase_(light-dependent,_648_nm) | Z placeholder (photons) | H -1, Z -1 | h_h + nadph_h + pchlda_h + photon646_h --> chlda_h + nadp_h |
| PRISM_design_growth | spectral_decomposition_of_an_optimized_LED_for_growth | Z placeholder (photons) | Z +0.59276 | photonVis_e --> 0.220793969934617 photon646_h + 0.638343344969578 photon673_u + 0.73362239147121 photon680_u |
| PRISM_fluorescent_cool_215W | spectral_decomposition_of_215W_cool_white_fluorescent_light_bulb | Z placeholder (photons) | Z -0.27533 | photonVis_e --> 9.86915907870449e-05 photon298_c + 0.097915549659156 photon437_u + 0.150739641718055 photon438_u + 0.107796806656087 photon450_h + 0.11138 photon490_h + 0.1946593756116 photon646_h + 0.030722582685521 photon673_u + 0.031357804045244 photon680_u |
| PRISM_fluorescent_warm_18W | spectral_decomposition_of_18W_warm_white_fluorescent_light_bulb | Z placeholder (photons) | Z -0.0983939 | photonVis_e --> 0.022608265086577 photon298_c + 0.152956409671983 photon437_u + 0.231224055045519 photon438_u + 0.152106146847886 photon450_h + 0.11038 photon490_h + 0.159331959324159 photon646_h + 0.035127278057213 photon673_u + 0.03787199644559 photon680_u |
| PRISM_growth_room | spectral_decomposition_of_growth_room | Z placeholder (photons) | Z -0.30534 | photonVis_e --> 4.11265742135554e-05 photon298_c + 0.075412270199012 photon437_u + 0.117405566728347 photon438_u + 0.083033179648876 photon450_h + 0.094128 photon490_h + 0.20332960252981 photon646_h + 0.05700259056653 photon673_u + 0.064307747854245 photon680_u |
| PRISM_high_pressure_sodium | spectral_decomposition_of_high_pressure_sodium_lamp | Z placeholder (photons) | Z -0.518038 | photonVis_e --> 0.015196534268239 photon437_u + 0.032805456604757 photon438_u + 0.027719995714444 photon450_h + 0.058194 photon490_h + 0.242892771920299 photon646_h + 0.051456782792204 photon673_u + 0.053696898066714 photon680_u |
| PRISM_incandescent_60W | spectral_decomposition_of_60W_incandescent_light_bulb | Z placeholder (photons) | Z -0.385049 | photonVis_e --> 1.65224841134181e-05 photon298_c + 0.009993878664639 photon437_u + 0.024602315301096 photon438_u + 0.016276171066119 photon450_h + 0.053691 photon490_h + 0.22991250033838 photon646_h + 0.12754084169748 photon673_u + 0.152917317912673 photon680_u |
| PRISM_metal_halide | spectral_decomposition_of_metal_halide_lamp | Z placeholder (photons) | Z -0.229873 | photonVis_e --> 0.12093046741464 photon437_u + 0.191654658116142 photon438_u + 0.134003914972887 photon450_h + 0.119084 photon490_h + 0.119049343117245 photon646_h + 0.040192911402728 photon673_u + 0.045211338768487 photon680_u |
| PRISM_red_LED_674nm | spectral_decomposition_of_red_LED_(674nm) | Z placeholder (photons) | Z +0.651196 | photonVis_e --> 0.265727748400781 photon646_h + 0.660147062166474 photon673_u + 0.725321425353414 photon680_u |
| PRISM_red_LED_array_653nm | spectral_decomposition_of_red_LED_array_(653nm) | Z placeholder (photons) | Z +0.340111 | photonVis_e --> 0.870291149591763 photon646_h + 0.274325356868054 photon673_u + 0.195494060752262 photon680_u |
| PRISM_solar_exo | spectral_decomposition_of_solar_light_measured_from_outer_space | Z placeholder (photons) | Z +0.0124798 | photonVis_e --> 0.009126633453201 photon298_c + 0.107054407672067 photon437_u + 0.224061663093904 photon438_u + 0.133591526308582 photon450_h + 0.20968 photon490_h + 0.17175215519119 photon646_h + 0.072897947189877 photon673_u + 0.084315507784363 photon680_u |
| PRISM_solar_litho | spectral_decomposition_of_solar_light_measured_from_Earth's_surface | Z placeholder (photons) | Z -0.0498595 | photonVis_e --> 4.26984325300204e-05 photon298_c + 0.088253056987783 photon437_u + 0.187390761056649 photon438_u + 0.114958292879089 photon450_h + 0.19581 photon490_h + 0.18566429641716 photon646_h + 0.08361649412414 photon673_u + 0.094404940130699 photon680_u |
| PRISM_white_LED | spectral_decomposition_of_white_LED | Z placeholder (photons) | Z -0.365097 | photonVis_e --> 0.000222203715307 photon298_c + 0.007117672622224 photon437_u + 0.045915569014989 photon438_u + 0.027352098019382 photon450_h + 0.15185 photon490_h + 0.25680923332143 photon646_h + 0.07032130315189 photon673_u + 0.075315322428205 photon680_u |
| PSIIblue | photosystem_II_(blue_light-activated) | Z placeholder (photons) | Z -4 | 2.0 h2o_u + 4.0 photon438_u + 2.0 pq_u --> o2D_u + 2.0 pqh2_u |
| PSIIred | photosystem_II_(red_light-activated) | Z placeholder (photons) | Z -4 | 2.0 h2o_u + 4.0 photon673_u + 2.0 pq_u --> o2D_u + 2.0 pqh2_u |
| PSIblue | photosystem_I_(blue_light-activated) | Z placeholder (photons) | Z -2 | fdxox_u + 2.0 pccu1p_u + 2.0 photon437_u --> fdxrd_u + 2.0 pccu2p_u |
| PSIred | photosystem_I_(red_light-activated) | Z placeholder (photons) | Z -2 | fdxox_u + 2.0 pccu1p_u + 2.0 photon680_u --> fdxrd_u + 2.0 pccu2p_u |
| RHODOPSINI | rhodopsin_photoisomerase_(spontaneous,_light-requiring) | Z placeholder (photons) | Z -1.6 | 1.6 photon490_h + rhodopsin_h --> bathorhodopsin_h |
| VITD3S | vitamin_D3_synthesis_(non-enzymatic,_UV-light_requiring) | Z placeholder (photons) | Z -1 | 7dhchsterol_c + photon298_c --> vitd3_c |
| STARCH300DEGRA | degradation_of_starch300_by_phosphorylase,_amylase,_dextrinase,_maltase_(chloroplast),_1:3_glc-A:g1p_(light-dep_rxn) | Hydrogen imbalance (starch degradation) | H +300 | 74.0 h2o_h + 225.0 pi_h + starch300_h --> 225.0 g1p_h + 75.0 glc-A_h |
| STARCH300DEGRB | degradation_of_starch300_by_phosphorylase,_amylase,_dextrinase,_maltase_(chloroplast),_1:3_glc-B:g1p_(light-dep_rxn) | Hydrogen imbalance (starch degradation) | H +300 | 74.0 h2o_h + 225.0 pi_h + starch300_h --> 225.0 g1p_h + 75.0 glc-B_h |
| CBFC | cytochrome_b6/f_complex | Electron carrier hydrogen content | H -2 | 2.0 h_h + 2.0 pccu2p_u + pqh2_u --> 4.0 h_u + 2.0 pccu1p_u + pq_u |
| FNORh | ferredoxin---NADP+_reductase | Electron carrier hydrogen content | H +3 | fdxrd_u + nadp_h --> fdxox_u + h_h + nadph_h |

**Table S5F** Metabolite formula corrections (Stage 6) present in the shipped model.

| **Reaction** | **Metabolite** | **Formula before** | **Formula in shipped model** |
| --- | --- | --- | --- |
| DHCER160_160 | dhcrm160160_r | C32H65NO3 | C34H68NO3 |
| DHCER170_250O | dhcrm170250(O)_r | C42H85NO4 | C43H86NO3 |
| DHCER200_160 | dhcrm200160_r | C36H73NO3 | C34H68NO3 |
| DHCER200_220O | dhcrm200220(O)_r | C42H81NO4 | C40H80NO3 |
| DHCER2219Z_1819Z2O | dhcrm2219Z1819Z(2O)_r | C40H77NO4 | C40H78NO3 |
| hexccoa(O)_TR | hexccoa(O)_r | C47H86N7O18P3S | C47H86N7O17P3S |
| PCHOL1402119Z_SYN | pchol1402119Z_c | C42H84NO8P | C43H83NO8P |
| PCHOL1419Z2119Z_SYN | pchol1419Z2119Z_c | C42H82NO8P | C43H81NO8P |
| PCHOL1719Z160_SYN | pchol1719Z160_c | C40H80NO8P | C41H79NO8P |
| PCHOL1719Z1719Z_SYN | pchol1719Z1719Z_c | C41H80NO8P | C42H79NO8P |
| PCHOL1719Z1819Z_SYN | pchol1719Z1819Z_c | C42H82NO8P | C43H81NO8P |
| PCHOL1719Z1829Z12Z_SYN | pchol1719Z1829Z12Z_c | C42H80NO8P | C43H79NO8P |
| PCHOL1719Z2039Z12Z15Z_SYN | pchol1719Z2039Z12Z15Z_c | C44H82NO8P | C45H81NO8P |
| PCHOL2019Z130_SYN | pchol2019Z130_c | C40H80NO8P | C41H79NO8P |
| PCHOL2119Z1023Z6Z_SYN | pchol2119Z1023Z6Z_c | C38H72NO8P | C39H71NO8P |
| PCHOL2119Z1119Z_SYN | pchol2119Z1119Z_c | C39H76NO8P | C40H75NO8P |
| PCHOL2119Z1216Z_SYN | pchol2119Z1216Z_c | C40H78NO8P | C41H77NO8P |
| PCHOL2119Z1219Z_SYN | pchol2119Z1219Z_c | C40H78NO8P | C41H77NO8P |
| PCHOL2119Z1226Z9Z_SYN | pchol2119Z1226Z9Z_c | C40H76NO8P | C41H75NO8P |
| pentacoscoa(O)_TR | pentacoscoa(O)_r | C46H89N7O18P3S | C46H84N7O17P3S |
| TAG1419Z1001133Z6Z9Z | tag1419Z1001133Z6Z9Z_h | C30H48O6 | C38H64O6 |
| TG1401001233Z6Z9Z | tag1401001233Z6Z9Z_h | C31H52O6 | C39H68O6 |
| TG1501016Z1123Z6Z | tag1501016Z1123Z6Z_h | C31H52O6 | C39H68O6 |
| TG1501116Z240 | tag1501116Z240_h | C45H84O6 | C53H100O6 |
| TG1501123Z6Z1226Z9Z | tag1501123Z6Z1226Z9Z_h | C33H54O6 | C41H70O6 |
| TG1601002429Z12Z | tag1601002429Z12Z_h | C45H86O6 | C53H98O6 |
| TG1839Z12Z15Z1123Z6Z2239Z12Z15Z | tag1839Z12Z15Z1123Z6Z2239Z12Z15Z_h | C55H90O6 | C54H88O6 |
| TG2046Z9Z12Z15Z1123Z6Z2319Z | tag2046Z9Z12Z15Z1123Z6Z2319Z_h | C56H100O6 | C57H96O6 |
| TG2236Z9Z12Z1123Z6Z2119Z | tag2236Z9Z12Z1123Z6Z2119Z_h | C48H84O6 | C57H98O6 |
| TG2519Z1023Z6Z1123Z6Z | tag2519Z1023Z6Z1123Z6Z_h | C54H94O6 | C49H84O6 |
| TG2601426Z9Z1433Z6Z9Z | tag2601426Z9Z1433Z6Z9Z_h | C56H98O6 | C57H100O6 |
| TG2608090 | tag2608090_h | C48H90O6 | C46H88O6 |
| TG2801101426Z9Z | tag2801101426Z9Z_h | C57H106O6 | C56H104O6 |
| TG2819Z1123Z6Z150 | tag2819Z1123Z6Z150_h | C58H106O6 | C57H104O6 |
| DHCER170_250O | dhcrm170250(O)_r | C43H91NO4 | C43H86NO3 |
| DHCER180_260O | dhcrm180260(O)_r | C44H89NO4 | C44H89NO3 |
| STARCH300S | starch300_h | C12H22O11 | C1800H2702O1501 |

**Table S5G** Proton stoichiometry of the thylakoid electron transport and ATP synthesis reactions present in the shipped model. The equation before is from the preceding model version and the equation after is from the shipped model. The change is the difference in the net coefficient of each proton species (negative, more consumed or less produced).

| **Reaction** | **Equation before** | **Equation in shipped model** | **Proton species** | **Change (mol H)** |
| --- | --- | --- | --- | --- |
| ATPSh | adp_h + pi_h --> atp_h + h2o_u | adp_h + 4.0 h_u + pi_h --> atp_h + h2o_h + 4.0 h_h | h_u, h_h | -4.000000, 4.000000 |
| CBFC | 2.0 pccu2p_u + pqh2_u --> 4.0 h_u + 2.0 pccu1p_u + pq_u | 2.0 h_h + 2.0 pccu2p_u + pqh2_u --> 4.0 h_u + 2.0 pccu1p_u + pq_u | h_h | -2.000000 |
| CEF | fdxrd_u + h_h + pccu2p_u --> fdxox_u + 2.0 h_u + pccu1p_u | fdxrd_u + 2.0 h_h + 2.0 pccu2p_u --> fdxox_u + 4.0 h_u + 2.0 pccu1p_u | h_u, h_h | 2.000000, -1.000000 |
| FNORh | fdxrd_u + 2.0 h_h + nadp_h --> fdxox_u + nadph_h | fdxrd_u + nadp_h --> fdxox_u + h_h + nadph_h | h_h | 3.000000 |
| PSIblue | fdxox_u + 2.0 pccu1p_u + 2.0 photon437_u --> fdxrd_u + 2.0 pccu2p_u | fdxox_u + 2.0 h_u + 2.0 pccu1p_u + 2.0 photon437_u --> fdxrd_u + 2.0 pccu2p_u | h_u | -2.000000 |
| PSIred | fdxox_u + 2.0 pccu1p_u + 2.0 photon680_u --> fdxrd_u + 2.0 pccu2p_u | fdxox_u + 2.0 h_u + 2.0 pccu1p_u + 2.0 photon680_u --> fdxrd_u + 2.0 pccu2p_u | h_u | -2.000000 |
| Htu | h_h <=> h_u | (empty stoichiometry) | h_u, h_h | -1.000000, 1.000000 |
